# Functional genetics of rice *PISTILLATA* genes unravels new roles and targets in flowering time, female fertility and parthenocarpy

**DOI:** 10.1101/2023.08.05.552136

**Authors:** Mohamed Zamzam, Ritabrata Basak, Sharad Singh, Sandhan Prakash, Raghavaram Peesapati, Imtiyaz Khanday, Sara Simonini, Ueli Grossniklaus, Usha Vijayraghavan

## Abstract

Diversification of transcription factors and of their downstream targets can contribute to new organ morphologies. An example is rice lodicule, a small fleshy petal homolog that aids floret opening, thus facilitates pollination and fertility. To understand mechanisms underlying its specification, we investigated the developmental functions of the rice *PISTILLATA* (*PI*) paralogs, *OsMADS2* and *OsMADS4.* Null *osmads2* mutants reiterated OsMADS2 nonredundant lodicule specification roles and revealed new roles in flowering time and floral organ number and fate. Doubly perturbed *osmads2^d8/d8^ osmads4kd* florets had severe abnormalities, were female infertile, yet initiated parthenocarpy. Ubiquitous *OsMADS4* overexpression rescued *osmads2* abnormalities. Target genes whose regulation can contribute to OsMADS2 functions were discovered by its genome-wide binding analyses and transcriptome profiling. Several targets are implicated in lodicule and stamen development, floral organ number, cell wall, cell shape and osmotic homeostasis processes. Some targets relevant to lodicule development are cell division regulators (Cyclin D6, Cyclin P4-1-like), aquaporin (PIP1A), peptide transporter (PTR2), vascular development regulator (HOX1) and cell wall modulator (GH9B16). Their deregulation underscores the perturbed cell division, tissue differentiation patterns and physiology of the malformed *osmads2* and *osmads2^d8/d8^ osmads4kd* lodicules. Altogether, we reveal novel roles for the rice *PI* paralogs in flowering time, panicle exsertion and embryo sac differentiation, divulge gene targets for lodicule development and provide mechanistic insights on the functional diversification of rice *PI* paralogs.

## Introduction

Floral organs in angiosperms are arranged in four concentric whorls. Their identity is controlled largely by the combinatorial action of homeotic transcription factors of the MIKC-type MADS-domain family (Lohmann and Weigel, 2002; Krizek and Fletcher, 2005). The ABC and the expanded ABCDE models of floral organ patterning, predict how five classes of homeotic transcription factors: A, B, C, D and E can pattern floral organs on the determinate floral meristem (Coen and Meyerowitz, 1991; Favaro et al., 2003; Pinyopich et al., 2003; Krizek and Fletcher, 2005). Although not strictly conserved across angiosperms, this model provides a baseline for gauging functional conservation *versus* divergence. The ABCDE model indicates that the specification of eudicot second and third whorl floral organs, petals and stamens, requires a pair of related B-class genes: *APETALA3* (*AP3*) and *PISTILLATA* (*PI*) in *Arabidopsis thaliana*, and *DEFICIENS* (*DEF*) and *GLOBOSA* (*GLO*) in *Antirrhinum majus* (Coen and Meyerowitz, 1991). Not surprisingly, studies in cereal models demonstrated that B-class genes are also required for specifying the identity of second and third whorl floral organs, lodicules and stamens (Ambrose et al., 2000; Nagasawa et al., 2003; Prasad and Vijayraghavan, 2003; Yadav et al., 2007). These reports support the interpretation that lodicules, in cereals and grasses, are eudicot petal homologs and indicate shared elements in the genetic control of lodicule and petal development. However, the remarkable morphological and functional differences of these organs indicate differences in their developmental pathways.

Rice genome constitutes two *PI* paralogs: *OsMADS2* and *OsMADS4* (Chung et al., 1995). The phenotypes of lodicules and stamens in single and double knockdown plants for these *PI*-clade genes, *i.e.*, *osmads2dsRNAi, osmads4dsRNAi*, and *osmads2- and osmads4-dsRNAi*, suggested their complete redundancy for stamen specification and unequal contribution for lodicule development (Prasad and Vijayraghavan, 2003; Yao et al., 2008). Yet, mechanisms underlying their unequal roles in lodicules are unknown.

The identification of DNA binding sites and target genes of transcription factors bring insights into their molecular functions. A previous study of our group, based on transcriptomic changes triggered by *OsMADS2* RNA interference-mediated gene silencing (RNAi), as detected by 22,000 probe-based microarrays, in a broad pool of developing floral tissues, showed that 9% of OsMADS2 targets had annotated functions in diverse aspects of cell division control and suggested a diversification compared to *Arabidopsis* PI (Yadav et al., 2007). Here, we first generated null alleles of *OsMADS2* by gene editing, carried out a detailed phenotypic analysis, and performed developmentally staged panicle transcriptome profiling. To unravel the gene regulatory networks underlying the sub-functionalized activities of OsMADS2, this transcriptome analysis was coupled with a determination of its genome-wide DNA occupancy in developing florets. Furthermore, we generated *osmads2 osmads4kd* plants, doubly compromised for both rice *PI* paralogs, and employed genetic complementation assays to test their functional conservation. These data provide a deeper understanding of the extent of functional divergence between the rice *PI* paralogs. Moreover, we deciphered novel processes regulated by rice *PI* paralogs, such as flowering time, panicle exsertion and ovule development. Importantly, we delineate some gene targets that underlie lodicule development and physiology, and the nonredundant role of OsMADS2 in regulating flowering time. Our data emphasize the importance of investigating homeotic genes outside the core eudicot models to gain novel insights into their evolutionary conservation and diversification.

## Results

### OsMADS2 controls floral organ number and flowering time

While the *OsMADS2 dsRN*Ai lines (Prasad and Vijayraghavan, 2003) and mutant reported very recently by Wang and colleagues (2024a) had abnormal lodicules, normal stamen morphology and fertility, Yao and colleagues (2008) described low fertility attributed to poor anther dehiscence. This discrepancy highlights the necessity of in-depth study of developmental phenotypes in several mutant alleles. In this respect, we generated multiple loss-of-function alleles of *OsMADS2* by CRISPR/Cas9 gene editing (Supplementary Fig. S1, A-C). The *osmads2^i1^-dbd* allele with a single base pair (bp) insertion caused a frameshift within the MADS DNA binding domain (DBD), leading to a premature stop codon. The *osmads2^i1^*, *osmads2^d2^*, and *osmads2^d8^* alleles contained a 1 bp insertion, a 2 bp deletion, and an 8 bp deletion, respectively and created premature stop codons within the coiled-coil keratin-like domain (K domain) that is involved in oligomerization. Segregation analyses of different *osmads2* alleles in the T4 progeny showed no evidence of segregation distortion from the expected Mendelian ratios (Supplementary Table S1).

Morphological characterization of pre-anthesis mature spikelets was performed on homozygotes for all mutant alleles, on biallelic *osmads2^i1/d2^* and on heterozygous *OsMADS2^+/d8^* plants. For detailed investigation, we focused on *osmads2^d8/d8^* plants using Scanning Electron Microscopy (SEM) and histological sectioning. Florets from the different homozygous and biallelic mutants consistently exhibited abnormally long lodicules and occasionally displayed increased organ number (Fig. 1B; Supplementary Fig. S2, B-I). The abaxial epidermis of these abnormal lodicules comprised elongated cells that phenocopy cells on the wild-type (WT) marginal tissue of the palea (MTP; Supplementary Fig. S2O). Furthermore, these defective lodicules had reduced thickness and reduced number of vascular bundles (Fig. 1J and Supplementary Fig. S3B). Additionally, we noted instances of cell wall lignification in the mutant lodicule (Fig. 1J) and a transformation of the normally bricklike shaped parenchyma cells that form several internal tissue layers of the wild type lodicule (Fig. 1L) to polygonal cells (Fig. 1M). These mutant phenotypes indicate a partial homeotic transformation of *osmads2* lodicule to MTP which, to a large extent, phenocopy the abnormalities of *OsMADS2* dsRNAi transgenic florets (Prasad and Vijayraghavan, 2003; Yadav et al., 2007), while lodicule phenotype reported in other *osmads2* mutants (Wang et al., 2024a) is not histologically characterized.

**Figure 1.**
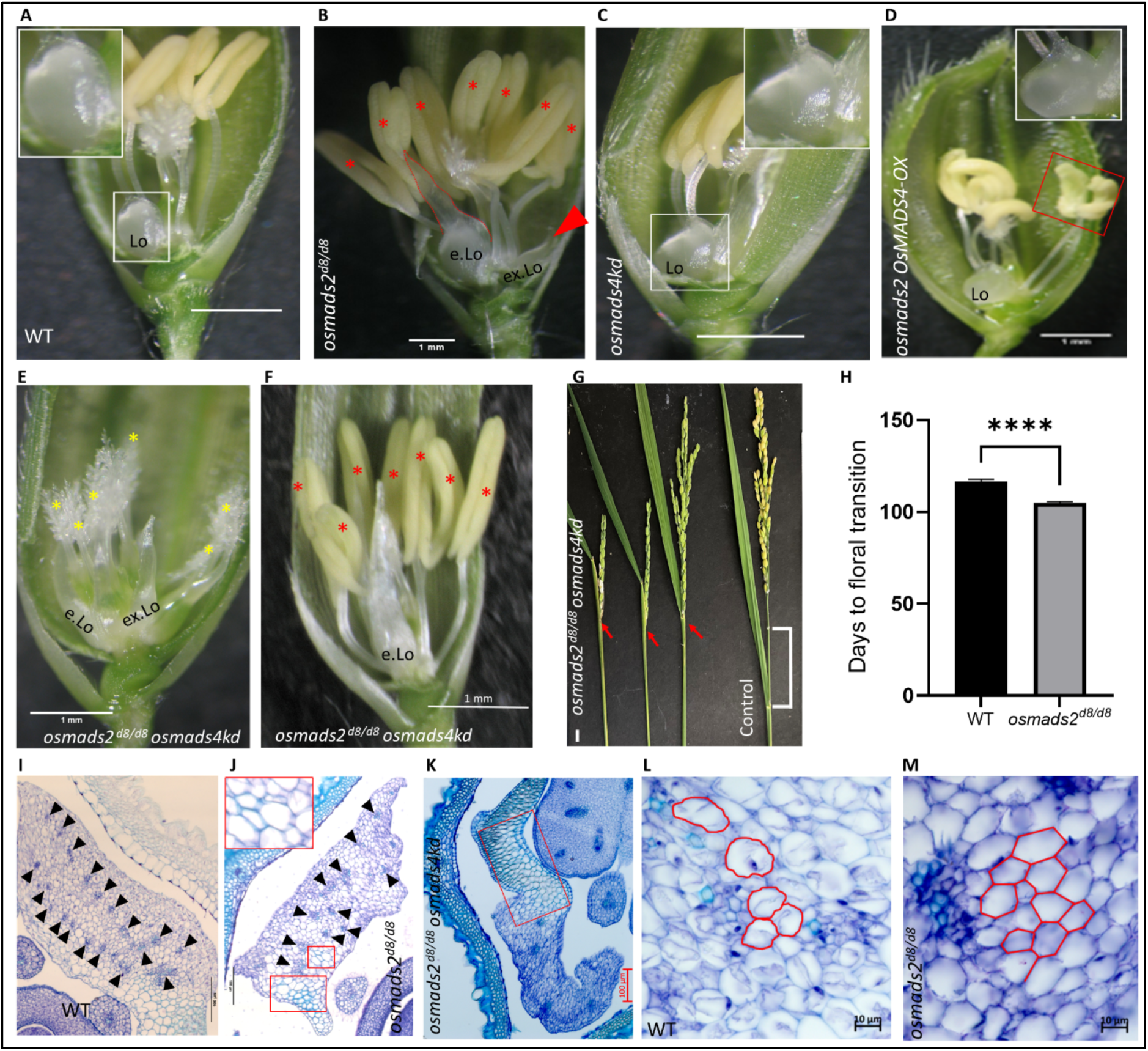
Phenotypic and histological analyses of the different genotypes studied. A, WT mature pre-anthesis spikelet dissected to show the lodicule. The inset shows a magnification of the normal lodicule. B, *osmads2* mutant spikelet dissected to show the abnormal organs. The red asterisks mark the increased stamen number and the red arrowhead marks the extra lodicule (ex.Lo). C, Spikelet dissected from *osmads4kd#1* showing normal floral organs. D, Spikelet from *osmads2 OsMADS4-OX* line #3 showing normal lodicule (complemented) but with abnormal crescent-like anthers (red rectangle). The number of floral organs is unchanged as compared to WT (n=300). E and F, Spikelets from *osmads2^d8/d8^ osmads4kd* of representative group I and group II phenotypes, respectively. The yellow asterisks represent the main and ectopic carpels in group I phenotype and the red asterisks mark the increased stamen number in group II phenotype. The elongated lodicule (e.Lo) is flattened throughout the proximal distal axis (compare lodicule in E and F *vs.* in A and B). G, Panicle partial enclosure phenotype in *osmads2^d8/d8^ osmads4kd* plants. H, Graph plot representing the number of days taken to floral transition in WT plants (n = 79) *vs. osmads2^d8/d8^* plants (n=125). Values are means ± the standard error of the mean (SEM). P < 0.0001 denoted as ****; Student’s *t* test. I-K, TS sections of lodicules in WT and *osmads2^d8/d8^*and *smads2^d8/d8^ osmads4kd*, respectively. The black arrowheads mark the vascular bundles which are reduced in number and of abnormal distribution across the lodicule of *osmads2^d8/d8^*. The red rectangles (in J and K) mark regions with green stained cell outlines, indicative of cell wall lignification in tissue sections stained with Toluidine blue. L and M, TS of lodicules in WT and *osmads2^d8/d8^* at high magnification showing the cell shape in the ground tissues. Compare the brick-like parenchyma in L to the polygonal like cells in M. Cell outlines are marked in red. Scale bars: 1 mm in A-F, 1 cm in G, 100 µm in I-K, 10 µm in L and M.

Unlike lodicules, gross stamen morphology and pollen viability in *osmads2^d8/d8^* plants, and other *osmads2* alleles, were normal and mutant plants were fully fertile (Fig. 1B; Supplementary Fig. S2M; Supplementary Fig. S3J; Supplementary Fig. S4, B and H), as is also recently reported by Wang et al. (2024a). Rarely, the alleles reported here show lodicule-stamen or stamen-carpel chimeric organs and even partial ‘superwoman-like’ phenotype, with reduced stamen number and concomitant increase in carpel number (Supplementary Fig. S2, D and H; Supplementary Table S2). The heterozygous *osmads2^+/d8^* florets were normal though very infrequently (n=2/100) lodicules were mildly elongated and organ number were increased (Supplementary Fig. S2, J and K). These defects in heterozygous florets may arise from stochastic fluctuation in gene expression (Elowitz et al., 2002; Balázsi et al., 2011), the amplitude of which is magnified upon loss of one allele.

Although all *osmads2* mutants had similar lodicule phenotypes, we focussed on *osmads2^d8/d8^* for detailed phenotypic and molecular studies. During inflorescence development, the *OsMADS2* transcript level in the *osmads2^d8/d8^* mutant was reduced by ∼ 4-fold (Supplementary Fig. S1D) and OsMADS2 was not detected in panicle nuclear protein lysates on a Western blot using anti-OsMADS2 antibody (Supplementary Fig. S1E). These data confirm that *osmads2^d8/d8^* is a null mutant. When *osmads2^d8/d8^* mutants were grown alongside WT plants in a greenhouse, while vegetative growth was equivalent, surprisingly, *osmads2^d8/d8^* plants commenced the floral transition nearly twelve days earlier than the WT (Fig. 1H). Interestingly, nearly 5% (n = 533) of *osmads2^d8/d8^*florets exhibited increased inner organ number (Supplementary Table S2), hinting at the possibility of increased cell proliferation or enlarged meristems in *osmads2* florets. Additionally, ectopic lodicules in *osmads2^d8/d8^*florets were positioned medially, opposite to the lateral lodicules, normally present in WT (Supplementary Fig. S3E). This indicates a previously unknown role of *OsMADS2* in mediating the asymmetric phyllotaxy of second whorl organs. Taken together, the in-depth study of developmental phenotypes of mutant alleles in *osmads2* revealed previously unreported processes regulated by *OsMADS2*, *i.e.* floral organ number and flowering time.

### Ubiquitous overexpression of *OsMADS4* complements *osmads2* phenotypes

The two *PI* clade rice paralogs encode proteins that are 81.7% identical and 93.4% similar (Supplementary Fig. S5B). Studies on *OsMADS4*, using knockdown lines, reported some variations with respect to its role in the regulation of lodicule and stamen development (Kang et al. 1998; Yao et al. 2008). Here, we examined *OsMADS4* function employing a specific dsRNAi cassette (Supplementary Fig. S5D). In *OsMADS4 dsRNAi* transgenic line #1 (termed *osmads4kd* hereafter), *OsMADS4* transcript in 2 cm panicles was ∼ 7-fold downregulated (Supplementary Fig. S5C). Despite this robust *OsMADS4* downregulation, *osmads4kd* florets were normal and plants were fertile (Fig. 1C; Supplementary Fig. S2M). These findings are consistent with the only report of mutants in *OsMADS4* locus (Wang et al., 2024a). Collectively, the detailed phenotypic characterization of *osmads2* and *osmads4kd* show that rice *PI* paralogs are fully redundant with respect to stamen specification and could contribute to differing extents for lodicule development.

To further interrogate this differential contribution, we ubiquitously overexpressed *OsMADS4* in the *osmads2^i/d2^* mutant. Interestingly, this complemented the different floral phenotypes of *osmads2*, *e.g*., the abnormal lodicule and the increased floral organ number (Fig. 1D). However, those *osmads2 OsMADS4-OX* transgenics (termed *osm2 OsM4-OX* hereafter) exhibited abnormal, crescent-shaped anthers (Fig. 1D). Histological sections confirmed a normal lodicule vascular pattern (Supplementary Fig. S3H) and revealed unsynchronized and poorly developed anther locules (Supplementary Fig. S3K). Consistent with anther defects, pollen viability and seed set were significantly reduced (Supplementary Fig. S2M; Supplementary Fig. S4, C and H). From this, we conclude that a precisely fine-tuned expression of *OsMADS4* is required for normal anther development.

In the T1 generation, we identified segregants lacking the *OsMADS4-OX* transgene, which we refer to as *osmads2 OsMADS4-tg^-^* (transgene free). As expected, these plants exhibited lodicule abnormalities typical of *osmads2* mutants. We recorded the flowering time of *osmads2 OsMADS4-tg^-^* plants and found that they flowered earlier as compared to WT and *osmads2 OsMADS4-OX* plants, by nearly seven days and ten days, respectively (Supplementary Fig. S6A). These data reiterate the earlier flowering phenotype of *osmads2^d8/d8^* - in other alleles, *i.e., osmads2^i1^* and *osmads2^d2^*, in homozygous or biallelic mutants. Importantly, these results demonstrate that ubiquitous overexpression of *OsMADS4* complemented the *osmads2* early flowering phenotype. Underscoring the contribution of *OsMADS2*, but not *OsMADS4* to flowering time (Supplementary Fig. S6B), is the expression of *OsMADS2* in the vegetative shoot apical meristem (SAM) of wild type 7 days post-germination seedlings, and whose levels are reduced in *osmads2* seedlings (Supplementary Fig. S6C). Further, our analysis of publicly available RNA-Seq data from 6-week-old SAMs of rice (Gómez-Ariza et al., 2019) and 9-day-old SAMs of *Arabidopsis* (Klepikova et al., 2015) showed that *OsMADS2* and *OsMADS16* are expressed in the SAM, while *OsMADS4*, *Arabidopsis PI* and *AP3* are not (Supplementary Fig. S6D). Taken together, these results suggest that *OsMADS2* expression in the SAM prevents a precocious floral transition and identify a diverging, novel function of *OsMADS2* that distinguishes it from *OsMADS4* and *Arabidopsis PI*. Our data on the complementation of different *osmads2* floral phenotypes suggest that the unequal roles of the rice PI paralogs for lodicule development (Yadav et al; 2007; Yao et al., 2008) is likely a consequence of differences in their expression levels and spatial patterns (Yadav et al., 2007) and that they likely encode equivalent proteins.

### *OsMADS4* reinforces lodicule function and identity

To understand the redundant contributions of the rice *PI* paralogs in floral development, we generated a double mutant by crossing *osmads4kd* and *osmads2^d8/d8^* plants. Among the F2 plants, derived from F1 of the genotype *osmads2^+/d8^ osmads4kd,* we identified *osmads2^d8/d8^ osmads4kd* plants. Those plants had no discernible vegetative abnormalities, though floral development was severely altered. The lodicules were severely flattened and elongated throughout their proximal-distal axis (Fig. 1, E and F). Consequence of severe lodicule misspecification as a glume-like organ, all florets remained unopened. Based on the severity of the phenotype in third whorl organs, the *osmads2^d8/d8^ osmads4kd* F2 and subsequent generation plants were classified into two groups: Florets of group I plants displayed strong superwoman-like phenotype, with all or majority of the stamens being replaced by ectopic carpels (Fig. 1E). Florets from group II displayed a milder superwoman-like phenotype, as all or most of the stamens were present but sterile (Fig. 1F; Supplementary Fig. S4, E-G). Segregation analysis of the *OsMADS4 dsRNAi* transgene across generations indicates that the T-DNA is stably inherited as a single insertion locus (65/89 plants carried the *OsMADS4 dsRNAi* transgene) which we found integrated in chromosome 8 intergenic region (Supplementary Fig. S7A). Using insertion site-specific PCR, we inferred that group I (average stamen number 2.4) and group II (average stamen number 5.3) phenotypes are associated with the *OsMADS4* dsRNAi transgene being homozygous and hemizygous, respectively (Supplementary Fig. S7, A and B; Supplementary Table S3).

Interestingly, in the unopened *osmads2 osmads4kd* florets, the stamen filaments did not elongate (Fig. 1F). Moreover, all *osmads2 osmads4kd* inflorescences were compromised for emergence from the flag leaf sheath (panicle enclosed, Fig. 1G), suggesting poor rachis elongation. Together, the phenotypes of enclosed panicles and poor stamen filament extension suggest a compromised hormone signaling of cell elongation in *osmads2 osmads4kd* inflorescences and florets.

We also observed that the inner SL in ∼6% *osmads2 osmads4kd* spikelets were overgrown and greenish, a phenotype seen in only 1.5% in *osmads2^d8/d8^*single mutants (Supplementary Fig. S2L). The identity of the abnormal floral organs in *osmads2 osmads4kd* spikelets were examined by SEM and histological sectioning. Compared to the *osmads2^d8/d8^* single mutant, *osmads2 osmads4kd* lodicules exhibited more severe abnormalities: the epidermal cells of the basal and distal parts of the malformed lodicules were elongated and midveins were observed along the proximal-distal axis (Supplementary Fig. S2P Supplementary Fig. S3C). Furthermore, the number of vascular bundles and the thickness of the lodicules were drastically reduced, the adaxial side was lined with bubble-shaped cells resembling those normally lining the adaxial lemma/palea surfaces, and importantly lodicule cell walls were lignified (Fig. 1K; Supplementary Fig. S3, C and D). The sterile anthers of *osmads2 osmads4kd* group II florets had underdeveloped locules and a persistent tapetum, which normally undergoes cell death in WT (Supplementary Fig. S3L). Moreover, sections of the abnormally enlarged SL revealed three vascular bundles, suggesting that they acquired a palea-like identity (Supplementary Fig. S3N).

Compared to the *osmads2* single mutant, in which only 5.4% of the florets had an increased number of floral organs, ∼ 44% of *osmads2^d8/d8^ osmads4kd* florets (58/130) had an increased number of inner organs (Fig. 1F; Supplementary Fig. S3, C and D). The increased number of organs was traced to an earlier developmental defect of increased floret meristem size in *osmads2^d8/d8^ osmads4kd* florets, as deduced by SEM of young meristems. SEM revealed a slightly enlarged meristem in *osmads2* florets. Notably, the FM size in the *osmads2^d8/d8^ osmads4kd* florets at spikelet stage Sp6 was evidently larger than that in either WT or *osmads2* florets (Supplementary Fig. S8). The enlarged FM size in *osmads2^d8/d8^ osmads4kd* spikelets is reminiscent of the FM enlargement in the *osmads16/spw1* mutant (Yun et al., 2013).

Altogether, we conclude that OsMADS4 can partially contribute to the identity of lodicules and fully support stamen development, likely by forming a functional B-class complex in the absence of OsMADS2, in general agreement with other studies (Yoshida et al., 2007; Wang et al., 2024a). Importantly, by silencing *OsMADS4* in the *osmads2^d8/d8^* null mutant we identify previously unreported roles for these rice *PI* paralogs in rachis elongation, FM size and floral merism.

### Rice *PI* paralogs are required for embryo sac development and suppress parthenocarpy

Florets of *osmads2^d8/d8^ osmads4kd* lines were male sterile due to either the absence of male organs or the lack of viable pollens (Fig. 1, E and F; Supplementary Fig. S4, E-G). To test the female fertility of these plants, carpels were dusted with WT pollens. Surprisingly, there was a complete failure of seed set from this crossing (Fig. 2A), despite the presence of ovule-like structures within the central carpel and some of the ectopic carpels in *osmads2^d8/d8^ osmads4kd* florets (Supplementary Fig. S3, C and D). These data suggest that the sterility of *osmads2^d8/d8^ osmads4kd* florets is not merely due to defective male organs.

**Figure 2.**
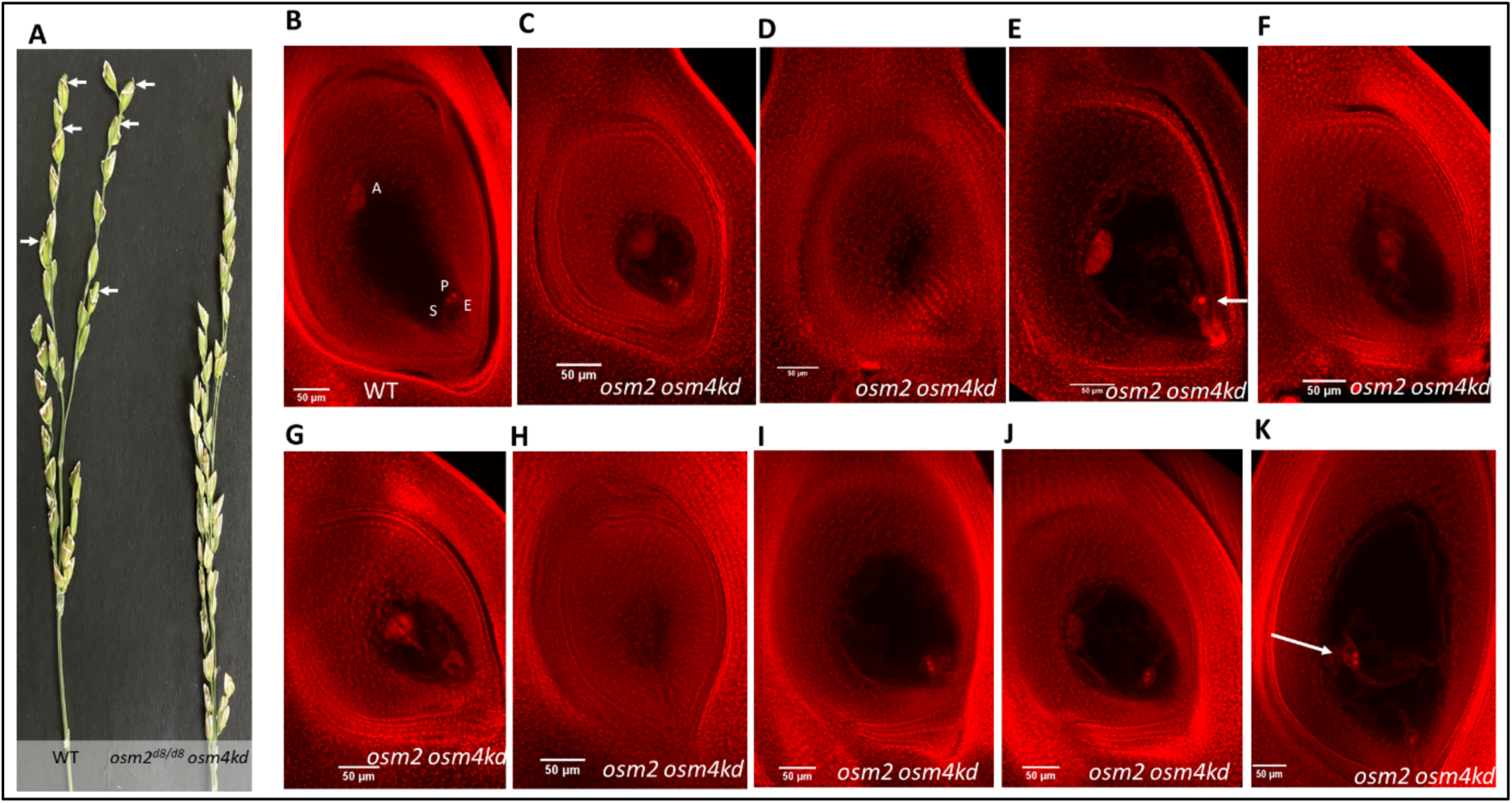
Female sterility in *osmads2^d8/d8^ osmads4kd* double mutants. A, Emasculated WT and *osmads2 osmads4kd* double mutant inflorescences dusted with pollens from WT, show the development of seed (white arrows) in WT but not in *osmads2 osmads4kd* (*osm2 osm4kd*) double mutant. B-K, Ovaries and ovules from pre-anthesis WT and *osmads2 osmads4kd* florets. B, Ovule of WT with normal embryo sac constituting A; antipodal cells, P; polar nuclei, E; egg cell, S; Synergid cells. C, Ovule from *osmads2^d8/d8^ osmads4kd* floret that has apparently normal embryo sac though smaller. D and H, Ovules with degenerated embryo sac (mass of stained cells, no full cavity). Optical sections passing through the center of the embryo sacs for panels D and H are provided in Supplementary Fig. S9, F and G, respectively. E, Embryo sac with single polar nuclei (white arrow). F, Embryo sac with no female unit (no egg cells, no polar nuclei and no synergids). G, Embryo sac with no polar nuclei. I, Embryo sac without antipodal cells. J, Embryo sac with no egg unit (no egg, no synergids). K, Embryo sac with no antipodal cells and with mislocated polar nuclei (white arrow). Scale bars in B-K are 50 µm.

A closer inspection of *osmads2^d8/d8^ osmads4kd* ovaries from fully mature florets revealed that some of the ovule-like structures, in the fourth whorl central carpels, bear undifferentiated nucellar tissue lacking differentiated embryo sacs (Supplementary Fig. S9, C-E). Other ovule abnormalities included embryo sac degeneration, embryo sac without egg apparatus, embryo sac without female germ unit, embryo sac without antipodal cells, embryo sac without polar nuclei, and embryo sac that lacks antipodal cells and have misplaced polar nuclei (Fig. 2, D-K). Interestingly, a few ovules (n=3/10) of *osmads2^d8/d8^ osmads4kd* florets from group II have smaller embryo sacs apparently encompassing the antipodal cells and the female germ unit (Fig. 2C), suggesting they may possess low female fertility or that their infertility arises from defective fertilization or seed/fruit development. RNA in situ hybridization of the rice *PI* paralogs showed localization of *OsMADS2* to the inner ovary walls and the periphery of the developing ovules, and a uniform expression of *OsMADS4* throughout the ovules (Fig. 3), pointing at their contributions to female fertility.

**Figure 3.**
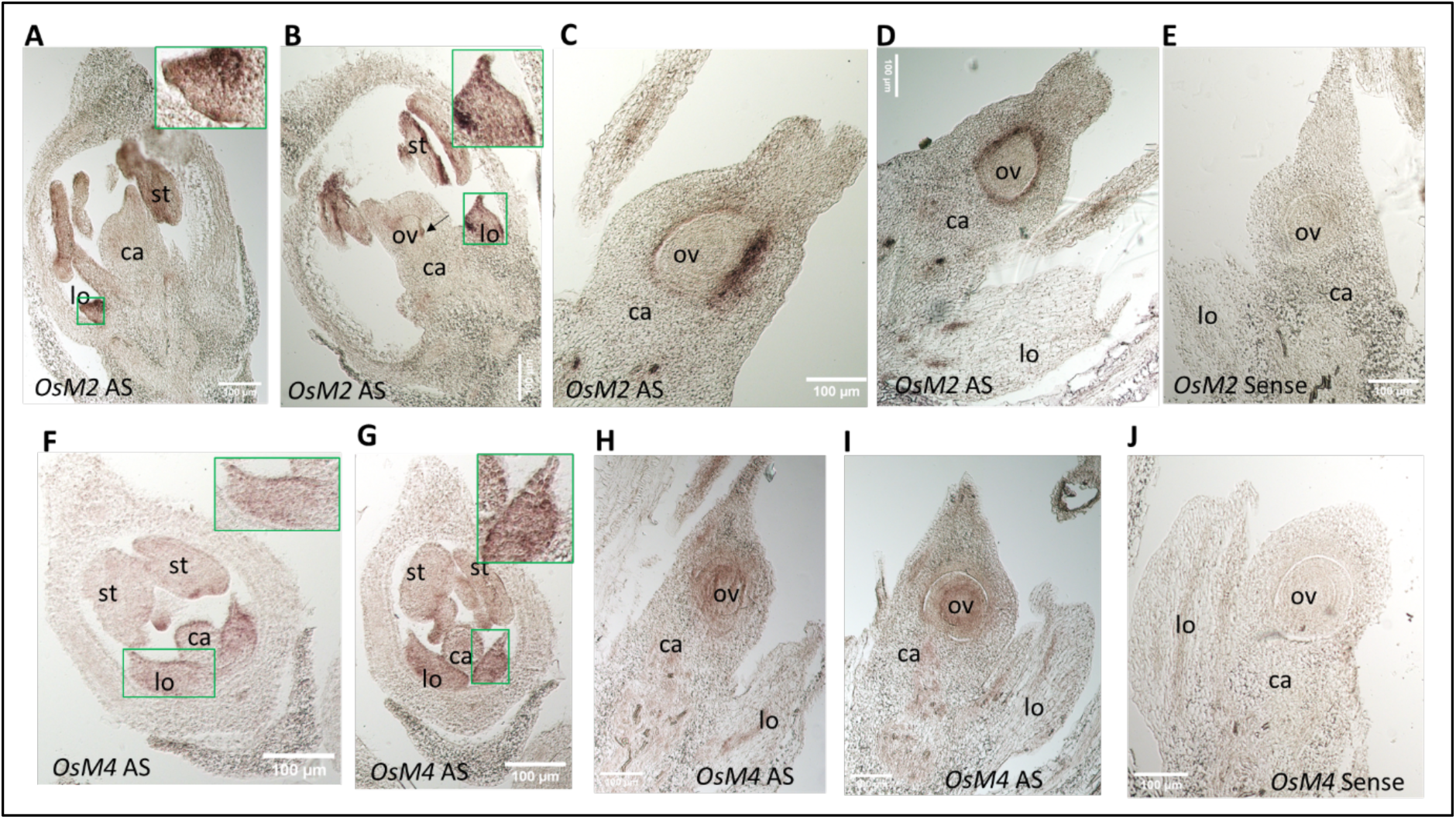
RNA *in situ* hybridization for *OsMADS2* and *OsMADS4* in developing ovaries with ovules. A and B, Sections showing signals for *OsMADS2* antisense probe (*OsM2* AS) in lodicules and stamens. The insets show the expression in lodicules which is enriched distally and at the edges. The black arrow in B points at signals in the initiated ovule. C and D, Sections showing *OsM2* AS signals in the inner ovary and outer layers of the developing ovules. E, Section showing no distinct signals for *OsMADS2* sense probe (*OsM2* Sense). F, Sections showing signals for *OsMADS4* antisense probe (*OsM4* AS) in lodicules and stamens. The insets show uniform expression in lodicules. H and I, Sections showing *OsM4* AS signals in the developing ovules. E, Section showing no distinct signals for *OsMADS4* sense probe (*OsM4* Sense). Abbreviations: lo; lodicule, ca, carpel, ov; ovaries. Scale bars are of 100 µm.

Surprisingly, many of the sterile *osmads2^d8/d8^ osmads4kd* florets of both groups displayed ovary expansion; a response that is normally triggered by fertilization (Fig. 4, A-C). Eventually, these expanded ovaries shrunk and aborted (Fig. 4D). Closer inspection showed that they lacked endosperms and embryos (Fig. 4, G, I and J), suggesting that fruit development initiated without the formation of the products of double fertilization, a phenomenon referred to as parthenocarpy. Altogether, these results point to previously unreported roles of the rice *PI* paralogs in embryo sac and seed/fruit development.

**Figure 4.**
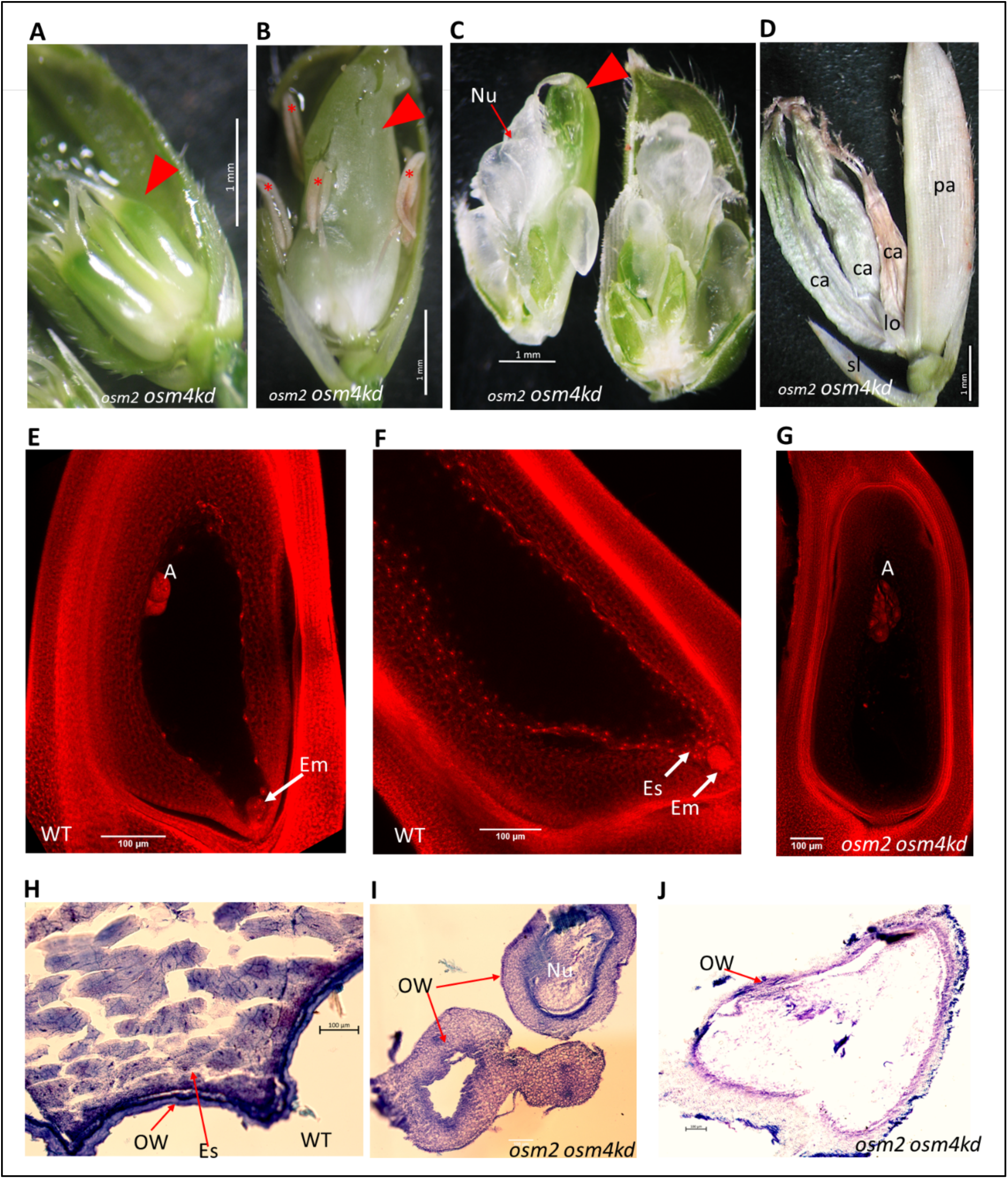
Parthenocarpy-like phenotypes in *osmads2^d8/d8^ osmads4kd* double mutant. A and B, Pericarpic growth in double mutant florets of type I and type II phenotypes, respectively. The red arrowhead points at the expanded pericarp. In B, asterisks mark the indehiscent empty sterile anthers. The filaments of these sterile anthers failed to extend within the unopened florets. C, Floret of *osmads2^d8/d8^ osmads4kd* double mutant with pericarpic growth. The growing ovary (red arrowhead) was manually dissected to show the over-proliferated nucellar mass inside (red arrow). D, Dry spikelet from *osmads2^d8/d8^ osmads4kd* double mutant where lemma is removed to show the expanded pericarps that are shrunken and dry. E-G, Confocal microscopy Z-stacked images of ovaries from WT (E and F) and *osmads2^d8/d8^ osmads4kd* double mutant (G). The double mutant ovary lacks embryo (Em) and endosperm (Es). H and I, Transverse section of WT and double mutant pericarps, respectively. J, Longitudinal section of double mutant expanded pericarp. Unlike WT, the double mutant pericarp lacks granular endosperm. Abbreviation: Carpel (ca), lodicule (lo), palea (pl), sterile lemma (sl), ovary wall (OW). Scale bars: 1 mm in A-D, 100 µm in E-J.

### Mapping genome-wide occupancy of OsMADS2 in developing florets

To shed light onto the mechanistic roles of OsMADS2 as a floral development regulator, we mapped by Chromatin ImmunoPrecipitation followed by deep sequencing (ChIP-Seq) its binding sites, in panicle tissues with florets in a wide range of developmental stages. Peaks of DNA occupancy were called by comparing the enrichment of aligned reads from ChIP over that from a mock immunoprecipitation (IgG). Peaks called from two independent replicates (24,169 and 10,603, respectively) were mapped with respect to transcription start site (TSS) of predicted genes (-3000 to +3000 bp from the TSS). We observed genome-wide pattern of greater enrichment of binding sites proximal to the TSS (Fig. 5A), as reported for other transcription factors, including the *Arabidopsis* Class B *At*PI and *At*AP3 (Kaufmann et al., 2009; Winter et al, 2011; Wuest et al., 2012). We identified 1536 peaks of high confidence that overlapped in the two biological replicates and showed ≥ 2-fold enrichment at a False Discovery Rate (FDR) ≤ 0.05, when compared to the mock IgG. These peaks were assigned to 2589 genes and were located in their vicinity up to 4000 bp from either the TSS or the end of the last exon (Supplementary Data Set 1).

**Figure 5:**
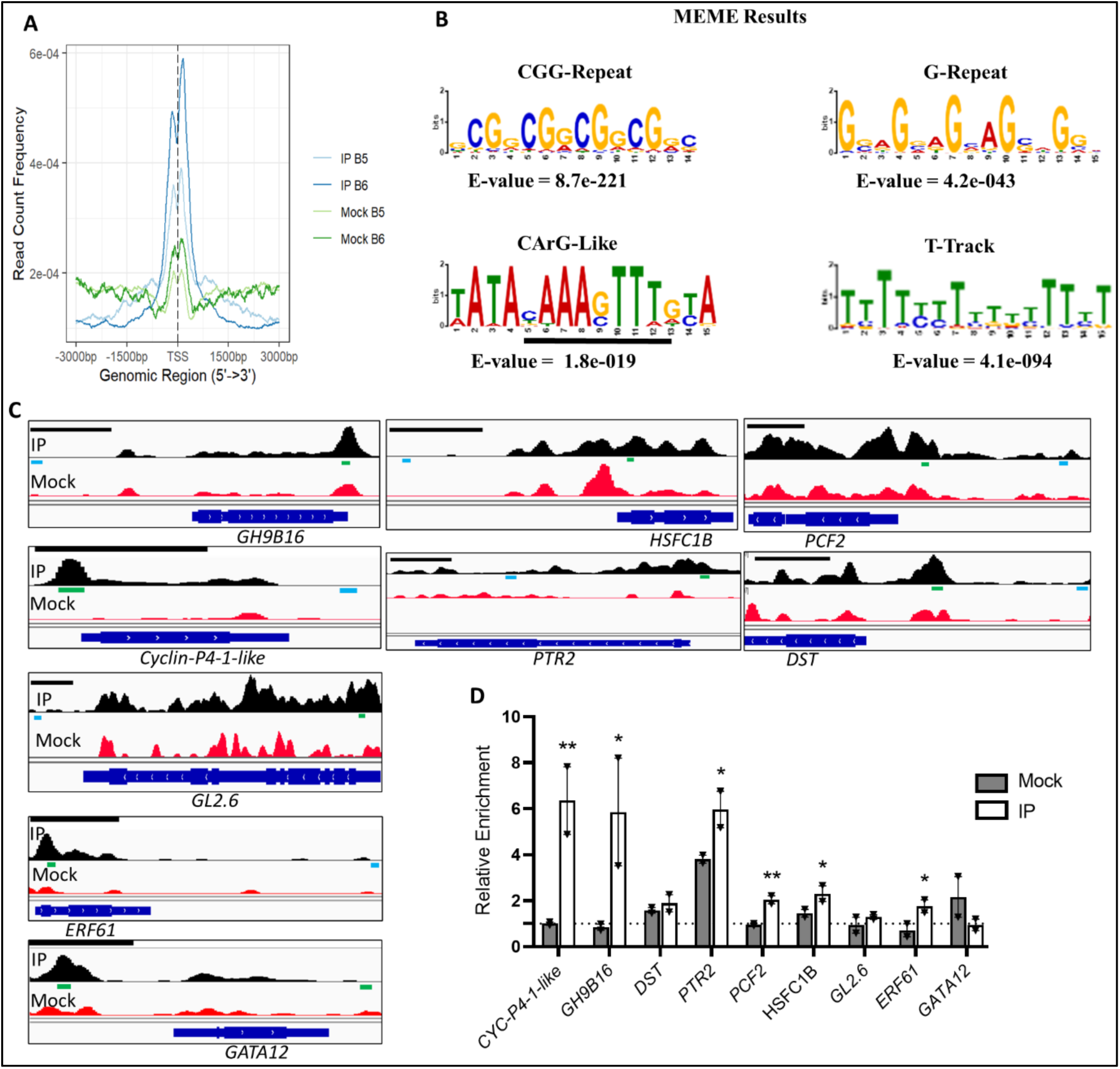
Analysis of genome-wide binding sites of OsMADS2 in WT panicles with developing florets. A, The plot shows relative distribution of OsMADS2 binding sites (ChIP-Seq peaks) mapped relative to TSS (from 3,000 bp upstream to 3,000 bp downstream). The blue lines represent peaks called by MACS2 algorithm using IP data over mock data. The green lines represent peaks called by MACS2 algorithm using Mock data over IP data. The Peaks of the IP over Mock show enrichment around the TSS. B, Significantly enriched motifs in the peak regions of the global data (MEME analysis). C and D, ChIP-qPCR based validation of some genomic loci bound by OsMADS2 in a pool of 0.5-1 cm inflorescences with developing florets. In C, a snapshot of IGV for each locus is shown where the ChIP-Seq peak regions and nonpeak regions (Control) are represented with green and cyan lines, respectively. The black line above each IGV tab is 1kb of the genome. In D, the bar plot represents the relative enrichment in IP and in mock. The relative enrichment (y-axis) is calculated by dividing the %input of the peak region PCR to the %input of nonpeak control region PCR. The dotted line represents a threshold of 1 on the y-axis which corresponds to the %input values of the nonpeak control region PCR after being divided by themselves. The data are retrieved from two independent biological replicates, each includes 2 technical replicates. Values are means ± SEM. Significance of enrichment in IP over mock tested by multiple unpaired Student’s *t* test. P <0.05 and P <0.001 are denoted as * and **, respectively.

We used the MEME-ChIP tool from MEME suite (Machanick and Bailey, 2011) to analyse the DNA sequence from the 1536 high confidence peaks and retrieved significantly enriched sequence motifs (e-value ≤ 0.05). These motifs included CGG-repeats (5’GCGGCGGCGGCGRC3’), T-tracks (TTTTYTTTTTTTTYT), G-repeats (5’GSAGVAGVAGSWGGV3’), and CArG-like (5’TATACAAAGTTTGYA3’) *cis-*elements (Fig. 5B; Supplementary Fig. S10A). The distribution of these motifs at the OsMADS2 binding regions was further investigated by scanning 400 bp segments from 3000 bp 3’ to 3000 bp 5’ of the peak midpoint. Our analyses indicated enrichment of these motifs in the 400 bp window of the peak compared to the 400 bp windows from the flanking sequence (Supplementary Fig. S10B). Surprisingly, no canonical CArG-box [5’CC(AT)6GG3’] motifs were retrieved from MEME analyses. Furthermore, our analyses confirmed that canonical CArG motifs are also not enriched in the 400 bp peak regions compared to the flanking sequences (Supplementary Fig. S10B). In *Arabidopsis*, the genomic binding sites for *At*PI/*At*AP3 are enriched for G-boxes (5′-CACGTG-3′) and GA-repeats (5′-GAGAGAGH-3′, Wuest et al., 2012). Our analyses showed that these motifs are also enriched in peak regions compared to the flanking sequences (Supplementary Fig. S10B). Although MADS domain proteins are known to bind to CArG-boxes, they have been shown to bind non-CArG box sequences *in vitro* (Hill et al., 1998). Together, our analyses do not exclude OsMADS2 binding to canonical CArG-boxes, rather our data suggest preferential association to chromatin that are enriched in other motifs including GCC-repeats and CArG-like sequences.

To validate the ChIP-Seq results, we tested a number of loci associated with the peaks from our genome-wide analyses by ChIP-qPCR on two independent chromatin preparations. These loci were chosen based on their annotations to genes functionally relevant to *OsMADS2* developmental roles. In panicle tissues of 0.5-1 cm, we validated OsMADS2 binding to *GLYCOSIDE HYDROLASE 9B16* (*GH9B16*), *HEAT STRESS TRANSCRIPTION FACTOR C1b* (*HSFC1B*), *PROLIFERATING CELL FACTOR 2* (*PCF2*), *CYCLIN-P4-1-LIKE*, *ETHYLENE RESPONSE FACTOR 61* (*ERF61*) and *PROTEIN TRANSPORTER 2* (*PTR2*, Fig. 5). In 1-2 cm inflorescences, we confirmed OsMADS2 binding to *BETA GLUCANASE 1/* (*EGL1,* Supplementary Fig. S10C). The functions of *PCF2* and *CYCLIN-P4-1-LIKE* in cell division can relate to the abnormalities in floral meristem size, organ number, and lodicule shape observed in *osmads2* mutants. *EGL1* and *GH9B16* encode endo-(1,3;1,4)-β-glucanase and glycoside hydrolase, respectively, are proposed to aid the partial hydrolysis of cell wall components to permit turgor-driven cell expansion (Cosgrove et al., 1999; Akiyama et al., 2009; Xie et al., 2013). Hence, these are important direct OsMADS2 target genes that can facilitate normal lodicule function. Similarly, *PTR2* can account for OsMADS2’s role in vascular patterning (Li et al., 2015) while, *HSFC1B*, as a target of OsMADS2 (Fig. 5, C and D; Fig. 6B), may contribute to osmotic regulation and lodicule function based on its role in the expression of aquaporins and ion transporters under stress conditions (Schmidt et al., 2012). Together, these results provide insights into the molecular roles of OsMADS2 deciphered from its binding sites in developing rice florets that, point to roles in controlling cell wall elasticity, vascular development, osmotic regulation, and cell proliferation.

**Figure 6.**
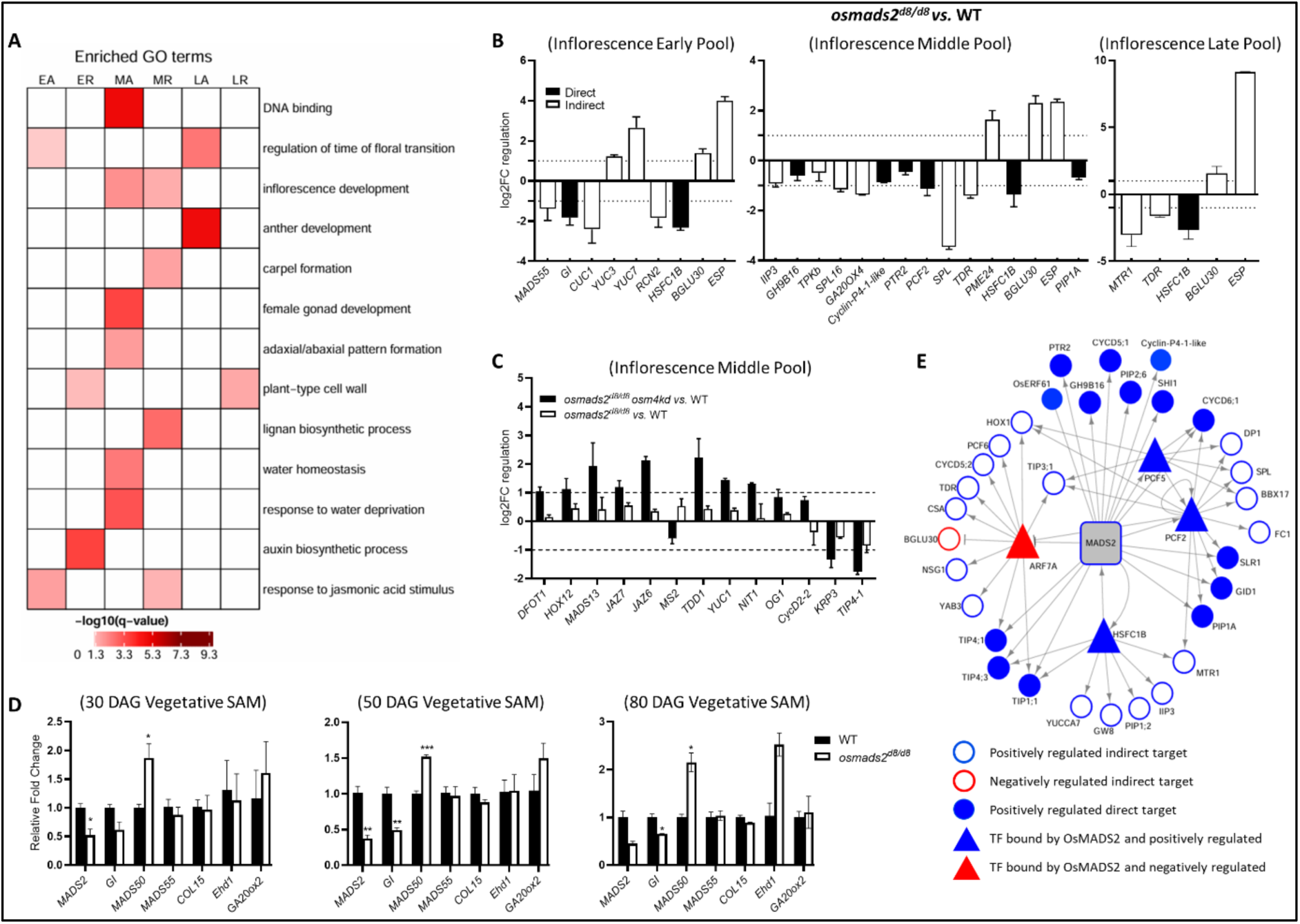
Gene ontology enrichment analysis and target genes regulated by OsMADS2. A, Heatmap representation of selected enriched GO terms in the differentially expressed gene sets. Oryzabase database (https://shigen.nig.ac.jp/rice/oryzabase/) GO annotation was used for this analysis and the significantly enriched GO terms were retrieved at qvalue ≤ 0.05. B and C, Plot representations of selected genes that are differentially expressed in panicles of early (0.2-0.5 cm), middle (0.5-1 cm) and late (1-2 cm) pools of *osmads2^d8/d8^*, as per RT-qPCR based validation of the RNA-Seq data (B), and RT-qPCR analysis of candidate genes redundantly regulated by OsMADS2 and OsMADS4 in floral tissue pool constituting 0.5-1 cm inflorescences (C). Values are means ± SEM. The data are from two biological replicates of *osmads2* or two biological replicates of *osamds2^d8/d8^ osmads4kd*, each includes 3 technical replicates. The average log2FC for three biological replicates of WT is equal to 0. The filled bars in panel B represent those genes that are deregulated as per RNA-seq and RT-qPCR data and bound by OsMADS2 as per ChIP-Seq analysis. D, graph plots displaying the relative expression of flowering time regulators interrogated by RT-qPCR in SAM (meristems and leaf primordia). Values are means ± SEM and derived from three biological replicates except for the plot of 80 DAG, the data are derived from two biological replicates. P <0.05, P <0.01d and P <0.001d denoted as *, ** and ***, respectively; Student’s *t* test. E, A small Gene Regulatory Network (GRN) constructed based on ChIP-Seq data and transcriptome (RNA-Seq) data from panicles of 0.5-1 cm (termed here middle developmental pool). The small GRN displays subset of direct and indirect OsMADS2 target genes of annotated functions in osmotic regulation, cell cycle, cell wall modulation, auxin biosynthesis and other relevant to lodicule and stamen development. The full GRN is presented as Supplementary Fig. 14. The filled blue and red nodes represent positively- and negatively regulated direct downstream targets of OsMADS2, respectively. The unfilled circles in blue and red color boundaries represent genes positively and negatively regulated by OsMADS2, respectively. Triangle shape denotes nodal OsMADS2 target genes coding for transcription factors (for e.g. PCF2 and PCF5) that were used to connect these direct target transcription factors to downstream DEGs that form the second layer in this GRN. The arrow shape and the T-shape indicate positive and negative regulation of each OsMADS2 gene target, respectively.

### Global transcriptome profiling of *osmads2^d8/d8^*

The role of OsMADS2 in the regulation of downstream target gene expression for specific developmental outcomes was studied in the developing panicles by global transcriptome profiling (RNA-Seq), and in the vegetative SAMs *via* RT-qPCR (Fig. 6; Supplementary Data Set 2). In the developing panicles, we compared the transcriptomes of *osmads2^d8/d8^* and WT across three progressive developmental pools. These pools comprised immature inflorescences: 0.2-0.5 cm (termed early hereafter), 0.5-1 cm (termed middle hereafter), and 1-2 cm (termed late hereafter). The early pool contained florets undergoing meristem specification, organ primordia initiation and patterning, the middle pool comprised florets with differentiating organs, and the late pool had florets undergoing organ growth and early gametogenesis. For each developmental pool, RNA-Seq was performed on two biological replicates of *osmads2^d8/d8^* and three biological replicates of WT. The high number of differentially expressed genes (DEG) in the “middle pool” (2215 of the total 2550 DEGs across three pools, at fold-change ≥ 2; FDR ≤ 0.05, Supplementary Data Set 2), is consistent with the high *OsMADS2* levels in these tissues (Supplementary Fig. S11A). This implies important roles for *OsMADS2* during organ differentiation. Among the DEGs, more than 82% of genes at any developmental pool were downregulated (Supplementary Fig. S11B), indicating OsMADS2 acts majorly as an activator of target gene transcription. Interestingly, in all WT tissue libraries, *OsMADS2* had higher expression levels than *OsMADS4* (Supplementary Fig. S11C), pointing to significant differences in the *cis*-elements driving these paralogous genes.

To uncover molecular functions, biological processes, and cellular components controlled by OsMADS2 the DEG datasets were first grouped into those activated or repressed by OsMADS2. Subsequently, gene ontology (GO) enrichment analysis (Fig. 6A; Supplementary Data Set 3) was performed. Several enriched GO terms relate to OsMADS2 roles in lodicule and stamen development. Notable are the terms “water homeostasis” and “response to water deprivation” in the activated DEG subset of the middle tissue pool, the terms “lignan biosynthesis” and “plant type cell wall” in the repressed DEG subsets of the middle and late pools, respectively. These GO terms relate to lodicules as cell wall elasticity and water transport facilitate their rapid swelling during anthesis. Enrichment of the GO term “anther development” in the activated subset of late pool and the term “carpel formation” being enriched in the repressed subset in the middle pool reflect on the stamen development roles for OsMADS2. This is consistent with the homeotic transformation of stamens to ectopic carpels in *osmads2* and *osmads2^d8/d8^ osmads4kd* florets. Phytohormones control both cell proliferation and organ differentiation. Remarkably, the term “auxin biosynthetic process” was enriched in the repressed DEG subset in the early pool (Fig. 6A and Supplementary Data Set 3). Possibly the elevated *YUCCA3* (*YUC3*) and *YUC7* expression in *osmads2^d8/d8^* young florets (Fig. 6B) underlies the increased organ number, given the auxin roles in this process (Krizek, 2011).

We present some OsMADS2 downstream targets that are annotated with relevant functions. Given the partial redundancy of *OsMADS2* and *OsMADS4,* we also discuss some biologically relevant DEGs that are affected at lower threshold of transcript fold change (≥ 1.5-fold change regulation at FDR ≤ 0.05) in *osmads2^d8/d8^*. The precocious flowering of *osmads2^d8/d8^* plants may be linked to lower expression of *OsGIGANTEA* (*OsGI*) and *OsMADS55* in the early inflorescence pool (Fig. 6B; Supplementary Data Set 2). Further interrogation of gene expression revealed steady downregulation of *OsGI* and upregulation of *OsMADS50* in *osmads^d8/d8^* vegetative SAMs of 30-day, 50-day, and 80-day post germination which also had increased expression (P value = 0.053) of *Early heading date 1* (*Ehd1*, Fig. 6D). These data are consistent with *osmads^d8/d8^* early flowering and suggest OsMADS2 could contribute to this trait *via* transcriptional regulation of such flowering regulators (Hayama et al., 2003; Doi et al., 2004; Ryu et al., 2009; Lee et al., 2012) during the vegetative growth.

The abnormal vasculature patterning in *osmads2* lodicules can be related to the deregulation of *PTR2, HOMEOBOX GENE 1* (*HOX1*), and *ILA1 INTERACTING PROTEIN 3* (*IIP3*, Fig. 6B; Supplementary Data Set 2). These genes have vascular development functions (Scarpella et al., 2000; Ning et al., 2011; Li et al., 2015). Cell wall thickening or lignification in *osmads2* lodicules is consistent with the upregulation *PECTIN METHYLESTERASE 24* (*PME24*) and *BETA GLUCOSIDASE 30* (*BGLU30*) (Fig. 6B; Supplementary Data Set 2). Particularly interesting is the report of PMEs regulating floret opening time by modulating pectin methylesterification in lodicules (Wang et al., 2022). Temporal floret opening is aided by lodicule expansion which is driven by the influx of water, associated with the accumulation of K^+^, osmotic solutes such as sugar and di- and tri-peptides, and the increased expression of aquaporins (Heslop-Harrison and Heslop-Harrison, 1996; Choi et al., 2020; Li et al., 2022). In this regard, several DEGs in *osmads2^d8/d8^* inflorescences encoded K+ channels, *e.g. Two-Pore K+ channel b* (*TPKb*, Ahmad et al., 2016), peptide transporter, *e.g. PTR2*, and aquaporins, *e.g. TONOPLAST INTRINSIC PROTEIN 4;1* (*TIP4-1*), *TIP4-3*, *PLASMA MEMBRANE INTRINSIC PROTEIN 1A* (*PIP1A*). The downregulation of these genes in *osmads2* (Fig. 6B; Supplementary Data Set 2) could compromise osmoregulation and turgor pressure in lodicules, and in turn cause the ever-closed *osmads2^d8/d8^ osmads4kd* florets. Notably among the DEGs of the middle pool dataset are *RICE SQUAMOSA PROMOTER-BINDING-LIKE 16* (*SPL16*) and *FLORAL ORGAN NUMBER 1* (*FON1;* Fig. 6B; Supplementary Dataset 2). SPL16 is a positive regulator of cell division (Wang et al., 2012) and its downregulation in *osmads2^d8/d8^* can relate to the reduced thickness or the reduced pararenchymatous layers in mutant lodicules. The reduced expression of *FON1* aligns with increased organ numbers (Suzaki et al, 2004) in *osmads2* and *osmads2 osmads4kd* florets. Consistent with the conserved functions of *PI*-clade genes for third whorl organ differentiation, several DEGs in *osmads2* inflorescences were implicated in anther and pollen development (Fig. 6B; Supplementary Data Set 2). Examples are *TAPETUM DEGENERATION RETARDATION* (*TDR*; Li et al., 2006), *SPOROCYTELESS* (*SPL*; Ren et al., 2018) and *MICROSPORE AND TAPETUM REGULATOR1* (*MTR1*; Tan et al., 2012). Some DEGs in *osmads2* inflorescences may underlie the compromised panicle exsertion seen in *osmads2 osmads4kd* plants. Among these are *EPIGENETIC SHORT PANICLE* (*ESP*), encoding a long noncoding RNA implicated in panicle length (Luan et al., 2019) and the gibberellic acid (GA) biosynthetic genes *GA20ox4* and *GA3ox2* (Fig. 6B; Supplementary Data Set 2).

The exacerbation of lodicule malformation and manifestation of other abnormal phenotypes in *osmads2 osmads4kd* plants, that are neither seen in *osmads2* nor in *osmads4kd* point at other target genes redundantly controlled by both PI-like factors. Towards this end, we curated from the literature functionally relevant genes and interrogated their expression levels in the doubly perturbed *osmads2 osmads4kd* inflorescences (Fig. 6C). For some of the tested genes, *i.e., Cyclin D2-2* (*CycD2-2*), *OPEN GLUME1* (*OG1*) and *MALE STERILITY 2* (*MS2*) differential expression, at a stringent at 2-fold cutoff, was not seen. Importantly, other genes were deregulated at this cutoff and this is consistent with the floral phenotypes observed in *osmads2 osmads4kd*. Among these are *DIURNAL FLOWER OPENING TIME 1* (*DFOT*), involved in lodicule development (Wang et al., 2022), *HOMEOBOX GENE 12* (*HOX12*), involved in panicle enclosure (Gao et al., 2016), *KIP-RELATED PROTEIN 3* (*KRP3*) regulating cell cycle (Mizutani et al., 2010), and auxin biosynthetic genes *YUC1* and *TRYPTOPHAN DEFICIENT DWARF 1*/*TDD1* (Sazuka et al., 2009). Additionally, *MADS13*, a Class D gene (Dreni et al., 2007) had elevated expression in *osmads2 osmads4kd*. However, it is uncertain whether this contributes to the defective ovule development or the excessive ovule proliferation. The auxin biosynthetic gene *NITRILASE 1/NIT1* (Abu-Zaitoon, 2024), the aquaporin *TONOPLAST INTRINSIC PROTEIN 4-1/TIP4-1* and the JAZ6 and JAZ7 encoding Jasmonate ZIM-domain proteins implicated in lodicule development (Cao et al., 2021; Wang et al., 2024b) are examples of genes bound by OsMADS2 (Supplementary Data Set 1) and deregulated in *osmads2 osmads4kd* florets.

The distinct roles of OsMADS2, as compared to *At*PI, suggest significant changes in their downstream targets. We thus compared the DEG datasets in *pi-1* (Wuest et al., 2012) with those in *osmads2^d8/d8^*. Orthologous group analysis indicates nearly 15% of *OsMADS2* downstream targets have orthologs that are deregulated in at least one floral developmental stage of *Arabidopsis pi-1* inflorescence (Supplementary Data Set 4). Examples of such conserved gene targets are *Oryza sativa HOMEOBOX 1* (*OsHOX1*), *Carbon Starved Anther* (*CSA*), and *BASIC HELIX-LOOP-HELIX PROTEIN 120* (*BHLH120*)*. BHLH120* and its *Arabidopsis* orthologs *HEC1* and *HEC2* were upregulated in the *osmads2^d8/d8^* and *pi-1* DEG datasets, respectively. The elevated expression of *HEC* orthologs in *pi* mutants suggest that the suppression of carpel development regulators by B-function genes may be evolutionarily conserved. Remarkably, the orthologs of several rice TCP transcription factor genes which are targets of OsMADS2 were also represented in *pi-1* DEG dataset. Examples are *OsPCF1*, *OsPCF5*, *OsTCP9*, and *OsTCP21*. Notably, orthologs of OsMADS2 downstream targets regulating flowering time *e.g. GI* and *MADS55*, and orthologs of the auxin biosynthetic genes *YUC3* and *YUC7*, were not reported in the *pi-1* DEG dataset. Taken together, these analyses provide clues to some evolutionary conserved or divergent functions of *OsMADS2* as compared to *AtPI*.

### OsMADS2 direct targets relevant to lodicule development and physiology have enhanced expression in the developing lodicules

Nearly 11% of the differentially expressed genes (1.5-fold, FDR ≤ 0.05) from all the three RNA-Seq data sets (Supplementary Fig. S11D), occur in the OsMADS2 ChIP-Seq data set, identifying them as direct gene targets. Yet other DEGs we propose can be regulated by transcription factors that are directly regulated by OsMADS2. We constructed OsMADS2 centric gene regulatory networks (GRNs) that link to certain target genes encoding OsMADS2 transcription factor targets in the early and middle tissue pools (Supplementary Fig. S12; Supplementary Fig. S13; Supplementary Data Set 5). OsMADS2 binding and regulation of some of genes in these GRNs were validated by ChIP-qPCR and RT-qPCR (Fig. 5C; Fig. 6, B and C). Further we present a small OsMADS2 centric GRN (Fig. 6E) that links to target genes implicated for cell division control, osmotic homeostasis, vascular development and other organ differentiation functions that are disrupted in either *osmads2* or *osmads2 osmads4kd* panicles.

The expression profile of some of these genes in wildtype mature floral organs was analysed by RT-qPCR (Fig. 7A; Supplementary Fig. S14A). We find enriched RNA levels in lodicules of pre-anthesis florets for *PIP1A*, *PTR2*, *CYCLINE-P4-*1-LIKE, *PCF2*, *GH9B16*, *HSFC1B*, *TPKb*. Additionally, the spatial temporal expression profile by RNA *in situ* hybridization showed that the transcripts for the direct target genes encoding aquaporin (*PIP1A*), peptide transporter (PTR2), and *CYCLINE-P4-*1-LIKE are abundant during lodicule organogenesis (Fig. 7B-S).

**Figure 7.**
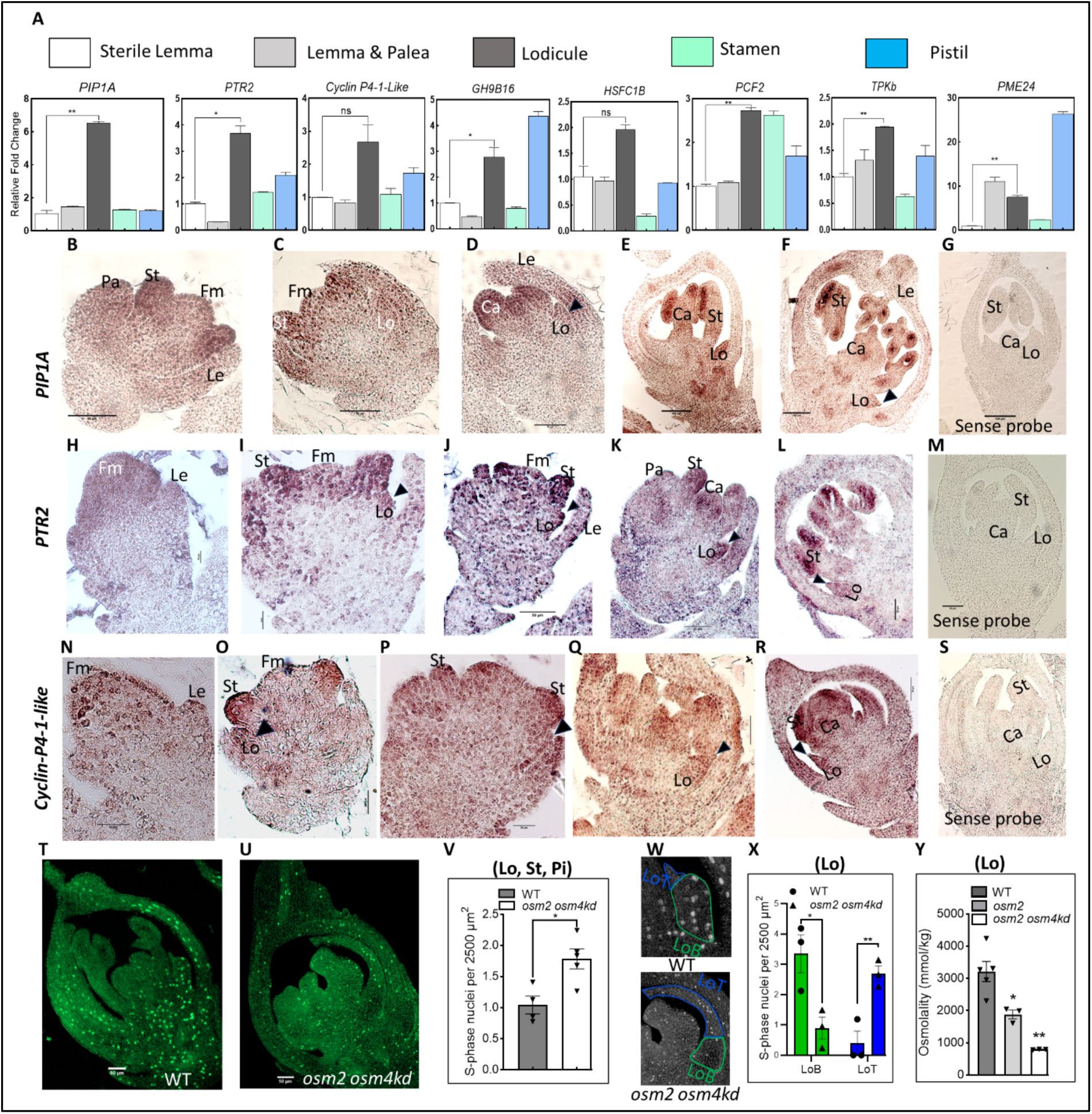
Spatiotemporal gene expression analysis of OsMADS2 targets and spatial distribution of S-phase cells in the developing floral organs. A, RT-qPCR analysis of lodicule developmentally relevant OsMADS2 gene targets examining their relative expression levels in the mature floral organs from pre-anthesis WT florets. The relative fold change is with respect to the expression in the sterile lemma. B-G, RNA *in situ* hybridization of *PIP1A* transcripts in developing WT florets. Sections in panels C-F and panel G are hybridized with antisense and sense probes, respectively. H-M, RNA *in situ* hybridization of *PTR2* transcripts in developing WT florets. Sections in panels H-L and panel M are hybridized with antisense and sense probes, respectively. N-S, RNA *in situ* hybridization of *Cyclin-P4-1-Like* transcripts in developing WT florets. Sections in panels N-R and panel S are hybridized with antisense and sense probes, respectively. The black arrowheads, in B-S, point at the lodicules. T and U, representative confocal micrographs of WT and *osmads2^d8/d8^ osmads4kd* florets, respectively. The florets are of equivalent stages as inferred from their roughly equal dimensions. The S-phase nuclei (bright green signals) are labelled with 5-ethynyl-20 -deoxyuridine (EdU). V, Bar graph plotting the number of S-phase cells per unit area, in the inner floral organs. W, confocal micrographs of WT and *osmads2^d8/d8^ osmads4kd* lodicules demonstrating the distribution of S-phase cells in the lodicule body (LoB, green outlines) and lodicule tip (LoT, blue outlines). The whole florets are shown in panel T and U. X, Bar graph plotting the number of S-phase cells per unit area, in the lodicule body and the lodicule tip. Y, Comparison of osmolality levels in the cell sap from WT, *osmads2^d8/d8^* and *osmads2^d8/d8^ osmads4kd* lodicules. Abbreviations: Fm, floral meristem, Le; lemma, Pa; palea, Lo; lodicule, St; stamen, Ca; carpel, Pi, pistil. Values are means ± SEM. P <0.05 and P <0.01d denoted as * and **, respectively; Student’s *t* test. Scale bars: 20 in I, P; 50 µm in A-D, H, J-O, Q-U; 100 µm in E-G.

Given the large number of cell cycle regulators, as direct and indirect targets of OsMADS2, and the floral or lodicule phenotypes of *osmads2 osmads4kd*, we examined cell proliferation by EdU labelling assay. We compared the number and distribution of cells in S phase in WT and *osmads2 osmads4kd* florets. The data uncovered that the number of S-phase cells per unit area in inner floral organs is significantly higher in *osmads2 osmads4kd* florets (Fig. 7T-V). We further interrogated the spatial distribution of S-phase cells in the lodicule body (LoB) and lodicule tip (LoT), two regions of the lodicule that are identified by differences in the abaxial epidermal cell width and in the number of their inner cell layers (Supplementary Fig. S14B). These data demonstrated significant reduction in the number of S-phase cells in *osmads2 osmads4kd* LoB, concomitant with increased number of S-phase cells in the LoT (Fig. 7W-X). This points at striking change in the spatial cell division of the abnormally flattened and apically elongated *osmads2 osmads4kd* lodicules as compared to the fleshy WT lodicules. Given failure of floret opening in *osmads2 osmads4kd* florets and the finding that a large number of direct and indirect OsMADS2 targets encode aquaporins and osmotic regulators, we determined the osmolality of cell saps from lodicules of WT and *osmads2 osmads4kd* pre-anthesis florets. Sap from *osmads2^d8/d8^*and *osmads2 osmads4kd* abnormal lodicule exhibited significantly reduced osmolality as compared to WT (Fig. 7Y). These data suggest that the failed opening of these mutant florets could arise from the defective lodicule physiology.

## Discussion

### Basis of divergence and novel Class B functions for the rice *PI* paralogs

In this study, we showed that *osmads2* null mutants have abnormal lodicules but unaffected stamens, while *osmads4kd* plants have normal florets as is reported by Yao and colleagues (2008) and also in a very recent report of mutants in *OsMADS4* (Wang et al., 2024a). Together, these data demonstrate that compared to stamen, lodicule development is more sensitive to loss of function or reduction in PI activity and suggest divergence of the rice *PI* paralogs for lodicule development. These studies also reinforce redundant contributions from OsMADS4 for lodicule development are apparent in the context of *osmads2* loss-of-function, as *osmads2^d8/d8^ osmads4kd* florets have more severe derangement of lodicule development. Interestingly, no organ-specific sub-functionalization neither redundancy of the maize *PI* paralogs, *Zmm16/Sts1* and *Zmm18/29*, are reported, since *sts1* mutants have a homeotic conversion of both lodicules and stamens into palea-like organs (Bartlett et al., 2015). Moreover, the rice *PI* paralogs display expression differences that may underlie functional divergence, since *OsMADS2* RNA levels are higher than those of *OsMADS4* (Supplementary Fig. S11D) in all developing panicle pools tested here. Furthermore, differences in expression patterns and abundance in floral organs is apparent in RNA *in situ* hybridization data (Yadav et al., 2007; Fig. 3). Additionally, *OsMADS2*, but not *OsMADS4* is expressed in the vegetative SAM. This we indicate is functionally relevant as we find early flowering in *osmads2^d8/d8^* and normal flowering in *osmads4kd* plants. This role for rice *OsMADS2* in flowering time represents a novel function for a cereal Class B gene. Recently, Shi and colleagues (2024) reported the interaction of daylily PI and AP3 with the flowering suppressor TERMINAL FLOWER 1, in yeast, and their overexpression advance flowering time in *Arabidopsis*, suggesting that Class B MADS factors from other monocot species could also regulate flowering time.

Unlike the lodicule, stamen development is, fully, redundantly controlled by the two rice PI paralogs, since *osmads2* and *osmads4kd* florets have normal stamens as is also seen in various knockdown lines and the recently reported *osmads4* mutants (Yadav et al., 2007; Yao et al., 2008; Wang et al., 2024). However, we report, for the first time, that homeotic conversion of stamen to carpel, in *osmads2 osmads4kd* florets, is associated with the copy number for *OsMADS4* dsRNAi transgene, suggesting that stamen development in *osmads2* mutant is sensitive to the dosage of OsMADS4 activity. By crossing *osmads2^d8/d8^* with *osmads4kd*, we generated stable and fertile *osmads2^d8/+^ osmads4kd* with a hemizygous single insertion of T-DNA for *OsMADS4* knockdown. This line we exploited as a valuable genetic stock to identify double mutant *osmads2^d8/d8^ osmads4kd* segregants over generations, for in depth characterization. In these plants we identified phenotypes, not seen in the single mutant parents of the cross. Thus, we uncover panicle exsertion and suppression of parthenocarpy as novel functions redundantly controlled by the rice PI paralogs. Both are essential physiological processes that ensure high quality grain yield in rice. Panicle enclosure, is a distinctive phenotype seen in nearly all male-sterile rice lines (Liang et al., 2008). It also triggers water stress induced sterility and impair effective pollination, leading to low seed setting (O’Toole and Namuco, 1983). Parthenocarpy, fruit formation from maternal tissue without fertilization, adversely affects reproductive success. Plants evolved several mechanisms by which they counteract this process (reviewed in Joldersma and Liu, 2018). While we show that rice *PI* paralogs redundantly suppress this process (Fig. 8), we point that this could be a conserved function or an example of convergent evolution for B-class activity, as loss of function in PI and AP3 orthologs triggers parthenocarpy in other dicot species, *e.g.* tomato and apple (Yao et al., 2001; Li et al., 2024).

**Figure 8.**
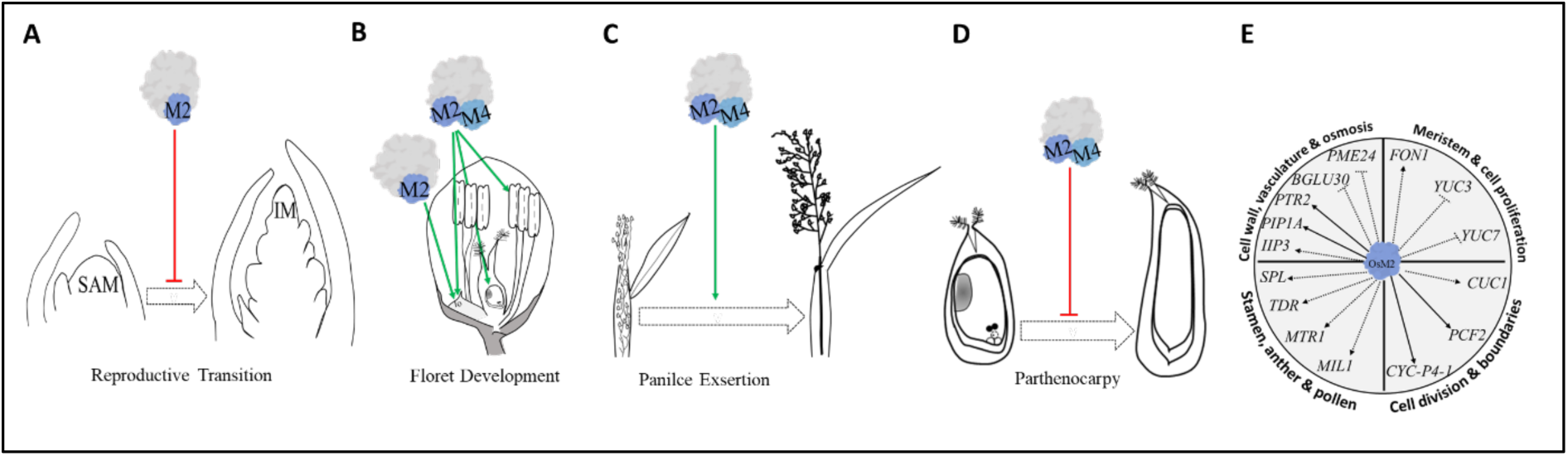
Graphical summary. A, OsMADS2 (M2) as part of a complex negatively regulates floral transition. B, OsMADS2 non-redudant role to promote normal lodicule development. Along with OsMADS4(M4) synergistically promotes normal lodicule, stamen and ovule development. C, OsMADS2 and OsMADS4 redundantly promotes inflorescence exsertion. D, OsMADS2 and OsMADS4 redundantly suppresses parthenocarpy. E, OsMADS2 regulates several downstream target genes that are annotated with functions in cell division, meristem size, meristem proliferation, cell wall modulation, vascular development, osmotic regulation, and stamen development. The solid lines and the dotted lines indicate direct and indirect regulation, respectively. The arrow shape and the T-shape indicate positive and negative regulation of each OsMADS2 gene target, respectively.

### Conserved functions of the rice PI paralogs

Expectedly *osmads2* and *osmads2 osmads4kd* florets had abnormal lodicules and stamens, consistent with the conserved requirement for B-Class function for second and third whorls organ development. However, these mutants also exhibited other abnormalities not reported in studies on these rice *PI* – clade genes. Some of these phenotypes we uncovered are similar to developmental defects reported in some mutants in *PI* – clade genes from other species. Many *osmads2* and *osmads2 osmads4kd* florets had increased stamen organ number, a phenotype seen in the maize *sterile tassel silky ear1* (*sts1*) mutant but not seen in the *Arabidopsis pi-1* mutant (Bowman et al., 1989; Yang et al., 2003; Bartlett et al., 2015). However, the increased second whorl organ number of *osmads2* florets (Fig. 1B) is not observed in the maize *sts1* mutant (Bartlett et al., 2015). Interestingly, in *Nigella damascene*, a dicot species with spiral not whorled flowers, an AP3 paralog influenced perianth organ number (Gonçalves et al., 2013). This observation hints at the likelihood of MADS-box class B genes having a role in the control of floral organ number in diverse species. Furthermore, the emergence of ectopic medial lodicules in *osmads2* mutant restores a whorl of primordia in the second whorl and indicates *OsMADS2* contributes to positional lodicule asymmetry, possibly through asymmetric expression in the second whorl as observed for its maize ortholog, *Sts1* (Bartlett et al., 2015).

Unexpectedly, *osmads2 osmads4kd* were female sterile and displayed varying abnormal ovule differentiation. Although the *Arabidopsis pi* and *ap3* mutants are female fertile (Krizek and Meyerowitz, 1996), mutants in the rice and barley *AP3* homologs, *OsMADS16* and *HvMADS16* (Nagasawa et al., 2003; Selva et al., 2023), are female sterile. While this suggests a conserved female fertility role for class B MADS factors from two related cereal species, the fertile maize *Si1* and *Sts1* mutants (Ambrose et al., 2000; Bartlett et al., 2015), indicate important rice-maize and barely-maize divergences in B-class activity.

The identity of floral organs is dictated by the combinatory functions of different classes of MADS domain transcription factors (Coen and Meyerowitz, 1991; Favaro et al., 2003; Pinyopich et al., 2003). In *Arabidopsis*, although ectopic expression of either *PI* or *AP3* alone does not significantly alter flower development (Jack et al., 1994; Krizek and Meyerowitz, 1996), their simultaneous ectopic expression converted sepals to petals, and carpels to stamens (Krizek and Meyerowitz, 1996). In rice, ectopic overexpression of *OsMADS16* enhanced *OsMADS4* expression and produced multiple stamen-like structures in the fourth whorl (Lee et al., 2003). Interestingly, in our study, ubiquitous overexpression of *OsMADS4* in *osmads2* mutants did not trigger homeotic transformations of floral organs. This indicates that the coexistence of PI and AP3 homologs, at appropriate levels, is a conserved prerequisite for ectopic B-class activity in rice and *Arabidopsis*. Importantly, we have shown that ubiquitous overexpression of the rice *PI* paralog *OsMADS4* led to underdeveloped stamens, a phenotype observed on overexpression of *OsMADS16*, the *AP3* homolog, in rice (Lee et al., 2003). Taken together, these data suggest that excessive, ubiquitous B-class activity is detrimental to normal rice anther development.

### Gene targets of the rice PI paralog, OsMADS2

The genome-wide binding sites and target genes of OsMADS2 identified here reveal its direct and indirect control of the expression of ∼2500 genes that can be assigned to different functional categories. Enrichment of GO terms “sequence-specific DNA-binding” and “transcription factor activity” in the DEG datasets of the early and middle pools (Supplementary Data Set 3), imply that during early floret development, OsMADS2 mediates organ formation, largely by controlling the expression of genes encoding transcriptional regulators. Whereas, during later stages of floret development, it acts more directly on target genes with fewer genes encoding downstream transcription factors, as has been also suggested for the *Arabidopsis* PI/AP3 (Wuest et al., 2012). This is consistent with 3.5% DEGs (1.5-fold, FDR ≤ 0.05) are bound by OsMADS2 from the late RNA-Seq data set as compared to 6.6% and 11% from the early and middle RNA-seq data sets, respectively (Supplementary Fig. S11D). Like seen for the *Arabidopsis pi-1* (Wuest et al., 2012), the number of DEGs in *osmads2* inflorescences increased with progressive flower development, specifically when we compare the early and middle DEG data sets. However, in the late data set, the number of DEGs sharply reduced which could be attributed to the increased heterogeneity of developing tissues in the late pool as it constitutes inflorescences ranging from 1 to 2 cm. Thus, we speculate that the actual number of genes regulated by OsMADS2 activity in late-stage data set may also presumably be similar to that of the middle data set.

The large number direct and indirect OsMADS2 targets imply it participates in the regulation of many pathways required for floral organ initiation and differentiation, consistent with phenotypes seen in *osmads2* and *osmads2 osmads4kd* plants. The negative regulation of the auxin biosynthetic genes *YUC3* and *YUC7* in the early developmental pool leads us to speculate that local auxin levels contribute to the control of floral organ number and the asymmetrical lodicule positioning. However, other organ phenotypes such as the excessive growth of ovule nucellar tissues and the parthenocarpy in the *osmads2 osmads4kd* florets suggest rice PI paralogs dynamically regulate auxin biosynthesis and transport during later stages of floral organ development. As a regulator of lodicule development, OsMADS2 controls the expression of many genes associated with cell wall modification, e.g. *PME24* and *BGLU30* and several *EXPANSINS* (Supplementary Data Set 3). In various studies, the opening time of rice florets was altered by manipulating cell wall characteristics through *PME40, PME42,* and *BETA-1,3-GLUCANASE 4* (*GNS4*, Wang et al., 2022; Xu et al., 2022). Therefore, the OsMADS2 targets (*PME40, PME42* and *BLGU30*) offer promising future screens for cleistogamy florets and/or for lines with advanced floret opening time. Such traits would facilitate breeding and mitigate heat-induced floret sterility. Modulation of genes for cell wall characteristics is associated with response to drought, as inferred from different omics data sets (reviewed in Veronico et al., 2022). In this context, it is worth noting that OsMADS2 regulates many genes associated with response to water stress (Supplementary Data Set 3). Such genes along with other targets involved in cell wall metabolism and the transport of K^+^ and osmotic solutes, may collectively facilitate lodicule function. From these categories, we interrogated the expression of various direct targets: *GH9B16*, *PME24*, *PIP1A*, *PTR2* and *TPKb*, and showed their enrichment expression in lodicules. The floret opening mediated by lodicule expansion coincides with stamen filament elongation. Like reported by Zhao and colleagues (2022), stamen filaments also failed to elongate in the unopened *osmads2^d8/d8^ osmads4kd* floret (Fig. 4B). This together suggest common triggers for cell expansion in lodicule and stamen filaments. The activity of these genes, as they enable cell wall loosening, water influx and cell expansion, may control these timely synchronised processes.

Although OsMADS2 targets constituted genes encoding various transcription factor families, the MADS-box gene family, despite being represented in the ChIP-Seq data set, were depleted from the three progressive DEG data sets (2-fold change; FDR ≤ 0.05) This implies that OsMADS2 could be regulating other MADS-box genes redundantly with other factors, *i.e.* OsMADS4 and OsMADS16. Given the physical interaction of OsMADS2 and OsMADS32/CFO1 (Wang et al. 2015), and that *OsMADS2* levels are normal in *cfo1* (Hu et al. 2021), the shared phenotypes (abnormal lodicules and chimeric floral organs) of *osmads2* and *cfo1*, imply that OsMADS2 and OsMADS32 function in a complex to regulate lodicule development and floral organ identity. In conclusion, the target genes of OsMADS2 unravelled in this study provide new insights into the mechanistic role of *OsMADS2* in rice floral development (Fig. 8). Importantly, stable and fertile *osmads2* mutants and the doubly perturbated *osmads2^d8/+^ osmads4kd* plants will enable future genetic and gene regulatory studies on the temporal and spatial control of tissue differentiation in floral organs.

## Materials and Methods

### Plasmid construction and plant transformation

For CRISPR/Cas9 gene editing of *OsMADS2*, two pairs of primers were designed for respective sets of 20 nucleotide guide RNAs (gRNAs). The first pair of primers (a and b) were annealed and cloned into the commercial pRGEB31 vector to construct a binary vector used to transform rice plants with intended editing in the OsMADS2 DNA-binding domain (designated gRNA1). The second pair of primers (c and d) were annealed and cloned similarly, and the construct was used to generate transgenic plants with intended editing in OsMADS2 K-domain (designated gRNA2). The sequences for the different oligos are given in Supplementary Table S4. For *OsMADS4* dsRNAi-based knockdown, a 290 bp specific fragment (from +1794 to +2083 of the *OsMADS4* genomic locus) was amplified using primers *OsMADS4* FP and *OsMADS4* RP (Supplementary Table 4). This fragment was then cloned into pBluescript in opposite orientations separated by a linker. The generated clone was verified by restriction digestion and Sanger sequencing. Subsequently, the insert in recombinant pBluescript was released and cloned into the binary rice expression vector pUN (Prasad et al., 2001), downstream to the maize ubiquitin promoter (*pZmUbi*), to express the *OsMADS4* hairpin RNA. For overexpression of *OsMADS4* in the *osmads2* mutant background, the *OsMADS4* cDNA was amplified with primers *OsMADS4* CDS FP and RP (Supplementary Table S4), from cDNA transcribed from TP309 panicle tissue RNA, and cloned into pBluescript. A clone verified by restriction digestion and Sanger sequencing was used to release *OsMADS4* cDNA, which was subsequently cloned downstream of the *pZmUbi* into a modified pCAMBIA3300 vector with a *pZmUbi* promotor and a *Nos* terminator. This Basta resistant vector was used to transform rice calli generated from mature seeds of *osmads2* mutant. Binary vectors were mobilized into the LBA4404 *Agrobacterium tumefaciens* strain, which was used to transform WT rice (*Oryza sativa* var. Japonica) TP309 embryogenic calli and transgenic lines were raised as described by Prasad et al. (2001). Plants were grown as reported by Prakash et al. (2022). For raising anti-OsMADS2 antibody, a gene-specific *OsMADS2* cDNA fragment encoding the amino acids 112 to 209 was cloned into pET-15b and the His-tagged protein was expressed in bacteria grown with 0.5 mm isopropylthio-β-galactoside at 37°C for 3 hours. The His-tagged OsMADS2 partial protein was purified by nickel-nitrilotriacetic acid agarose chromatography, followed by the removal of the His tags by thrombin cleavage. The purified proteins were used to raise rabbit polyclonal antibodies that were affinity purified using the antigen. The specificity of the purified antibody was established by Western blots on nuclear proteins from panicles (Supplementary Fig. S1E).

### Genotyping of CRISPR/Cas9 edited *OsMADS2* lines

Genomic DNA regions flanking the editing sites were PCR-amplified using primers *OsMADS2* E3-I3 FP and *OsMADS2* IV RP (Supplementary Table 3). The PCR amplicon was digested with *Sac*I to identify putative edits in *OsMADS2*. Subsequently, the PCR amplicon from putatively edited plants was cloned into *Eco*RV site of pBluescript and at least two pBSKS clones were given for Sanger sequencing to identify the exact edits. To identify plants that had transmissible edits in the germline, independent genotyping was done for two subsequent leaves of the same plant and across different generations. Once mutant plants with stably inherited alleles were identified, Sanger sequencing of PCR products was utilized for further genotyping in advanced generations.

### Amplifying the genomic DNA flanking the T-DNA integration site

Restriction Site Extension PCR (Ji and Braam, 2010) was utilized to amplify the genomic DNA flanking the T-DNA harbouring the *OsMADS4* dsRNAi cassette. Briefly, primers p1300 5.0 and p1300 5.1, specific to the pCAMBIA1300 left border (Supplementary Table 3), were used to perform the primary and secondary PCRs, respectively. Two amplicons of ∼ 1.2 and ∼ 1.4 kb were obtained from genomic DNA digested with *Sac*I and *Kpn*I, respectively. These amplicons were gel-purified and Sanger sequenced. The genomic sequence flanking the T-DNA left border was then mapped to the rice genome using blastn. To screen transgenic plants for hemi- or homozygosity with respect to the *OsMADS4* dsRNAi cassette, two primers (*OsM4* ds IS FP and RP; Supplementary Table 3) flanking the insertion site were designed and utilized to amplify the WT genomic sequence from the hemizygous segregants.

### Phenotypic Characterization

Freshly collected pre-anthesis mature spikelets were examined and photographed using a Leica Wild M3Z stereomicroscope fitted with digital camera (Canon Powershot G5). Pollens from rice anthers at anthesis were stained with either iodine potassium iodide (I_2_KI) or 4′,6- diamidino-2-phenylindole (DAPI) for 1-5 mins on a microscope slide and examined under the microscope. Phenotyping of ovaries and ovules by whole-mount eosin B-staining confocal laser scanning microscopy was done according to Zeng and colleagues (Zeng et al., 2007). For SEM, fresh spikelet and inflorescence samples were fixed in 0.1 M sodium phosphate buffer at pH 7.0 containing 3% glutraldehyde (Sigma-Aldrich, St. Louis, USA) for 12-24 hrs. Tissue samples were then post-fixed in 1% osmium tetroxide for 24-48 hrs or until performing the next step. Next, tissue samples underwent graded ethanol dehydration and critical point drying using Pelco CPD2 (Ted Pella, California, USA). Critical point dried samples were mounted on an SEM stub, using double stick conductive carbon tape, sputter coated, and examined. An Ultra 55 FE-SEM with EDS instrument (Carl Zeiss, Oberkochen, Germany) was used for imaging with help from the technician in charge at the Centre for Nano Science and Engineering (CeNSE) at IISc. For histological analysis, tissues were fixed in FAA solution (10% Formalin, 5% Glacial Acetic Acid, 50% Ethanol), dehydrated in an ethanol series, and embedded in paraffin. Sections of 8 um were stained with 0.5% toluidene blue (FisherBiotech, Massachusetts, USA).

For 5-ethynyl-2’-deoxyuridine (EdU) labelling, Click-iT® EdU Alexa Fluor® 488 Imaging Kit (ThermoFisher, C10337) was utilized. Panicle tissues of 0.7 cm length were excised and placed in 25 µM EdU solution made in half strength MS (Murashige and Skoog) for 2 hrs, then fixed and embedded in paraffin as described above. Sections of 8 um were dewaxed and processed for detection according to the manufacturer’s instructions. Sections from WT and *osmads2^d8/d8^ osmads4kd* florets of equivalent developmental stage (having roughly equal dimensions) were imaged with Zeiss 710 confocal microscope. Number of S-phase cells in the inner floral organs or in the lodicule were counted manually across all sections of a single floret. The area of the inner floral organs, the area of the tip of the lodicule and the area of the body of the lodicule across all sections of a single floret were measured in FiJi (ImageJ). The number of S-phase cells per unit area in the inner organs were plotted and statistically analyzed for 4 WT florets and 5 *osmads2 osmads4kd* florets, while in case of lodicule the N was equal to 3. For measuring the osmolality of the sap from the lodicules, nearly 80 pairs of lodicules, per biological replicate, were dissected from pre-anthesis florets collected between 09:00-11:00 am. Lodicules were then crushed in a 1.5 ml microcentrifuge tube with an autoclaved micro-pestle, followed by a quick spin at 6000g. The sap was then collected and the osmolality was measured with a vapour pressure osmometer (Vapro5600, Wescor). Three biological replicates of *osmads2^d8/d8^* and *osmads2^d8/d8^ osmads4kd* lodicules were plotted and compared to five replicates of WT.

### Western Blotting

Nuclear proteins were isolated according to Busk and Pagès (1997). Protein concentration was estimated by the Bradford method according to the manufacturer’s instructions (Sigma Aldrich, Missouri, USA). The protein was separated on an SDS PAGE gel of 12% and transferred to PVDF membrane with a 0.5 µm pore size through Trans-Blot® SD semi-dry transfer cell (Bio-Rad, USA). Blocking was done for 1 hr, followed by incubation with the primary antibody (polyclonal affinity-purified primary antibody against the 11.44 kDa C-terminal region of OsMADS2) for 12 hrs. Washing was done for 3X for 5 min, after which the membrane was incubated with an HRP-conjugated secondary antibody for 1 hr. The membrane was then washed 3X for 5 min each and developed using the Immobilon Forte Western HRP Substrate (Cat# WBLUF0100, Millipore, USA). Images were acquired using an ImageQuant LAS4000 Chemidoc instrument (GE HealthCare Bio-Sciences AB, Japan) of the Microbiology and Cell Biology Department at IISc.

### ChIP-sequencing, read alignment, peak calling, and gene assignment

Wild-type TP309 panicles were used for the ChIP. The tissue pool ∼200mg constituted 4 panicles (x) 0.3cm, 4x 0.4cm, 3x 0.5cm, 1x 0.6cm, 1x 0.7cm,1x 0.8cm, 1x 0.9cm, 1x 1cm, 1x 1.5cm and 1x 2cm. The tissues were fixed with 1% formaldehyde in a fixation buffer. OsMADS2 ChIP was performed using SimpleChIP® Enzymatic Chromatin IP Kit (Cat #9002, CST, Massachusetts, USA). MNaseI based chromatin was prepared with DNA fragments ranging from 150bp to 900bp. For the ChIP, 8μg of chromatin was used and incubated with 6μg of affinity-purified anti-OsMADS2 antibody. The DNA-protein complexes were pulled down using protein-A magnetic beads. Simultaneously, a negative control sample using rabbit IgG antibody (mock IP) instead of the anti-OsMADS2 antibody was processed in the same way. Immunoprecipitated DNA was eluted using the elution buffer provided in the kit and reverse cross-linked using 2M RNase A and 4M proteinase K in 5M NaCl for 6 hours at 65°C. Reverse-crosslinked ChIP DNA was purified using the phenol/chloroform purification and precipitation method. Approximately 6 ng of ChIP DNA was processed to prepare a library for NGS by using the Kapa kit (Roche, Basel, Switzerland) following the manufacturer’s instruction with one modification: library amplification was performed before the double-size selection. NGS was performed using the Illumina HiSeq2500 machine at the Functional Genomics Center Zurich. Sequencing depth ranged from 22 million to 39 million single-end reads per sample. Raw fastq files were processed with FASTQC and Fastp for filtering low quality reads and adaptor trimming. Next, short read alignment was performed by bowtie2 version 2.3.5.1 using MSU-TIGR v7 as the reference genome. The bam files with alignment information were process with SAMBAMBA (0.6.6) for sorting and filtering uniquely mapped reads. For peak calling, the uniquely mapped reads were processed with MACS2 (version 2.2.6; Zhang et al., 2008) using the default parameters, keep-dup parameter as AUTO, and effective genome size of 3.50e+08. Briefly, two biological replicates (B5 and B6) of IP and Mock-IP were used. First, peaks from IP B5 were retrieved using Mock-IP B5 as a control. Another set of peaks from IP B5 were retrieved using Mock-IP B6 as a control. Next, peaks from IP B6 were recovered using Mock-IP B6 as a control. To determine confident peaks from B5 IP, overlapping peaks from IP B5 over Mock-IP B5 and IP B5 over Mock-IP B6 were recovered and merged. These confident peaks from IP B5 were then merged with overlapping peaks from IP B6. These overlapping, merged peaks from IP B5 and IP B6 were designated as peaks of high confidence. Next, these peaks were assigned to genes in their neighbourhood up to 4000 bp from either the start or end of the gene. Finding overlapping peaks and assignment of peaks to genes were performed in R studio using the ChIPpeakAnno library (Zhu et al., 2010) functions, findOverlapsOfPeaks and annoPeaks, respectively. For annoPeaks function, bindingType and bindingRegion were set to “fullRange” and to -4000 and +4000, respectively. Peak distribution with respect to TSS and 3000 bp flanking regions was performed with plotAvgProf from the ChIPseeker Bioconductor package (Yu et al., 2015), using coordinates from “Oryza_sativa.IRGSP-1.0.53.gtf.gz”.

### ChIP-Seq motif analyses

Coordinates of the confident peaks were defined from IP B5. The midpoints of these peaks were determined and a set of 500 bp (-250 to 249 from the midpoint) sequences was extracted. In these 500 bp sequences, the repeats were masked using RepeatMasker (https://www.repeatmasker.org/) and then submitted to MEME-ChIP from the MEME suite (Machanick and Bailey, 2011). From MEME analyses, we retrieved significantly enriched motifs with E-values ≤ 0.05. Analysis of motif distribution in peak regions and flanking 3000 bp at 400 bp windows utilized the R function vmatchPattern from the Biostrings Bioconductor package (Pagès et al., 2022).

### RNA-Sequencing and orthologous group analysis

RNA-sequencing was performed for three progressive panicle pools, named early, middle and late. The early pool constituted 3 panicles (x) of 0.2 cm, 4x 0.3 cm, 4x 0.4 cm, 4x 0.5 cm. The middle pool constituted 1x 0.5 cm, 2x 0.6 cm, 2x 0.7 cm, 1x 0.8 cm, 1x 0.9 cm, 1x 1 cm. The late pool constituted 1x 1cm, 1x 1.2 cm, 1x 1.5 cm, 1x 2cm. For each pool, the number of biological replicates of WT and *osmads2^d8/d8^* tissues was three and two, respectively. First, total RNA was extracted from panicle tissue using the TRIzol R reagent (Invitrogen, Carlsbad, CA, USA). RNA was then treated with DNAse I (M0303S NEB, Massachusetts, USA) to remove contaminating gDNA according to the manufacturer’s protocol. RNA quality and concentration were examined on a 4150 TapeStation (Catalog: G2992AA, Agilent Technologies, California, USA) and Qubit (Life Technologies, California, USA), respectively. RNA of integrity number (RIN) > 7 was used for library construction. Total RNA of 500 ng was used to enrich mRNA using the NEBNext Poly (A) mRNA magnetic isolation module (Catalog: E7490, New England Biolabs) following the manufacturer’s protocol. Enriched mRNAs were used for library preparation using the NEBNext® Ultra™ II RNA Library Prep Kit for Illumina (Catalog: E7775S, New England Biolabs). Sequence data was generated using the Illumina HiSeq, pair-end 2X150bp chemistry. Sequencing depth ranged from 14 million to 31 million paired-end reads per sample. Data was checked for base call quality distribution, percentage of bases above Q20, Q30, %GC, and sequencing adapter contamination using FastQC and MultiQC software (Ewels et al., 2016). All the samples passed the QC threshold (Q30>90%). Raw sequence reads were processed to remove adapter sequences and low-quality bases using fastp (Chen et al., 2018). The QC passed reads were then mapped onto the indexed *Oryza sativa* ssp. japonica cv. reference genome (RAP-DB; https://rapdb.dna.affrc.go.jp/), using the STAR v2 aligner (Dobin et al., 2013). On average, 99.9% of the reads aligned to the reference genome. Raw reads were counted with the featureCounts function from the Rsubread package (Liao et al., 2019). Differential expression analysis was carried out using the edgeR package (Robinson et al., 2010) after normalizing the read counts based on trimmed mean of M (TMM) values. Replicates from *osmads2^d8d/d8^* were compared to replicates from WT. Genes with an absolute log2 fold change ≥ 1 and an adjusted p-value ≤ 0.05 were considered significant. The RNA-Seq raw data from 6-week-old SAMs of rice (Gómez-Ariza et al., 2019) were downloaded from NCBI and mapped as described above. The RNA-Seq raw data from 9-day old SAMs of *Arabidopsis* (Klepikova et al., 2015) were downloaded from NCBI and mapped to the TAIR9 Genome Release. Hypergeometric GO enrichment analysis was conducted by using GO annotations of Oryzabase (https://shigen.nig.ac.jp/rice/oryzabase/locale/change?lang=en) as the global dataset, and *osmads2* DEG datasets as local datasets. Significantly enriched GO terms (qValue ≤ 0.05) and the associated genes were curated in a supplemental dataset. For orthologous group analysis, a list of orthologous groups (file name: APK_ftp_file.txt) was downloaded from the Rice Genome Annotation Project (http://rice.uga.edu/annotation_pseudo_apk.shtml). This list was reorganized to contain only rice genes with corresponding *Arabidopsis* orthologs, and then downsized to only those rice genes that were differentially expressed in at least one stage. This list was then overlapped with the *pi-1* dataset (Wuest et al., 2012) to identify the rice downstream targets of OsMADS2 that have an ortholog represented in *pi-1* data set of differentially expressed genes.

### RT-qPCR, ChIP-qPCR and RNA *in situ* hybridization

Synthesizing of cDNAs, RT-qPCR reactions and calculation of fold-change were performed as described in Prakash et al. (2022). Primers used and their sequences are given in Supplementary Table S4. For ChIP-qPCR, WT TP309 panicles of length 0.5-1 cm or 1-2 cm (∼200mg) were used. The tissues were fixed with 1% formaldehyde in fixation buffer. OsMADS2 ChIP was performed using the SimpleChIP® Enzymatic Chromatin IP Kit (Cat #9002, CST, Massachusetts, USA). MNaseI-based chromatin was prepared with a DNA size range of 150bp to 900bp. For the ChIP, 2 µl were kept as input DNA and 90 µl of chromatin was taken and incubated with 6μg of affinity-purified anti-OsMADS2 antibody overnight. The DNA-protein complexes were pulled down using protein-A magnetic beads. Simultaneously, a negative control sample (Mock IP, no primary antibody) was processed similarly. Immunoprecipitated DNA was eluted using the elution buffer provided in the kit and reverse cross-linked using 2M RNase A, 4M proteinase K in 5M NaCl for 6 hours at 65°C. The DNA was then purified using column purification. For ChIP-qPCR, the final ChIP DNA was diluted 10 times and 1 µl was used for qPCR with locus-specific primers (Appendix D). For each tested locus, two regions (peak and nonpeak) were amplified using DNA from input, IPed, and mock samples. First %input was calculated as follows: %input for IP samples= 2^(CT.in-log2(DF)-CT.ip) and %input for Mock samples= 2^(CT.in-log2(DF)-CT.mock). CT.in, CT.ip, and CT.mock refer to the average CT value for input, IPed, and mock samples respectively. DF refers to the dilution factor of the input. The relative enrichment was then calculated by dividing the %input in the peak region to %input in the nonpeak region. Hence, the values for the nonpeak regions were all set to 1. The data were derived from two independent biological replicates of OsMADS2 ChIPed DNA along with respective Mock sample, each include two technical replicates. The *in situ* hybridization experiments were performed as described by Yadav et al. (2007), using *OsMADS2* gene-specific RNA probe of ∼644bp cDNA sequences, excluding the MADS domain (Nandi et al., 2000), *OsMADS4* gene-specific RNA probe of 289bp cDNA sequences contain the C terminal and 3’UTR (Yadav et al., 2007) and gene-specific RNA probes for *CYCLIN-P4-1-LIKE*, *PIP1-1* and *PTR2* (Supplementary Table 3).

### Inference of Gene Regulatory Networks

We overlapped 2,589 OsMADS2-bound genes (ChIP-Seq output) with the DEGs (1.5-fold change and FDR ≤ 0.05, RNA-Seq output) from two progressive inflorescence stages (early and middle pools) to identify direct target genes (genes common to ChIP-Seq and RNA-Seq datasets). Transcription factors from these direct targets were extracted and used as first layer nodal genes to link OsMADS2 regulation to its indirectly regulated downstream genes (DEGs that are not present in the ChIP-Seq dataset). For this, promotor analysis was conducted to identify indirect targets with at least a single occurrence of DNA-binding motifs for WRKY8 (TTGACY), ERF65 (AGCCGCC), OsHOX2 (CAAT(G/C)ATTG), OsHOX4 (CAAT(T/A)ATTG), OsARF7A (TGTCTC), OsPCF2 (GGNCCCAC), OsPCF5 (GTGGNCCC), OsSPL2 (TNCGTACAA), and HSFC1B (TTCTAGAA/GAANNTTC). Occurrences of these motifs were checked in 600 bp of assumed promotor sequence (-500 to +100 bp with respect to the TSS). The binding site information was retrieved from the PlantPAN3.0 database (Chow et.al., 2019). Cytoscape software (Shannon et al., 2003) was used to draw the network.

### Statistical Analysis

Student’s *t* tests and Chi-square were performed using the GraphPad Prism software (version 9.0.0). Chi-square was used to test conformity of observed segregation ratio with expected Mendelian ratio. Two-tailed Student’s *t* tests used to compare the differences between two groups. Multiple unpaired Student’s *t* tests were used to compares the means of multiple groups. The number of individuals and replicates is indicated for each experiment either in the figure legends or the materials and methods section. P values <0.05 were considered statistically significant. The results of the statistical analyses are listed in Supplementary Data Set S6.

## Supporting information

Main MS and figures as a word file

Supplementary data (figures and table) as a word file

Supplementary Data Set 1

Supplementary Data Set 2

Supplementary Data Set 3

Supplementary Data Set 4

Supplementary Data Set 5

Supplementary Data Set 6

## Accession numbers

RNA-Seq and ChIP-Seq data obtained in this study are deposited to NCBI Gene Expression Omnibus (https://www.ncbi.nlm.nih.gov/geo/query/acc.cgi) with number GSE260916 and GSE260918 and will become publicly available immediately upon publication. Other public transcriptome data obtained from NCBI Sequence Read Archive (https://www.ncbi.nlm. nih.gov/sra) are for 6-week-old SAMs of rice: SRR5052708 and 9-day-old SAMs of *Arabidopsis*: SRR2073174. The accession numbers, as per RAP database (https://rapdb.dna.affrc.go.jp/), for genes mentioned in results section are as follow: *OsMADS2*: Os01g0883100; *OsMADS4*: Os05g0423400; *GH9B16*: Os08g0114200; *HSFC1B*: Os01g0733200; *PCF2*: Os08g0544800; *PTR2*: Os12g0638200; *CYCLIN-P4-1-LIKE*: Os07g0231500; *EGL1*: Os05g0375400; *YUC1*: Os01g0645400; *YUC3*: Os01g0732700; *YUC7*: Os04g0128900; *TDD1*: Os04g0463500; *NIT1*: Os02g0635200; *GI*: Os01g0182600; *OsMADS50*: Os03g0122600; Os*MADS55*: Os06g0217300; *HOX1*: Os10g0561800; *IIP3*: Os02g0575700; *PME24*: Os08g0220400; *BGLU30*: Os09g0491100; *TPKb*: Os07g0108800; *TIP4-1*: Os05g0231700; *TIP4-3*: Os01g0232000; *PIP1A*: Os02g0666200; *SPL16*: Os08g0531600; *FON1*: Os06g0717200; *TDR*: Os01g0293100; *MTR1*: Os02g0491300; *SPL*: Os05g0501800; *ESP*: Os01g0356951; *GA20ox4*: Os05g0421900; *MADS13*: Os12g0207000; *DFOT1*: Os01g0611000; *HOX12*: Os03g0198600; *JAZ6*: Os03g0402800; *JAZ7*: Os07g0615200; *KRP3*: Os11g0614800.

## Supplementary Data

**aSupplementary Figure S1**. Generation and characterization of CRISPR/Cas9-derived *osmads2* heritable mutant alleles.

**Supplementary Figure S2**. Additional phenotypic analysis of the different genotypes studied.

**Supplementary Figure S3.** Supporting histological analysis of floral organs in different genotypes studied.

**Supplementary Figure S4.** Analysis of pollen viability in the different genotypes studied.

**Supplementary Figure S5.** Expression analysis (fold-change) of the B-class genes *OsMADS2* and *OsMADS4* in *osmads4kd#1* transgenics.

**Supplementary Figure S6.** Analysis of flowering time in different genotypes studied and meta-analysis of expression status of B-class MADS domain genes in rice and *Arabidopsis* shoot apical meristems.

**Supplementary Figure S7.** Characterization of the integration site and illustration of genotyping for zygosity of the *OsMADS4* dsRNAi cassette.

**Supplementary Figure S8.** Scanning electron micrographs (SEM) of developing floral meristems and floret primordia in WT and rice *PI*-clade gene variants *osmads2^d8/d8^* and *osmads2^d8/d8^ osmads4kd*.

**Supplementary Figure S9.** Supporting histological analyses of ovule differentiation in WT and *osmads2 osmads4kd* doubly perturbed florets.

**Supplementary Figure S10.** Supporting analyses for DNA *cis-*regulatory motifs in peak regions bound by OsMADS2 and ChIP-qPCR validation of binding at *EGL1*.

**Supplementary Figure 11**. Supporting gene expression analyses.

**Supplementary Figure S12.** Gene Regulatory Network constructed based on ChIP-Seq and transcriptome data (RNA-Seq, tissue pool constituting 0.2-0.5 cm panicles).

**Supplementary Figure S13.** An OsMADS2 Gene Regulatory Network (GRN) constructed based on ChIP-Seq and transcriptome data (RNA-Seq, tissue pool constituting 0.5-1 cm panicles).

**Supplementary Figure S14.** Supporting spatiotemporal gene expression analysis of OsMADS2 targets and characterization of the WT lodicule.

**Supplementary Table S1:** Analysis of segregation of different *osmads2* mutant alleles using the Chi-square test.

**Supplementary Table S2:** Summary of spikelet phenotyping data for WT and *osmads^d8/d8^*

**Supplementary Table S3:** Summary of genotyping and phenotyping data from the two phenotypic groups of *osmads2 osmads4kd* double mutant plants.

**Supplementary Table S4:** Sequences of the different primers and oligos used in this study

**Supplementary Data Set 1:** ChIP-Seq peaks enriched in an anti-OsMADS2 ChIP experiment.

**Supplementary Data Set 2:** List of DEGs in different *osmads2* data sets.

**Supplementary Data Set 3:** List of Gene Ontology (GO) terms enriched in different *osmads2* DEG subsets

**Supplementary Data Set 4:** List of DEGs in *osmads2* that have orthologs deregulated in the *Arabidopsis pi-1* mutant

**Supplementary Data Set 5:** Supporting analysis of the GRNs. List of DEGs in early and middle *osmads2* expression data sets with respective number of DNA binding motifs for the different nodal transcription factors.

**Supplementary Data Set 6:** Supplementary Data Set 6. Summary of statistical analyses

## Acknowledgments

We acknowledge the DBT-IISc Partnership Divisional Bio-imaging and Greenhouse Facilities. We thank the Functional Genomics Center Zurich for access to its excellent infrastructure to perform next-generation sequencing for the ChIP-Seq experiment. We thank Prof. MS Sheshshayee, University of Agricultural Sciences, GKVK, Bangalore, for providing access to instrument and technical assist for measuring the osmolality. We thank Saurabh Bhatt from UVR laboratory for generating the PTR2 specific clone used for RNA *in situ* hybridization. We acknowledge inputs from members of the UVR laboratory during the course of this study and inputs from Prof. Kalika Prasad, IISER Pune, during the course of manuscript preparation. We also thank Murthy and Jagadeesh for assistance in plant growth and care in field plots and transgenic greenhouses.

## Author contributions

UV, UG and MZ designed the research. MZ, SS^1^, SP, RB, IK, and SS^2^ performed the experiments. RP and MZ performed the bioinformatic analyses. Manuscript writing was done by MZ, UG, and UV with contributions from IK and SS. All authors read and approved the final manuscript.

## References

Abu-Zaitoon YM. Phylogenetic analysis of putative genes involved in the tryptophan-dependent pathway of auxin biosynthesis in rice. Appl Biochem Biotechnol. 2014:172(5):2480–2495. 10.1007/S12010-013-0710-4/FIGURES/7

Ahmad I, Devonshire J, Mohamed R, Schultze M, and Maathuis FJM. Overexpression of the potassium channel TPKb in small vacuoles confers osmotic and drought tolerance to rice. New Phytol. 2016:209(3):1040–1048. 10.1111/NPH.13708

Akiyama T, Jin S, Yoshida M, Hoshino T, Opassiri R, and Ketudat Cairns JR. Expression of an endo-(1,3;1,4)-beta-glucanase in response to wounding, methyl jasmonate, abscisic acid and ethephon in rice seedlings. J Plant Physiol. 2009:166(16):1814–1825. 10.1016/J.JPLPH.2009.06.002

Ambrose BA, Lerner DR, Ciceri P, Padilla CM, Yanofsky MF, and Schmidt RJ. Molecular and Genetic Analyses of the *Silky1* Gene Reveal Conservation in Floral Organ Specification between Eudicots and Monocots. Mol Cell. 2000:5(3):569–579. 10.1016/S1097-2765(00)80450-5

Balázsi G, Van Oudenaarden A, and Collins JJ. Cellular decision making and biological noise: from microbes to mammals. Cell. 2011:144(6):910–925. 10.1016/J.CELL.2011.01.030

Bartlett ME, Williams SK, Taylor Z, Deblasio S, Goldshmidt A, Hall DH, Schmidt RJ, Jackson DP, and Whipplea CJ. The Maize *PI/GLO* Ortholog *Zmm16/sterile tassel silky ear1* Interacts with the Zygomorphy and Sex Determination Pathways in Flower Development. Plant Cell. 2015:27(11):3081–3098. 10.1105/TPC.15.00679

Bowman JL, Smyth DR, and Meyerowitz EM. Genes directing flower development in *Arabidopsis*. Plant Cell. 1989:1(1):37–52. 10.1105/TPC.1.1.37

Busk PK and Pagès M. Microextraction of Nuclear Proteins from Single Maize Embryos. Plant Mol Biol Report. 1997:15(4):371–376. 10.1023/A:1007428802474/METRICS

Cao L, Tian J, Liu Y, Chen X, Li S, Persson S, Lu D, Chen M, Luo Z, Zhang D, et al. Ectopic expression of OsJAZ6, which interacts with OsJAZ1, alters JA signaling and spikelet development in rice. Plant J. 2021:108(4):1083–1096. 10.1111/TPJ.15496

Chen S, Zhou Y, Chen Y, and Gu J. fastp: an ultra-fast all-in-one FASTQ preprocessor. Bioinformatics. 2018:34(17):i884–i890. 10.1093/BIOINFORMATICS/BTY560

Choi MG, Kim EJ, Song JY, Choi SB, Cho SW, Park CS, Kang CS, and Park Y Il. Peptide transporter2 (PTR2) enhances water uptake during early seed germination in *Arabidopsis thaliana*. Plant Mol Biol. 2020:102(6):615–624. 10.1007/s11103-020-00967-3

Chow CN, Lee TY, Hung YC, Li GZ, Tseng KC, Liu YH, Kuo PL, Zheng HQ, and Chang WC. PlantPAN3.0: a new and updated resource for reconstructing transcriptional regulatory networks from ChIP-seq experiments in plants. Nucleic Acids Res. 2019:47(D1):D1155– D1163. 10.1093/NAR/GKY1081

Chung YY, Kim SR, Kang HG, Noh YS, Park MC, Finkel D, and An G. Characterization of two rice MADS box genes homologous to *GLOBOSA*. Plant Sci. 1995:109(1):45–56. 10.1016/0168-9452(95)04153-L

Coen ES and Meyerowitz EM. The war of the whorls: genetic interactions controlling flower development. Nat 1991 3536339. 1991:353(6339):31–37. 10.1038/353031a0

Cosgrove DJ. Enzymes and other agents that enhance cell wall extensibility. Annu Rev Plant Physiol Plant Mol Biol. 1999:50:391–417. 10.1146/ANNUREV.ARPLANT.50.1.391

Dobin A, Davis CA, Schlesinger F, Drenkow J, Zaleski C, Jha S, Batut P, Chaisson M, and Gingeras TR. STAR: ultrafast universal RNA-seq aligner. Bioinformatics. 2013:29(1):15–21. 10.1093/BIOINFORMATICS/BTS635

Doi K, Izawa T, Fuse T, Yamanouchi U, Kubo T, Shimatani Z, Yano M, and Yoshimura A. Ehd1, a B-type response regulator in rice, confers short-day promotion of flowering and controls *FT*-like gene expression independently of Hd1. Genes Dev. 2004:18(8):926. 10.1101/GAD.1189604

Dreni L, Jacchia S, Fornara F, Fornari M, Ouwerkerk PBF, An G, Colombo L, and Kater MM. The D-lineage MADS-box gene *OsMADS13* controls ovule identity in rice. Plant J. 2007:52(4):690–699. 10.1111/J.1365-313X.2007.03272.X

Elowitz MB, Levine AJ, Siggia ED, and Swain PS. Stochastic gene expression in a single cell. Science. 2002:297(5584):1183–1186. 10.1126/SCIENCE.1070919

Ewels P, Magnusson M, Lundin S, and Käller M. MultiQC: summarize analysis results for multiple tools and samples in a single report. Bioinformatics. 2016:32(19):3047–3048. 10.1093/BIOINFORMATICS/BTW354

Favaro R, Pinyopich A, Battaglia R, Kooiker M, Borghi L, Ditta G, Yanofsky MF, Kater MM, and Colombo L. MADS-Box Protein Complexes Control Carpel and Ovule Development in *Arabidopsis*. Plant Cell. 2003:15(11):2603–2611. 10.1105/TPC.015123

Gao S, Fang J, Xu F, Wang W, and Chu C. Rice HOX12 regulates panicle exsertion by directly modulating the expression of ELONGATED UPPERMOST INTERNODE1. Plant Cell. 2016:28(3):680–695. 10.1105/TPC.15.01021

Gómez-Ariza J, Brambilla V, Vicentini G, Landini M, Cerise M, Carrera E, Shrestha R, Chiozzotto R, Galbiati F, Caporali E, et al. A transcription factor coordinating internode elongation and photoperiodic signals in rice. Nat Plants 2019 54. 2019:5(4):358–362. 10.1038/s41477-019-0401-4

Gonçalves B, Nougué O, Jabbour F, Ridel C, Morin H, Laufs P, Manicacci D, and Damerval C. An APETALA3 homolog controls both petal identity and floral meristem patterning in *Nigella damascena* L. (Ranunculaceae). Plant J. 2013:76(2):223–235. 10.1111/TPJ.12284

Hayama R, Yokoi S, Tamaki S, Yano M, and Shimamoto K. Adaptation of photoperiodic control pathways produces short-day flowering in rice. Nat 2003 4226933. 2003:422(6933):719–722. 10.1038/nature01549

Heslop-Harrison Y and Heslop-Harrison JS. Lodicule function and filament extension in the grasses: Potassium ion movement and tissue specialization. Ann Bot. 1996:77(6):573–582. 10.1093/AOB/77.6.573

Hill TA, Day CD, Zondlo SC, Thackeray AG, and Irish VF. Discrete spatial and temporal cis-acting elements regulate transcription of the *Arabidopsis* floral homeotic gene *APETALA3*. Development. 1998:125(9):1711–1721. 10.1242/DEV.125.9.1711

Hu Y, Wang L, Jia R, Liang W, Zhang X, Xu J, Chen X, Lu D, Chen M, Luo Z, et al. Rice transcription factor MADS32 regulates floral patterning through interactions with multiple floral homeotic genes. J Exp Bot. 2021:72(7):2434–2449. 10.1093/JXB/ERAA588

Jack T, Fox GL, and Meyerowitz EM. *Arabidopsis* homeotic gene *APETALA3* ectopic expression: transcriptional and posttranscriptional regulation determine floral organ identity. Cell. 1994:76(4):703–716. 10.1016/0092-8674(94)90509-6

Ji J and Braam J. Restriction Site Extension PCR: A novel method for high-throughput characterization of tagged DNA fragments and genome walking. PLoS One. 2010:5(5):e10577. 10.1371/JOURNAL.PONE.0010577

Joldersma D and Liu Z. The making of virgin fruit: the molecular and genetic basis of parthenocarpy. J Exp Bot. 2018:69(5):955–962. 10.1093/JXB/ERX446

Kang HG, Jeon JS, Lee S, and An G. Identification of class B and class C floral organ identity genes from rice plants. Plant Mol Biol. 1998:38(6):1021–1029. 10.1023/A:1006051911291/METRICS

Kaufmann K, Muiño JM, Jauregui R, Airoldi CA, Smaczniak C, Krajewski P, and Angenent GC. Target genes of the MADS transcription factor SEPALLATA3: integration of developmental and hormonal pathways in the *Arabidopsis* flower. PLoS Biol. 2009:7(4):0854– 0875. 10.1371/JOURNAL.PBIO.1000090

Klepikova A V., Logacheva MD, Dmitriev SE, and Penin AA. RNA-seq analysis of an apical meristem time series reveals a critical point in *Arabidopsis thaliana* flower initiation. BMC Genomics. 2015:16(1):1–16. doi:10.1186/s12864-015-1688-9

Krizek BA and Fletcher JC. Molecular mechanisms of flower development: an armchair guide. Nat Rev Genet 2005 69. 2005:6(9):688–698. 10.1038/nrg1675

Krizek BA and Meyerowitz EM. The *Arabidopsis* homeotic genes *APETALA3* and *PISTILLATA* are sufficient to provide the B class organ identity function. Development. 1996:122(1):11–22. 10.1242/DEV.122.1.11

Krizek BA. Auxin regulation of Arabidopsis flower development involves members of the AINTEGUMENTA-LIKE/PLETHORA (AIL/PLT) family. J Exp Bot. 2011:62(10):3311–3319. 10.1093/JXB/ERR127

Lee JH, Park SH, and Ahn JH. Functional conservation and diversification between rice OsMADS22/OsMADS55 and *Arabidopsis* SVP proteins. Plant Sci. 2012:185–186:97–104. 10.1016/J.PLANTSCI.2011.09.003

Lee S, Jeon JS, An K, Moon YH, Lee S, Chung YY, and An G. Alteration of floral organ identity in rice through ectopic expression of *OsMADS16*. Planta. 2003:217(6):904–911. 10.1007/S00425-003-1066-8

Li N, Zhang DS, Liu HS, Yin CS, Li XX, Liang WQ, Yuan Z, Xu B, Chu HW, Wang J, et al. The rice *tapetum degeneration retardation* gene is required for tapetum degradation and anther development. Plant Cell. 2006:18(11):2999–3014. 10.1105/TPC.106.044107

Li Q, Tong T, Jiang W, Cheng J, Deng F, Wu X, Chen ZH, Ouyang Y, and Zeng F. Highly Conserved Evolution of Aquaporin PIPs and TIPs Confers Their Crucial Contribution to Flowering Process in Plants. Front Plant Sci. 2022:12:2945. 10.3389/FPLS.2021.761713/BIBTEX

Li S, Wei K, Zhang L, Ning Y, Lu F, Wang X, Guo Y, Liu L, Li X, Zhu C, et al. Fine mapping and candidate gene validation of tomato gene *Carpelloid Stamen and Parthenocarpy* (*CSP*). Horticulturae. 2024:10(4):403. 10.3390/HORTICULTURAE10040403/S1

Li Y, Ouyang J, Wang YY, Hu R, Xia K, Duan J, Wang Y, Tsay YF, and Zhang M. Disruption of the rice nitrate transporter OsNPF2.2 hinders root-to-shoot nitrate transport and vascular development. Sci Reports 2015 51. 2015:5(1):1–10. 10.1038/srep09635

Liang M, Deng L, Liu J, He A, and Chen L. Interaction between the *eui* gene and thermo-sensitive genic male sterility in rice. Euphytica. 2008:164(3):637–643. 10.1007/S10681-008-9657-X/TABLES/2

Liao Y, Smyth GK, and Shi W. The R package Rsubread is easier, faster, cheaper and better for alignment and quantification of RNA sequencing reads. Nucleic Acids Res. 2019:47(8):e47–e47. 10.1093/NAR/GKZ114

Lohmann JU and Weigel D. Building beauty: the genetic control of floral patterning. Dev Cell. 2002:2(2):135–142. 10.1016/S1534-5807(02)00122-3

Luan X, Liu S, Ke S, Dai H, Xie XM, Hsieh TF, and Zhang XQ. Epigenetic modification of *ESP*, encoding a putative long noncoding RNA, affects panicle architecture in rice. Rice. 2019:12(1). 10.1186/S12284-019-0282-1

Machanick P and Bailey TL. MEME-ChIP: motif analysis of large DNA datasets. Bioinformatics. 2011:27(12):1696–1697. 10.1093/BIOINFORMATICS/BTR189

Mizutani M, Naganuma T, Tsutsumi KI, and Saitoh Y. The syncytium-specific expression of the Orysa;KRP3 CDK inhibitor: implication of its involvement in the cell cycle control in the rice (*Oryza sativa* L.) syncytial endosperm. J Exp Bot. 2010:61(3):791–798. 10.1093/JXB/ERP343

Nagasawa N, Miyoshi M, Sano Y, Satoh H, Hirano H, Sakai H, and Nagato Y. *SUPERWOMAN1* and *DROOPING LEAF* genes control floral organ identity in rice. Development. 2003:130(4):705–718. 10.1242/DEV.00294

Nandi Ak, Kushalappa K, Prasad K, and Vijayraghavan U. A conserved function for *Arabidopsis SUPERMAN* in regulating floral-whorl cell proliferation in rice, a monocotyledonous plant. Curr Biol. 2000:10(4):215–218. 10.1016/S0960-9822(00)00341-9

Ning J, Zhang B, Wang N, Zhou Y, and Xiong L. Increased Leaf Angle1, a Raf-Like MAPKKK that interacts with a nuclear protein family, regulates mechanical tissue formation in the lamina joint of rice. Plant Cell. 2011:23(12):4334–4347. 10.1105/TPC.111.093419

O’Toole JC and Namuco OS. Role of panicle exsertion in water stress induced sterility. Crop Sci. 1983:23(6):1093–1097. 10.2135/CROPSCI1983.0011183X002300060017X

Pagès H, Aboyoun P, Gentleman R, and DebRoy S. Biostrings: Efficient manipulation of biological strings. R package version 2.64.0, (2022). https://bioconductor.org/packages/Biostrings

Pinyopich A, Ditta GS, Savidge B, Liljegren SJ, Baumann E, Wisman E, and Yanofsky MF. Assessing the redundancy of MADS-box genes during carpel and ovule development. Nat 2003 4246944. 2003:424(6944):85–88. 10.1038/nature01741

Prakash S, Rai R, Zamzam M, Ahmad O, Peesapati R, and Vijayraghavan U. OsbZIP47 is an integrator for meristem regulators during rice plant growth and development. Front Plant Sci. 2022:13:883. 10.3389/FPLS.2022.865928/BIBTEX

Prasad K and Vijayraghavan U. Double-stranded RNA interference of a rice *PI/GLO* paralog, *OsMADS2*, uncovers its second-whorl-specific function in floral organ patterning. Genetics. 2003:165(4):2301–2305. 10.1093/GENETICS/165.4.2301

Ren L, Tang D, Zhao T, Zhang F, Liu C, Xue Z, Shi W, Du G, Shen Y, Li Y, et al. OsSPL regulates meiotic fate acquisition in rice. New Phytol. 2018:218(2):789–803. 10.1111/NPH.15017

Robinson MD, McCarthy DJ, and Smyth GK. edgeR: a Bioconductor package for differential expression analysis of digital gene expression data. Bioinformatics. 2010:26(1):139–140. 10.1093/BIOINFORMATICS/BTP616

Ryu CH, Lee S, Cho LH, Kim SL, Lee YS, Choi SC, Jeong HJ, Yi J, Park SJ, Han CD, et al. OsMADS50 and OsMADS56 function antagonistically in regulating long day (LD)- dependent flowering in rice. Plant Cell Environ. 2009:32(10):1412–1427. 10.1111/J.1365-3040.2009.02008.X

Scarpella E, Rueb S, Boot KJM, Hoge JHC, and Meijer AH. A role for the rice homeobox gene *Oshox1* in provascular cell fate commitment. Development. 2000:127(17):3655–3669. 10.1242/DEV.127.17.3655

Schmidt R, Schippers JHM, Welker A, Mieulet D, Guiderdoni E, and Mueller-Roeber B. Transcription factor OsHsfC1b regulates salt tolerance and development in *Oryza sativa* ssp. japonica. AoB Plants. 2012:2012(1). 10.1093/AOBPLA/PLS011

Selva C, Yang X, Shirley NJ, Whitford R, Baumann U, and Tucker MR. HvSL1 and HvMADS16 promote stamen identity to restrict multiple ovary formation in barley. J Exp Bot. 2023:74(17):5039–5056. 10.1093/JXB/ERAD218

Shannon P, Markiel A, Ozier O, Baliga NS, Wang JT, Ramage D, Amin N, Schwikowski B, and Ideker T. Cytoscape: a software environment for integrated models of biomolecular interaction networks. Genome Res. 2003:13(11):2498–2504. 10.1101/GR.1239303

Shi J, Hou F, Dong Y, Pan Y, Zhou Q, Zhang Z, Liu Y, and Liu W. Functional validation of two B-class MADS-box genes *HmPI* and *HmAP3* from *Hemerocallis middendorffii*. Vitr Cell Dev Biol - Plant. 2024:1–12. 10.1007/s11627-024-10470-9

Sazuka T, Kamiya N, Nishimura T, Ohmae K, Sato Y, Imamura K, Nagato Y, Koshiba T, Nagamura Y, Ashikari M, et al. A *rice tryptophan deficient dwarf mutant*, *tdd1*, contains a reduced level of indole acetic acid and develops abnormal flowers and organless embryos. Plant J. 2009:60(2):227–241. 10.1111/J.1365-313X.2009.03952.X

Suzaki T, Sato M, Ashikari M, Miyoshi M, Nagato Y, and Hirano HY. The gene *FLORAL ORGAN NUMBER1* regulates floral meristem size in rice and encodes a leucine-rich repeat receptor kinase orthologous to *Arabidopsis* CLAVATA1. Development. 2004:131(22):5649– 5657. 10.1242/DEV.01441

Tan H, Liang W, Hu J, and Zhang D. *MTR1* encodes a secretory fasciclin glycoprotein required for male reproductive development in rice. Dev Cell. 2012:22(6):1127–1137. 10.1016/J.DEVCEL.2012.04.011

Veronico P, Rosso LC, Melillo MT, Fanelli E, De Luca F, Ciancio A, Colagiero M, and Pentimone I. Water stress differentially modulates the expression of tomato cell wall metabolism-related genes in *Meloidogyne incognita* feeding sites. Front Plant Sci. 2022:13:776. 10.3389/FPLS.2022.817185/BIBTEX

Wang H, Zhang L, Cai Q, Hu Y, Jin Z, Zhao X, Fan W, Huang Q, Luo Z, Chen M, et al. OsMADS32 interacts with PI-like proteins and regulates rice flower development. J Integr Plant Biol. 2015:57(5):504–513. 10.1111/JIPB.12248

Wang L, Li Q-L, Hu J-P, Yuan Z, Wang L, Li Q-L, Hu J-P, and Yuan Z. Neofunctionalization of B-class genes in regulating rice flower development. Seed Biol 2024 1e013. 2024a:3(1). 10.48130/SEEDBIO-0024-0012

Wang M, Zhu X, Huang Z, Chen M, Xu P, Liao S, Zhao Y, Gao Y, He J, Luo Y, et al. Controlling diurnal flower-opening time by manipulating the jasmonate pathway accelerates development of indica–japonica hybrid rice breeding. Plant Biotechnol J. 2024b:22(8):2267– 2281. 10.1111/PBI.14343

Wang M, Zhu X, Peng G, Liu M, Zhang S, Chen M, Liao S, Wei X, Xu P, Tan X, et al. Methylesterification of cell-wall pectin controls the diurnal flower-opening times in rice. Mol Plant. 2022:15(6):956–972. 10.1016/J.MOLP.2022.04.004

Wang S, Wu K, Yuan Q, Liu X, Liu Z, Lin X, Zeng R, Zhu H, Dong G, Qian Q, et al. Control of grain size, shape and quality by OsSPL16 in rice. Nat Genet 2012 448. 2012:44(8):950–954. 10.1038/ng.2327

Winter CM, Austin RS, Blanvillain-Baufumé S, Reback MA, Monniaux M, Wu MF, Sang Y, Yamaguchi A, Yamaguchi N, Parker JE, et al. LEAFY target genes reveal floral regulatory logic, cis motifs, and a link to biotic stimulus response. Dev Cell. 2011:20(4):430–443. 10.1016/J.DEVCEL.2011.03.019

Wuest SE, O’Maoileidigh DS, Rae L, Kwasniewska K, Raganelli A, Hanczaryk K, Lohan AJ, Loftus B, Graciet E, and Wellmer F. Molecular basis for the specification of floral organs by APETALA3 and PISTILLATA. Proc Natl Acad Sci U S A. 2012:109(33):13452–13457. 10.1073/PNAS.1207075109/SUPPL_FILE/SD04.XLSX

Xie G, Yang B, Xu Z, Li F, Guo K, Zhang M, Wang L, Zou W, Wang Y, and Peng L. Global identification of multiple OsGH9 family members and their involvement in cellulose crystallinity modification in rice. PLoS One. 2013:8(1):50171. 10.1371/JOURNAL.PONE.0050171

Xu P, Wu T, Ali A, Zhang H, Liao Y, Chen X, Tian Y, Wang W, Fu X, Li Y, et al. EARLY MORNING FLOWERING1 (EMF1) regulates the floret opening time by mediating lodicule cell wall formation in rice. Plant Biotechnol J. 2022:20(8):1441–1443. 10.1111/PBI.13860

Yadav SR, Prasad K, and Vijayraghavan U. Divergent regulatory OsMADS2 functions control size, shape and differentiation of the highly derived rice floret second-whorl organ. Genetics. 2007:176(1):283–294. 10.1534/GENETICS.107.071746

Yang Y, Xiang H, and Jack T. *pistillata-5*, an *Arabidopsis* B class mutant with strong defects in petal but not in stamen development. Plant J. 2003:33(1):177–188. 10.1046/J.1365-313X.2003.01603.X

Yao JL, Dong YH, and Morris BAM. Parthenocarpic apple fruit production conferred by transposon insertion mutations in a MADS-box transcription factor. Proc Natl Acad Sci U S A. 2001:98(3):1306–1311. 10.1073/PNAS.98.3.1306

Yao SG, Ohmori S, Kimizu M, and Yoshida H. Unequal genetic redundancy of rice PISTILLATA orthologs, OsMADS2 and OsMADS4, in lodicule and stamen development. Plant Cell Physiol. 2008:49(5):853–857. 10.1093/PCP/PCN050 10.1073/PNAS.0907896106/SUPPL_FILE/0907896106SI.PDF

Yoshida H, Itoh JI, Ohmori S, Miyoshi K, Horigome A, Uchida E, Kimizu M, Matsumura Y, Kusaba M, Satoh H, et al. *superwoman1*-cleistogamy, a hopeful allele for gene containment in GM rice. Plant Biotechnol J. 2007:5(6):835–846. 10.1111/J.1467-7652.2007.00291.X

Yu G, Wang LG, and He QY. ChIPseeker: an R/Bioconductor package for ChIP peak annotation, comparison and visualization. Bioinformatics. 2015:31(14):2382–2383. 10.1093/BIOINFORMATICS/BTV145

Yun D, Liang W, Dreni L, Yin C, Zhou Z, Kater MM, and Zhang D. *OsMADS16* genetically interacts with *OsMADS3* and *OsMADS58* in specifying. Mol Plant. 2013:6(3):743– 756. 10.1093/mp/sst003

Zeng YX, Hu CY, Lu YG, Li JQ, and Liu XD. Diversity of abnormal embryo sacs in indica/japonica hybrids in rice demonstrated by confocal microscopy of ovaries. Plant Breed. 2007:126(6):574–580. 10.1111/J.1439-0523.2007.01380.X

Zhang Y, Liu T, Meyer CA, Eeckhoute J, Johnson DS, Bernstein BE, Nussbaum C, Myers RM, Brown M, Li W, et al. Model-based analysis of ChIP-Seq (MACS). Genome Biol. 2008:9(9):1–9. 10.1186/gb-2008-9-9-r137

Zhao ZX, Yin XX, Li S, Peng YT, Yan XL, Chen C, Hassan B, Zhou SX, Pu M, Zhao JH, et al. miR167d-ARFs module regulates flower opening and stigma size in rice. Rice. 2022:15(1):1–15. 10.1186/s12284-022-00587-z

Zhu LJ, Gazin C, Lawson ND, Pagès H, Lin SM, Lapointe DS, and Green MR. ChIPpeakAnno: A Bioconductor package to annotate ChIP-seq and ChIP-chip data. BMC Bioinformatics. 2010:11(1):1–10. 10.1186/1471-2105-11-237/TABLES/2

