## Supplementary for "Functional genetics of rice *PISTILLATA* genes unravels new roles and targets in flowering time, female fertility and parthenocarpy"

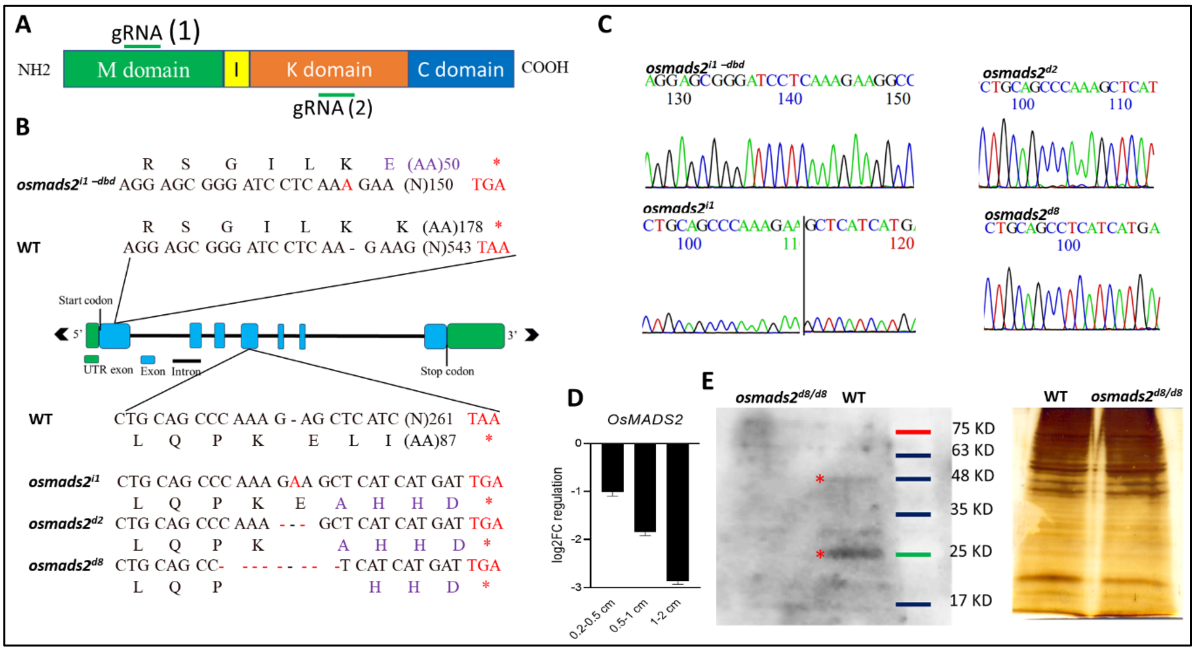


**Supplementary Figure S1**. **Generation and characterization of CRISPR/Cas9-derived *osmads2* heritable mutant alleles**. **(Supports Figure 1).** A, Schematic representation of OsMADS2 protein (209 amino acids) showing MADS-domain between amino acids 2-80 and K-domain between amino acids 83-156. The position of the two gRNAs (1 and 2) for CRISPR-Cas9 gene editing are marked by green horizontal lines. B, Schematic representation showing positions of the different edited mutant alleles with respect to *OsMADS2* genomic locus, the nature of editing and the predicted encoded protein sequences with respect to the WT allele. All mutant alleles are predicted to generate truncated proteins due to the introduction of premature stop codons in their mRNAs. C, Sanger sequencing chromatogram representations of the different mutant alleles. D, qPCR depicting fold change in the abundance of *OsMADS2* transcripts in inflorescences of lengths 0.2-0.5 cm, 0.5-1 cm and 1-2 cm comparing WT to *osmads2*$^{d8/d8}$. Values are means ± the standard error of the mean (SEM). E, Western blot (left panel) performed with OsMADS2 specific antibodies on nuclear lysate from inflorescence tissues. No detectable OsMADS2 monomer/dimers (red asterisk) in *osmads2*$^{d8/d8}$ tissue lysates while OsMADS2 protein and a predicted dimeric form is clearly detected in WT. Right panel is the silver-stained gel demonstrating equal loading of nuclear protein lysates.

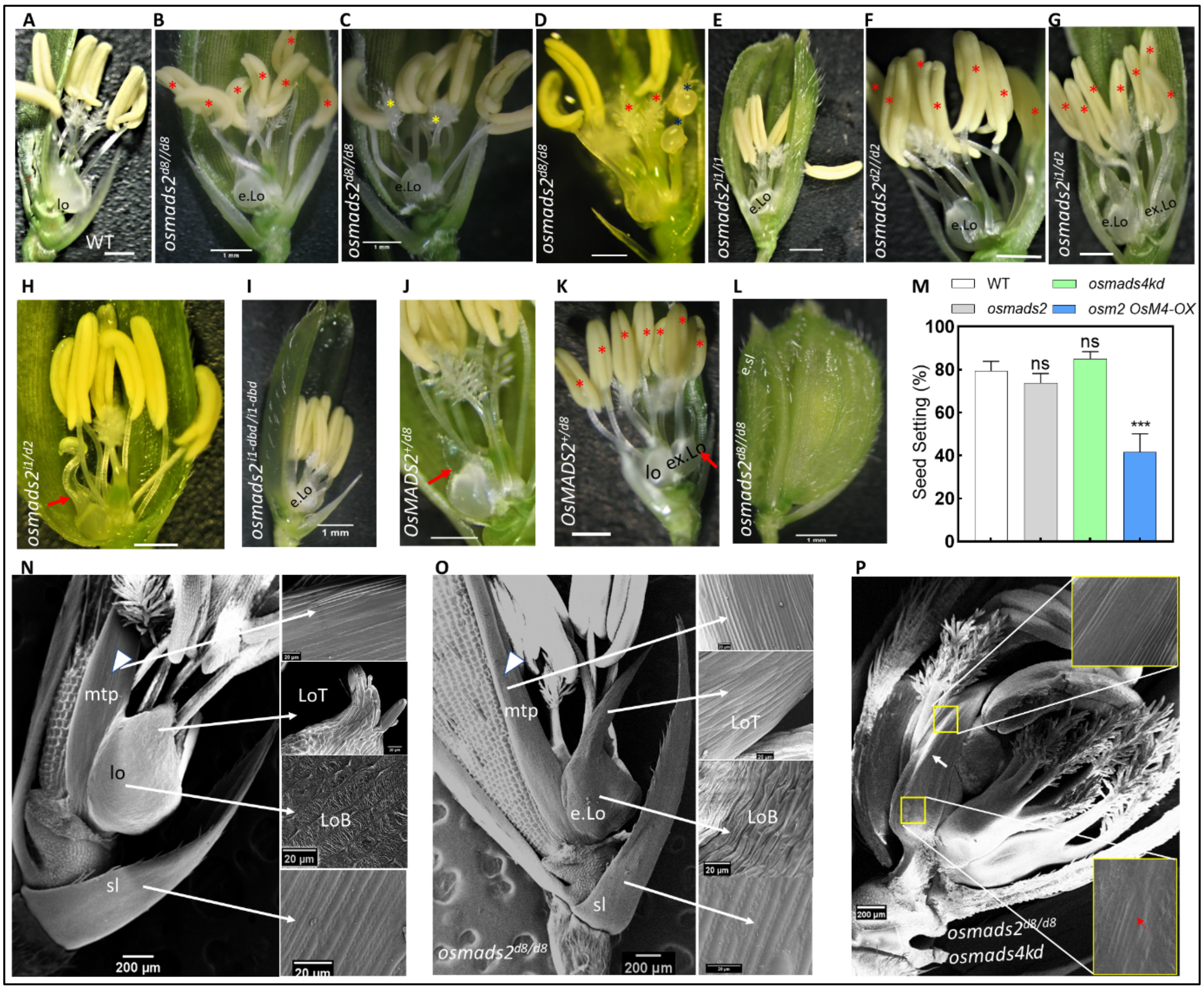


**Supplementary Figure S2**. **Additional phenotypic analysis of the different genotypes studied. (Supports Figure 1).** In A-K and N-P, the lemma and palea were partially dissected to expose the internal organs. A, A WT spikelet with normal floral organs. B, An *osmads2*$^{d8/d8}$ spikelet with elongated lodicules and increased stamen number (red asterisks). C, An *osmads2*$^{d8/d8}$ spikelet with elongated lodicules and two carpels (yellow asterisks). D, An *osmads2*$^{d8/d8}$ spikelet with ectopic carpels (replacing the stamens, red asterisks). The dark blue asterisks mark the stamen-carpel chimeric organs in this spikelet that resemble partially *spw*-like. E-I, Spikelets with different allelic combinations mutant in of OsMADS2. The red asterisks in F and G, mark the increased stamen number. The red arrow in H marks the chimeric lodicule-stamen. J, A heterozygous *osmads2*$^{+/d8}$ spikelet having slightly elongated lodicule. K, A heterozygous *osmads2*$^{+/d8}$ spikelet having extra lodicule and increased stamen number. L, An *osmads2*$^{d8/d8}$ spikelet with the inner sterile lemma being transformed into an elongated organ, a phenotype noted in 3/222 florets, while noted in 44/765 florets in *osmads2*$^{d8/d8}$ *osmads4kd* as compared to 1/508 florets in WT (WT *vs* *osmads2*$^{d8/d8}$ *osmads4kd*, P value < 0.0001, Fisher's exact test). M, Bar graph of seed setting percentage in the different genotypes studied. Values are means ± SEM, P < 0.01 denoted as **; Student's *t* test. N-P, Spikelets from WT, *osmads2*$^{d8/d8}$ and *osmads2*$^{d8/d8}$ *osmads4kd* plants. Open arrowhead points to the marginal tissue of palea (mtp) in N and O. The elongated cells of the epidermal layer of the mutant lodicules phenocopy the cells of the epidermal layers in mtp and in the sterile lemma/sl (Compare the magnified micrographs of lodicule in O and P *vs*. the magnified micrographs of the mtp and of the sl in N and O). Scale bars: 1 mm in A-L, and 200 µm in N-P. Abbreviations: elongated lodicule (e.Lo), extra lodicule (ex.Lo), elongated sterile lemma (e.sl).

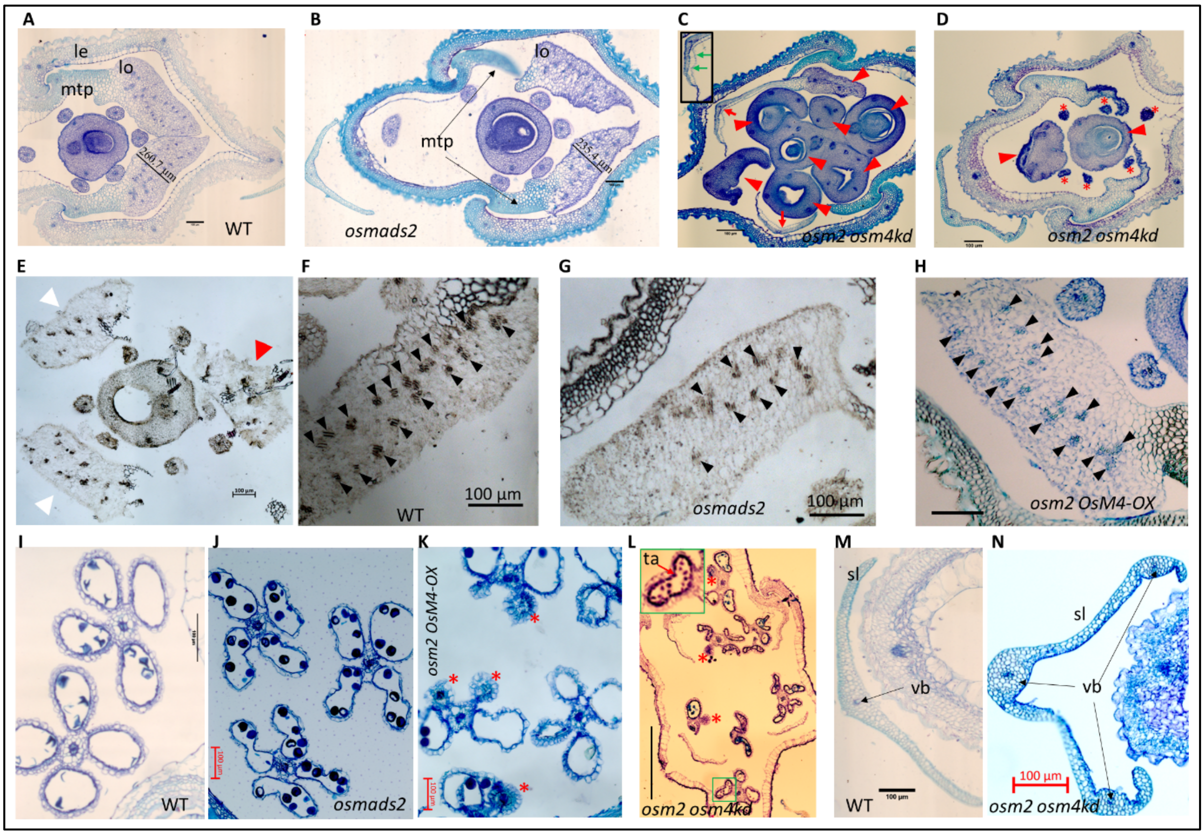


**Supplementary Figure S3. Supporting histological analysis of floral organs in different genotypes studied. (Supports Figure 1).** A and B, Transverse section (TS) at the basal region of WT and *osmads2*$^{d8/d8}$ florets, respectively. The black box marks the peripheral regions of the lodicule that underlie the lemma which are overgrown in the transformed lodicules as compared to WT lodicules. The black lines and numbers denote the width of the lodicule across the abaxial-adaxial axis. C, TS in double mutant *osmads2*$^{d8/d8}$ *osmads4kd* spikelet with phenotype group I. Red arrows point to the midvein in the abnormal lodicule. The lodicule is stained green as for organs with lignified cell walls. The inset shows that the adaxial side of the abnormal lodicule is lined with bubble-shaped cells resembling those lining the lemma and palea. D, TS of double mutant *osmads2*$^{d8/d8}$ *osmads4kd* spikelet with group II phenotype. The internal floral organ number is increased in C and D. The stamens in D are indicated by asterisks and the red arrowheads in C and D point towards the central and ectopic carpels. The abnormal lodicule is stained green as of the mtp. The sterile lemma is abnormal. E, TS of an *osmads2* mutant floret where the extra second whorl lodicule is marked by a red triangle in the second whorl and is clearly positioned external to the stamen whorl. F, TS of WT floret showing internal tissue organization in a mature pre-anthesis WT lodicule stained with Phloroglucinol. The vascular bundles (black arrowheads) are arranged into two parallel rows. G, TS of a lodicule from *osmads2* mutant. Vascular bundle (black arrowheads) number is reduced overall. H, TS of lodicule from an *osmads2 OsMADS4-OX* floret showing rescue of vascular bundle arrangement to the normal pattern and number (black arrowheads). I-L, Transverse sections of anthers in WT, *osmads2, osmads2 OsMADS4-OX*, and *osmads2 osmads4kd* florets. The red asterisks (K and L) mark the underdeveloped anthers. The red arrow in L (inset) marks the tapetum layers which fail to undergo programmed cell death. M and N, Sections of the sterile lemma. The sterile lemma in *osmads2*$^{d8/d8}$ *osmads4kd* florets (N) is abnormal as compared to WT (M). Scale bars are of 100 µm. Abbreviation: le; lemma, pa; palea, lo; lodicule, mtp; marginal tissue of palea, vp; vascular bundles, gl; glumes, ovl; ovule-like, ta; tapetum.

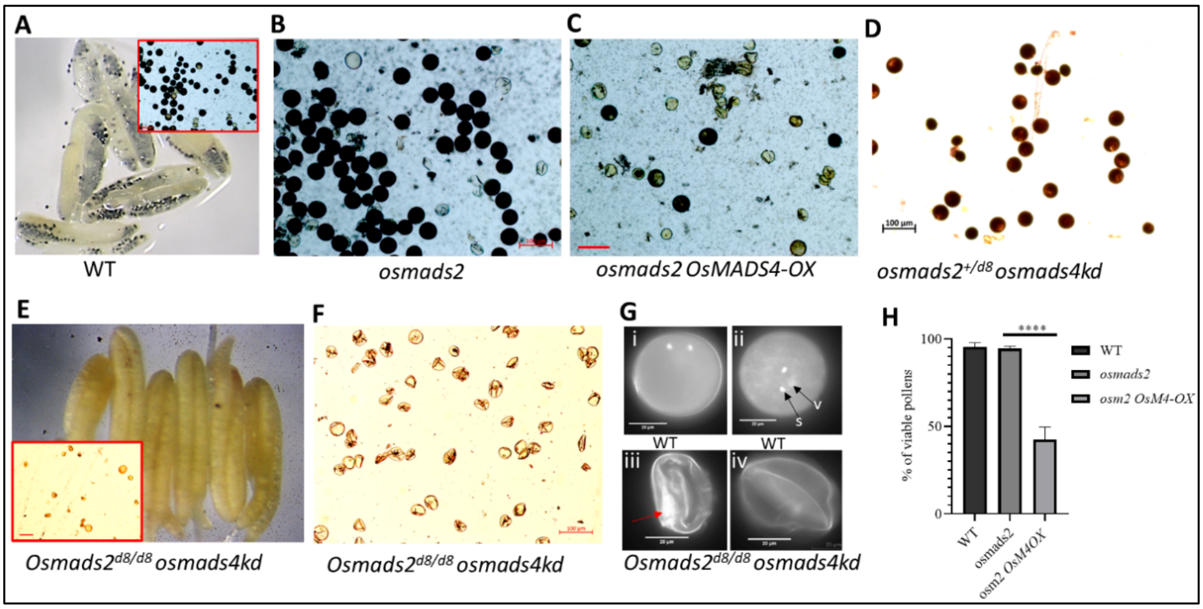


**Supplementary Figure S4. Analysis of pollen viability in the different genotypes studied. (Supports Figure 1).** A-D, Iodine potassium iodide (IKI2) staining for assays of *in vitro* pollen viability. Pollens (Inset) and anthers from WT (A), pollens from *osmads2* mutant (B), and *osmads2*$^{+/d8}$ *osmads4kd* (D) have taken up the dye and are darkly stained. In *osmads2 OsMADS4-OX* (C), there is a significant loss in pollen viability as dye uptake is poor and pollen grains are shrunk. In *osmads*$^{d8/d8}$ *osmads4kd* double mutant (E and F), anthers and pollens are not stained and there is a decreased number of gametes per microscopic field. In F, the microscopic field shows inviable pollens collected from multiple anthers. All images of pollen assay in A-F panels are of the same magnification and a representative scale bar of 100 µm. g, Pollen staining with DAPI showing spherical binucleate (i) and trinucleate (ii) pollens in WT and shrunken pollens (iii and iv) with abnormal nuclear signal (red arrow) in *osmads*$^{d8/d8}$ *osmads4kd* double mutant. S; sperm nuclei, V; vegetative nuclei. H, Graphical plot representing *in vitro* pollen viability in different genotypes. Values are means ± SEM. $P < 0.0001$ denoted as ****; Student's *t* test.

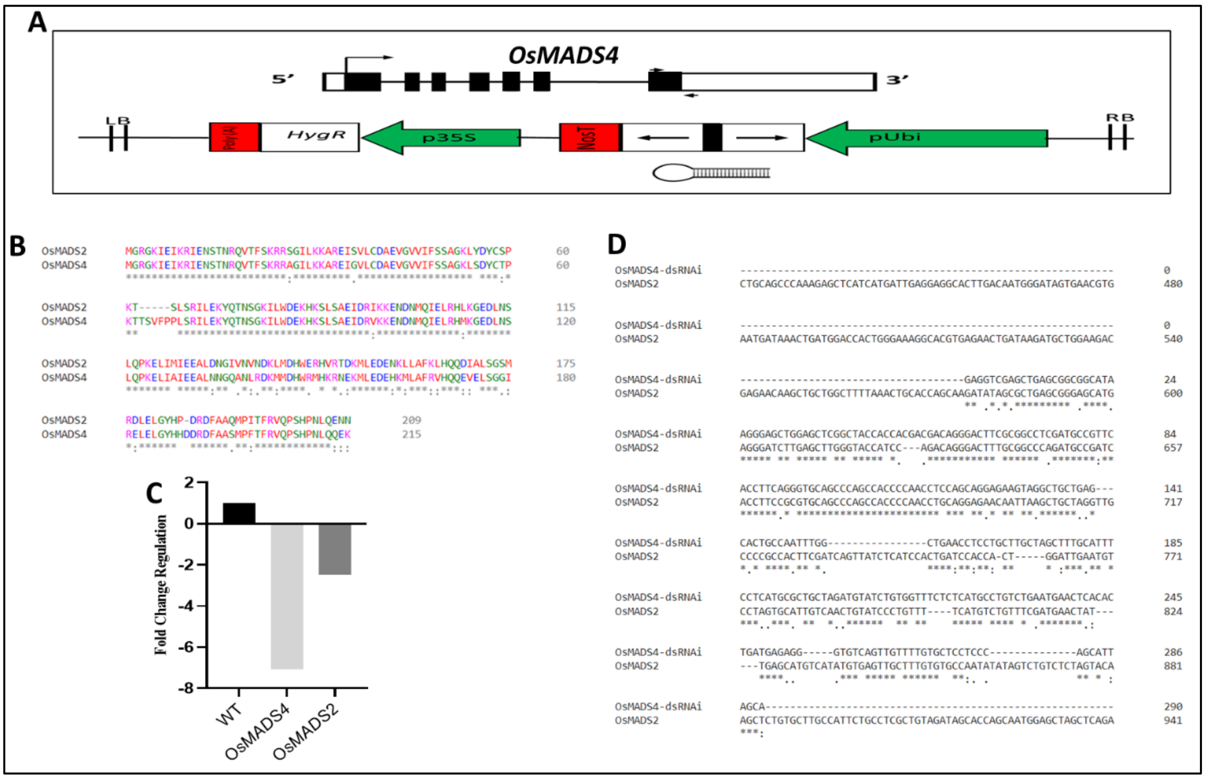


**Supplementary Figure S5. Expression analysis (fold-change) of the B-class genes *OsMADS2* and *OsMADS4* in *osmads4kd#1* transgenics. (Supports Figure 1).** A, Schematic representation of *OsMADS4* locus and the T-DNA segment in the rice binary expression vector used to generate the transgenic *osmads4kd#1*. The black arrows on *OsMADS4* locus schematic indicate the region amplified to construct the hairpin dsRNAi for knockdown in transgenics. B, Amino acid sequence alignment of OsMADS2 and OsMADS4. C, RT-qPCR shows relative expression levels of *OsMADS4* and *OsMADS2* in the *osmads4kd#1 vs.* WT inflorescences. D, Global alignment of *OsMADS2* transcript against the 290 bp *OsMADS4* specific region utilized to construct the hairpin loop for dsRNAi-mediated silencing of *OsMADS4*. The alignment shows that there is no stretch of identical 24 nucleotides.

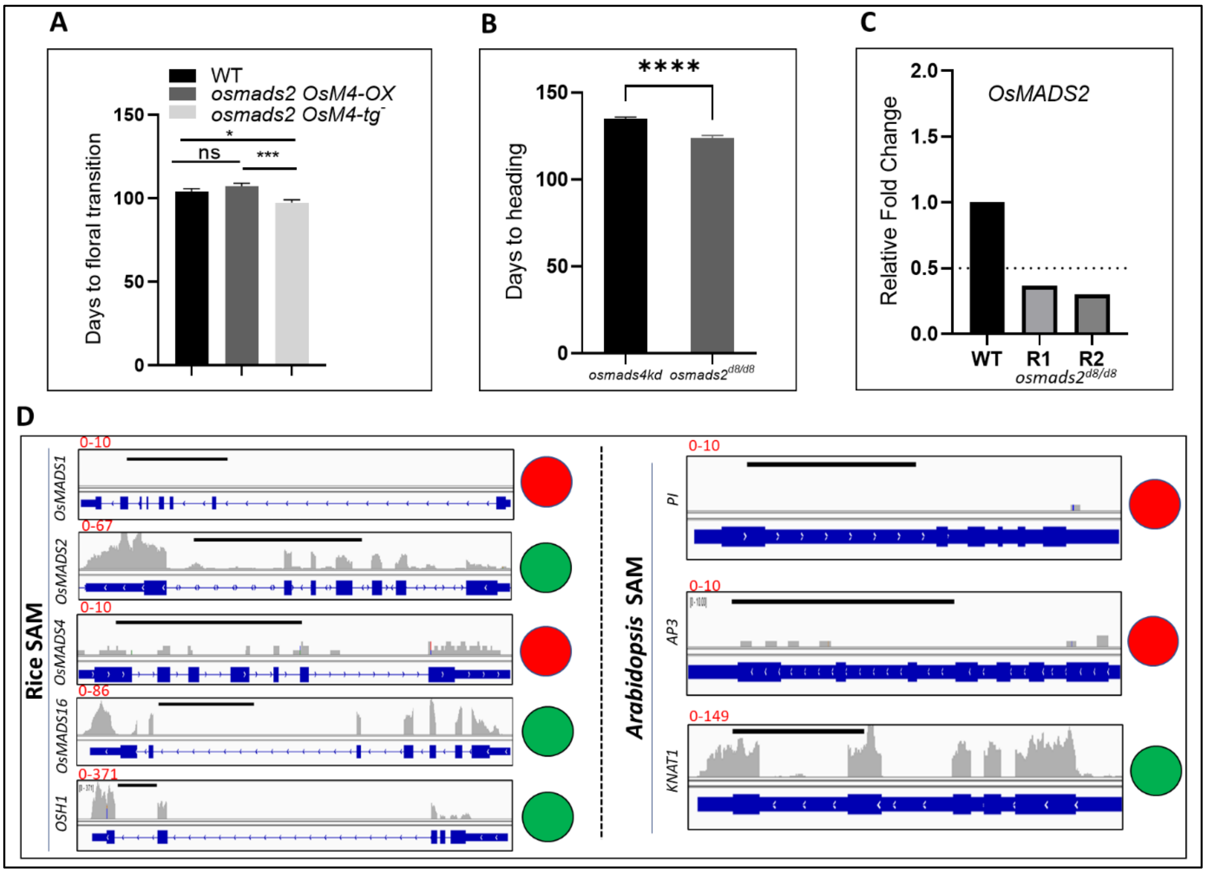


**Supplementary Figure S6. Analysis of flowering time in different genotypes studied and meta-analysis of expression status of B-class MADS domain genes in rice and *Arabidopsis* shoot apical meristems**. **(Supports Figure 1).** A, A bar graph of days taken to floral transition in WT (n=7), *osmads2 OsMADS4-OX* (n=14) and *osmads2 OsMADS4-tg*$^-$ segregants (n=13). B, Bar graph of days taken to panicle heading (emergence from flag leaf) in *osmads4kd vs. osmads2*$^{d8/d8}$ plants (n=34 and 33, respectively). In A and B, values are mean ± SEM. $P < 0.05$, $P < 0.001$ and $P < 0.0001$ denoted as *, *** and ****, respectively. C, A bar graph depicting nearly ~ 3-fold downregulation of *osmads2* mutant transcripts in two biological replicates of 5-7 days post germination SAM tissues. D, Integrated Genome Browser views showing gene expression in SAMs from 6-week-old rice plants and 9 days post germination *Arabidopsis* plants. No expression of *OsMADS1* (negative control) and *OsMADS4*, contrasts with transcripts clearly detected from *OsMADS2*, *OsMADS16*, *OSH1* (a meristem marker, positive control) in SAMs of 6-week-old rice plants. The SAMs in 9 days post germination *Arabidopsis* plants express *KNAT1* (AT4G08150), a member of class I *knotted1-like* homeobox gene family (meristem marker, positive control) but do not express *PI* or *AP3* Class B genes. Green filled circles and red filled circles denoted expressed and not expressed genes, respectively. Information on sources of the utilized available RNA-Seq raw files and mapping are described in materials and method section.

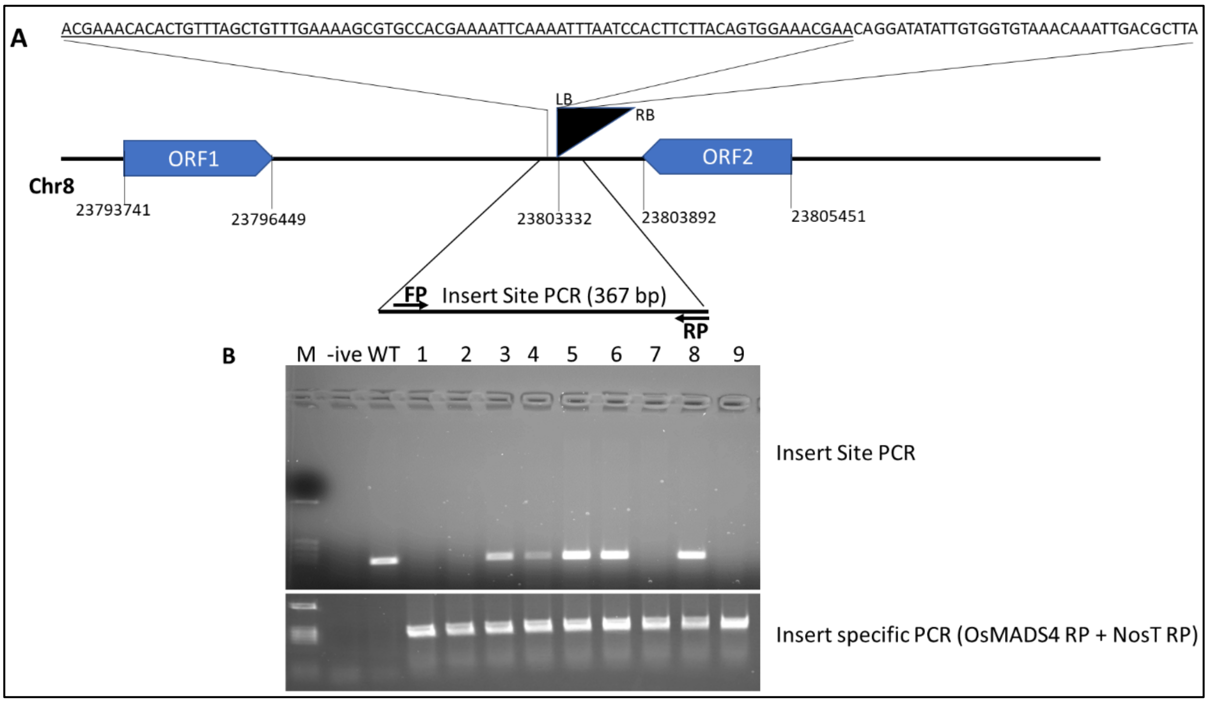


**Supplementary Figure S7. Characterization of the integration site and illustration of genotyping for zygosity of the *OsMADS4* dsRNAi transgenic cassette**. **(Supports Figure 1).** A, Diagram shows the T-DNA insertion site in the rice genome of transgenic rice *osmads4kd#1*. The black filled triangle indicates the T-DNA insertion. The underlined sequence is the genomic DNA flanking the T-DNA left border (LB). the genome coordinates are as per IRGSP-1.0 genome assembly. The black arrows on the magnified genomic region represents the forward and the reverse primers utilized to screen this insertion site for zygosity of the integrated T-DNA. B, Genotyping data of some *osmads2 osmads4kd* double mutant segregants (Lane 1-9). The upper panel is an insert site PCR and the lower panel is an insert specific PCR where the transcriptionally fused OsMADS4 dsRNAi-NOS terminator sequence is amplified. M is a DNA size marker, -ive is a no-DNA control. Samples that are positive to the insert specific PCR and negative to the insert site PCR are homozygous for the integrated T-DNA.

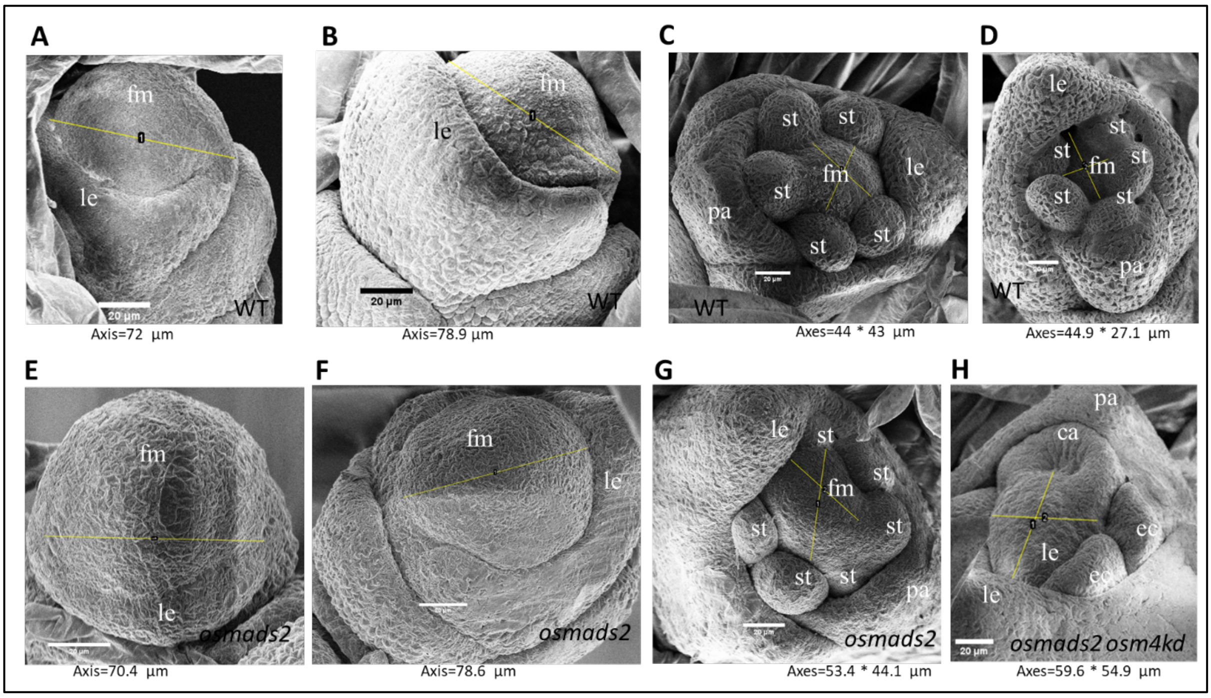


**Supplementary Figure S8. Scanning electron micrographs of developing floral meristems and floret primordia in WT and rice *PI*-clade gene variants *osmads2*$^{d8/d8}$ and *osmads2*$^{d8/d8}$ *osmads4kd*. (Supports Figure 1).** A-D, Developing WT spikelets. E-G, Developing spikelets from *osmads2*$^{d8/d8}$ mutants. H, A spikelet from the double mutant *osmads2*$^{d8/d8}$ *osmads4kd*. Scale bars of 20 µm are indicated in each panel by a white/ black bar. The thin yellow lines give dimensions of FM and these axis values are displayed below each panel. Compare panels C ,D, G and H. Abbreviations: st; stamen, le; lemma, fm; floral meristem, ec; ectopic carpel, pa; palea.

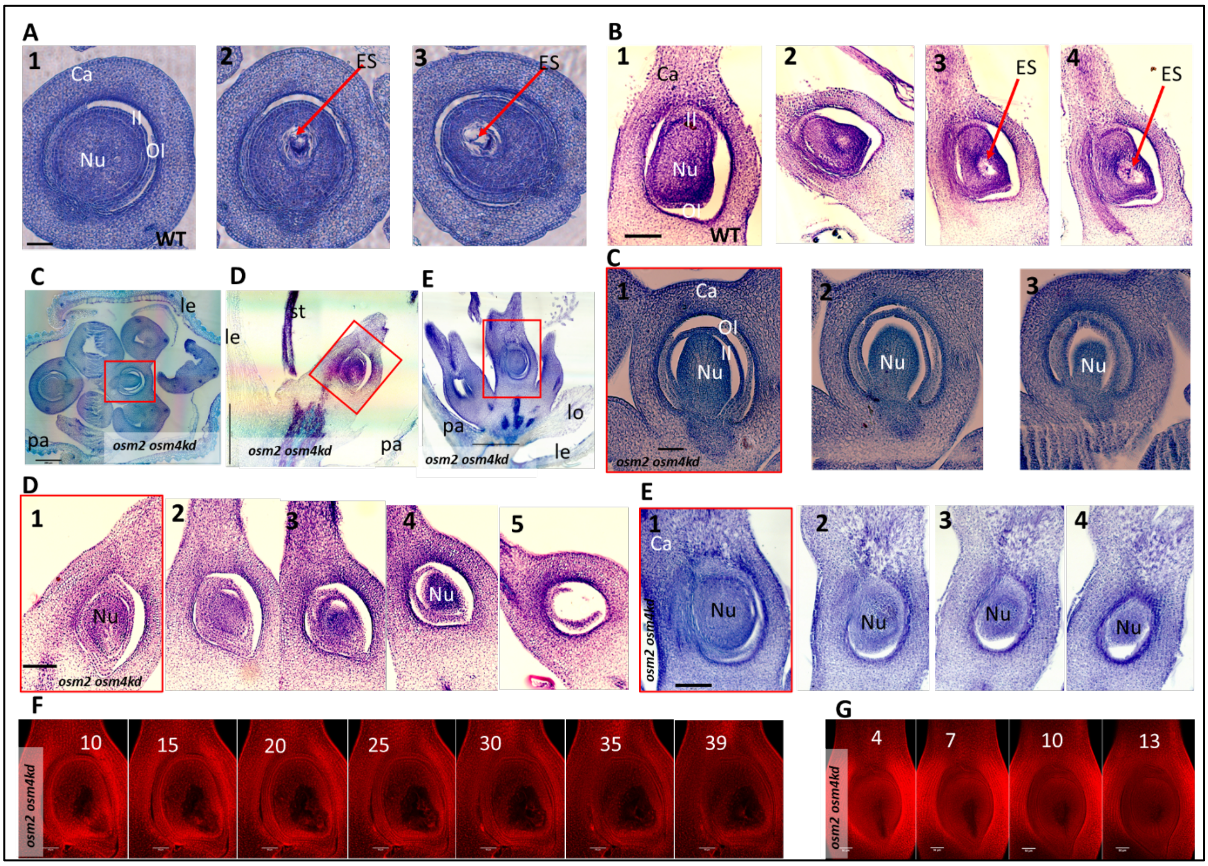


**Supplementary Figure S9. Supporting histological analyses of ovule differentiation in WT and *osmads2 osmads4kd* doubly perturbed florets. (Supports Figure 3).** A, Three transverse sections (TSs) across different basal to apical Z-axis planes in a WT ovary. B, Four longitudinal sections (LSs) across different Z-axis plane of a WT ovary. These serial sections show the embryo sac (ES) in differentiated WT ovule. C, TS in mature double mutant group I floret and three TSs across different basal to apical Z-axis planes of the central fourth whorl ovary. D, LS in mature double mutant group II floret and five LSs of the ovary and ovule. E, LS in mature double mutant group I floret and four LSs of the central fourth whorl ovary and ovule. Unlike WT, the double mutant ovary bears undifferentiated nucellar mass (Nu) that lacks the embryo sac. F, Optical sections (frames 10-39) in a double mutant ovary presented in Figure 2D. Sections/frames 20-30 pass through the center of the degenerated embryo sac. G, Optical sections (frames 4-13) in ovary presented in Figure 2H. Sections/frames 7 and 10 pass through the center of the degenerated embryo sac. Abbreviations: Inner integument (II), Outer Integument (OI), Lemma (le), Palea (pa), Stamen (st). Scale bars: 50 µm in A-G Z-axis plane sections in the ovaries while 200 µm in main panels of C-E displaying the full florets.

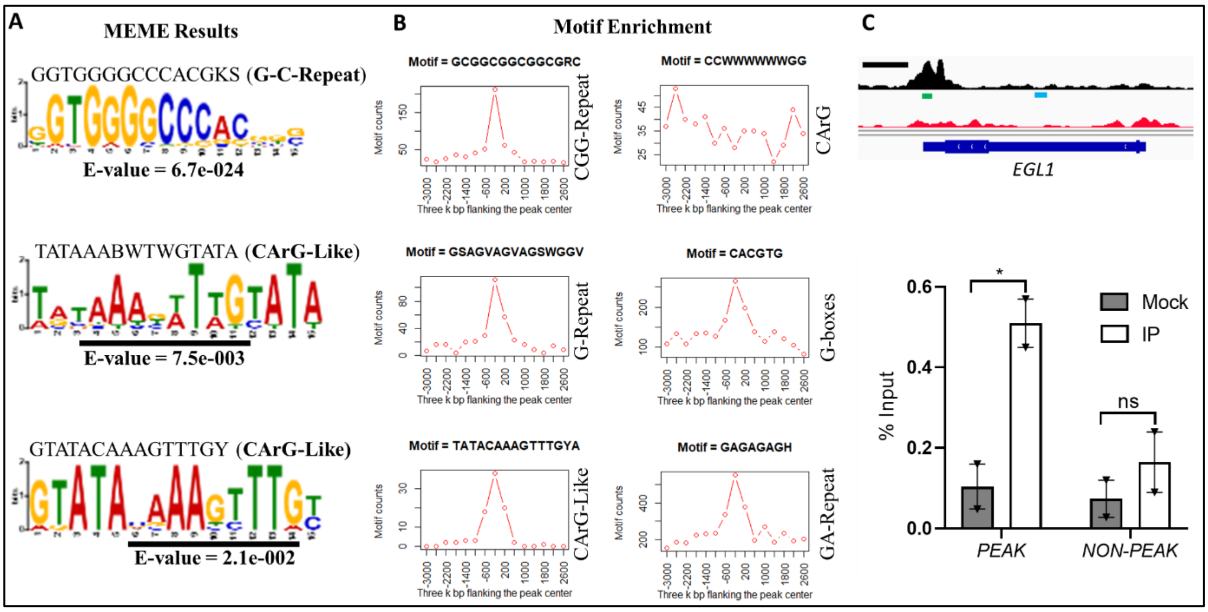


**Supplementary Figure S10. Supporting analyses for DNA *cis*-regulatory motifs in peak regions bound by OsMADS2 and ChIP-qPCR validation of binding at *EGL1*. (Supports Figure 5).** A, Representation of significantly enriched motifs in the peaks regions (MEME results). B, Relative distribution of the given motif around the OsMADS2 binding peak midpoint (considering 3,000 bp upstream and 3,000 bp downstream of the midpoint). Motifs retrieved by MEME analysis (CG-Repeat, G-Repeat, CArG-like), and motifs reported by Wuest et al. (2012, GA-Repeat and G-boxes) are enriched at/near the peak midpoint, while canonical CArG motifs are not enriched in the peak region. C, Validation of OsMADS2 binding to the *EGL1* locus in a WT panicle tissue pool of 1-2 cm inflorescences. A snapshot from IGV is shown where the amplified peak and nonpeak regions (Control) are marked by green and cyan lines, respectively. The black line above the IGV represents the scale for 1 kb genomic segment. In the bar graph y-axis represents the DNA pulled down by OsMADS2 association as a % of the Input value. Values are means ± SEM. Significance of enrichment of DNA segments in the OsMADS2 Chromatin IP over mock is tested by multiple unpaired Student's *t* test. P <0.05 denoted as *.

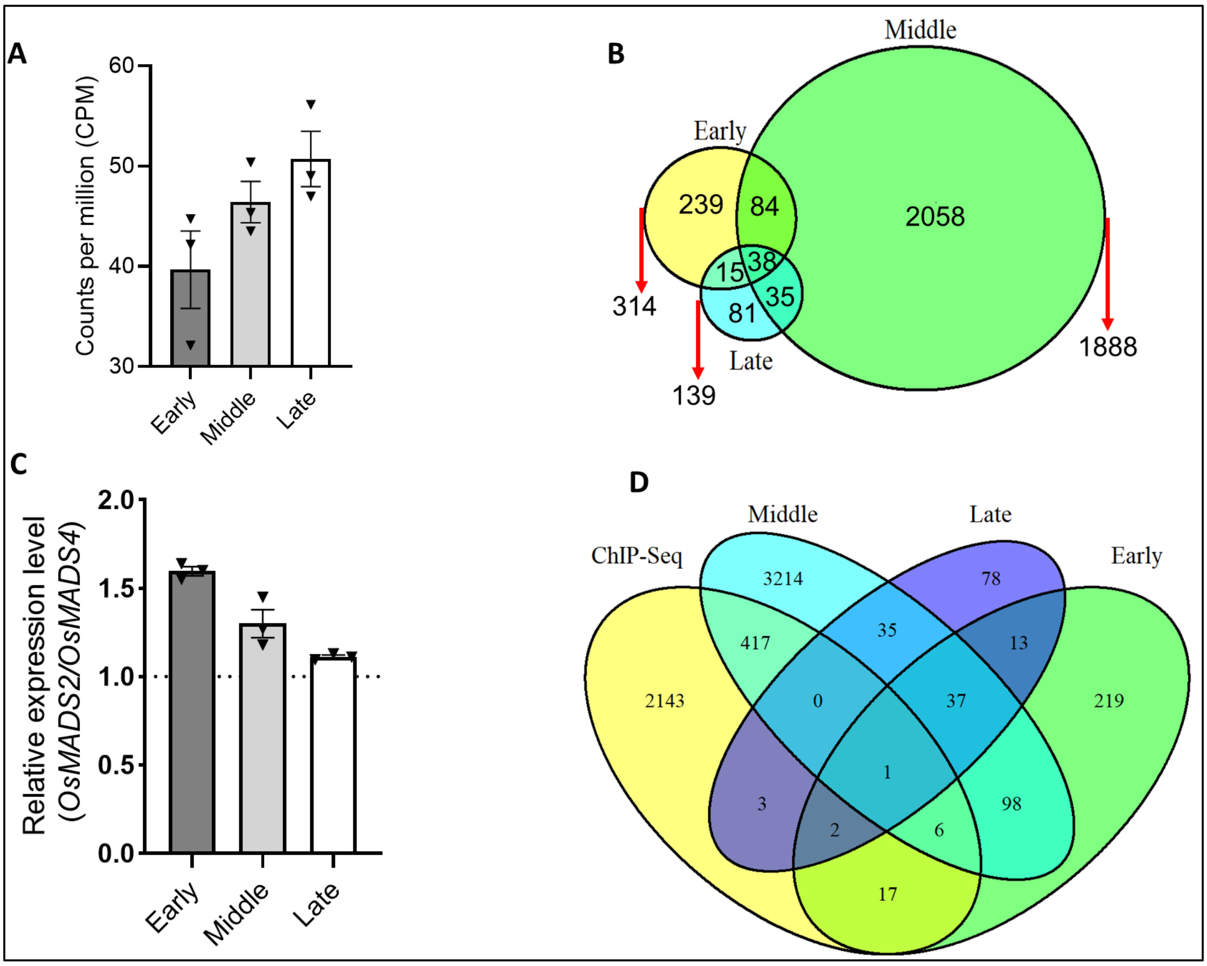


**Supplementary Figure 11**. **Supporting gene expression analyses. (Supports Figure 5 and 6).** A, Plot showing the normalized relative expression levels of *OsMADS2* with increasing expression levels in middle pool (comprising 0.5-1 cm inflorescences) as compared to the early pool (0.2-0.5 cm) inflorescences. B, Venn diagram representing the number of significant DEGs (absolute log2FC ≥1 and qValue ≤0.05) in three different developmental pools of inflorescences. C, Plot for the relative expression level of *OsMADS2* to *OsMADS4*. The trimmed mean of M (TMM) value for *OsMADS2* was divided by the TMM value for *OsMADS4* from the same RNA-seq library. In A and C, values are means ± SEM. D, Numerical representation of OsMADS2 directly and indirectly regulated genes. The Venn diagram shows overlap between the different RNA-Seq datasets (foldChange ≥1.5 and qValue ≤0.05) and the ChIP-Seq dataset.

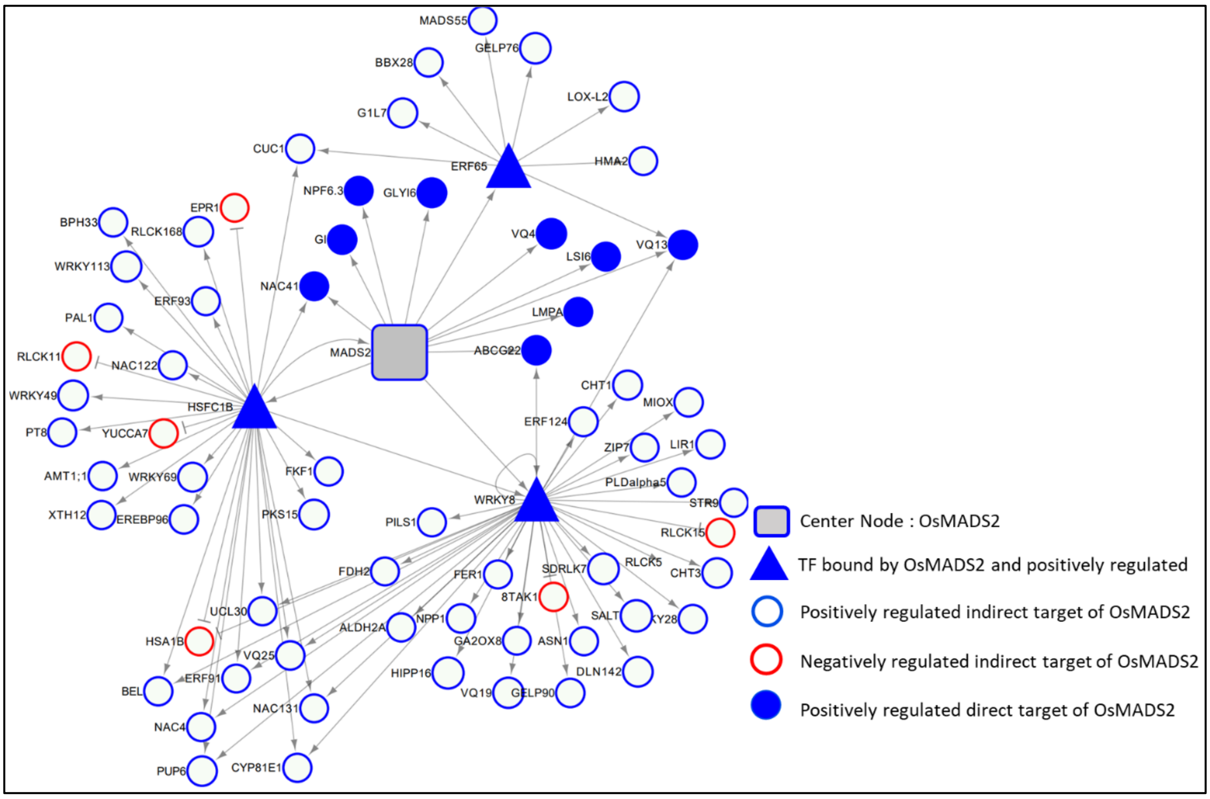


**Supplementary Figure S12. Gene Regulatory Network constructed based on ChIP-Seq and transcriptome data (RNA-Seq, tissue pool constituting 0.2-0.5 cm panicles). (Supports Figure 5 and 6).** The blue and red filled nodes represent positively- and negatively regulated direct downstream targets of OsMADS2, respectively. The unfilled circles of blue and red color boundaries represent positive and negative regulation by OsMADS2, respectively. The arrow shape and the T-shape indicate positive and negative regulation of the nodal gene, respectively. Tringle shape nodal genes represent transcription factors that were used to connect direct target transcription factors to downstream DEGs that form the second layer in this GRN.

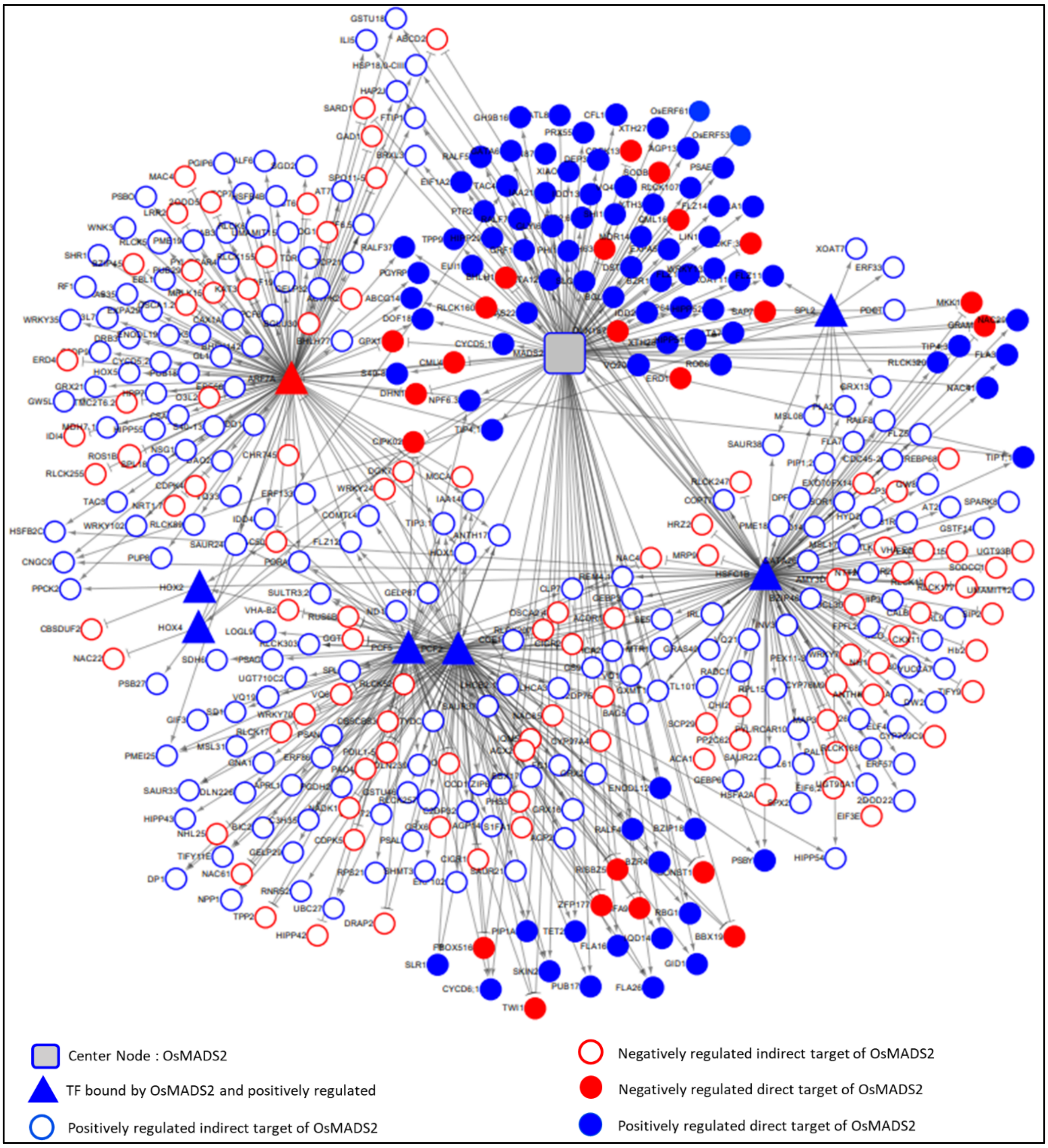


**Supplementary Figure S13. An OsMADS2 Gene Regulatory Network (GRN) constructed based on ChIP-Seq and transcriptome data (RNA-Seq, tissue pool constituting 0.5-1 cm panicles). (Supports Figure 5 and 6).** The filled blue and red nodes represent positively- and negatively regulated direct downstream targets of OsMADS2, respectively. The unfilled circles in blue and red color boundaries represent genes positively and negatively regulated by OsMADS2, respectively. Triangle shape denotes nodal OsMADS2 target genes coding for transcription factors (for e.g. PCF2 and PCF5) that were used to connect these direct target transcription factors to downstream DEGs that form the second layer in this GRN. The arrow shape and the T-shape indicate positive and negative regulation of each OsMADS2 gene target, respectively.

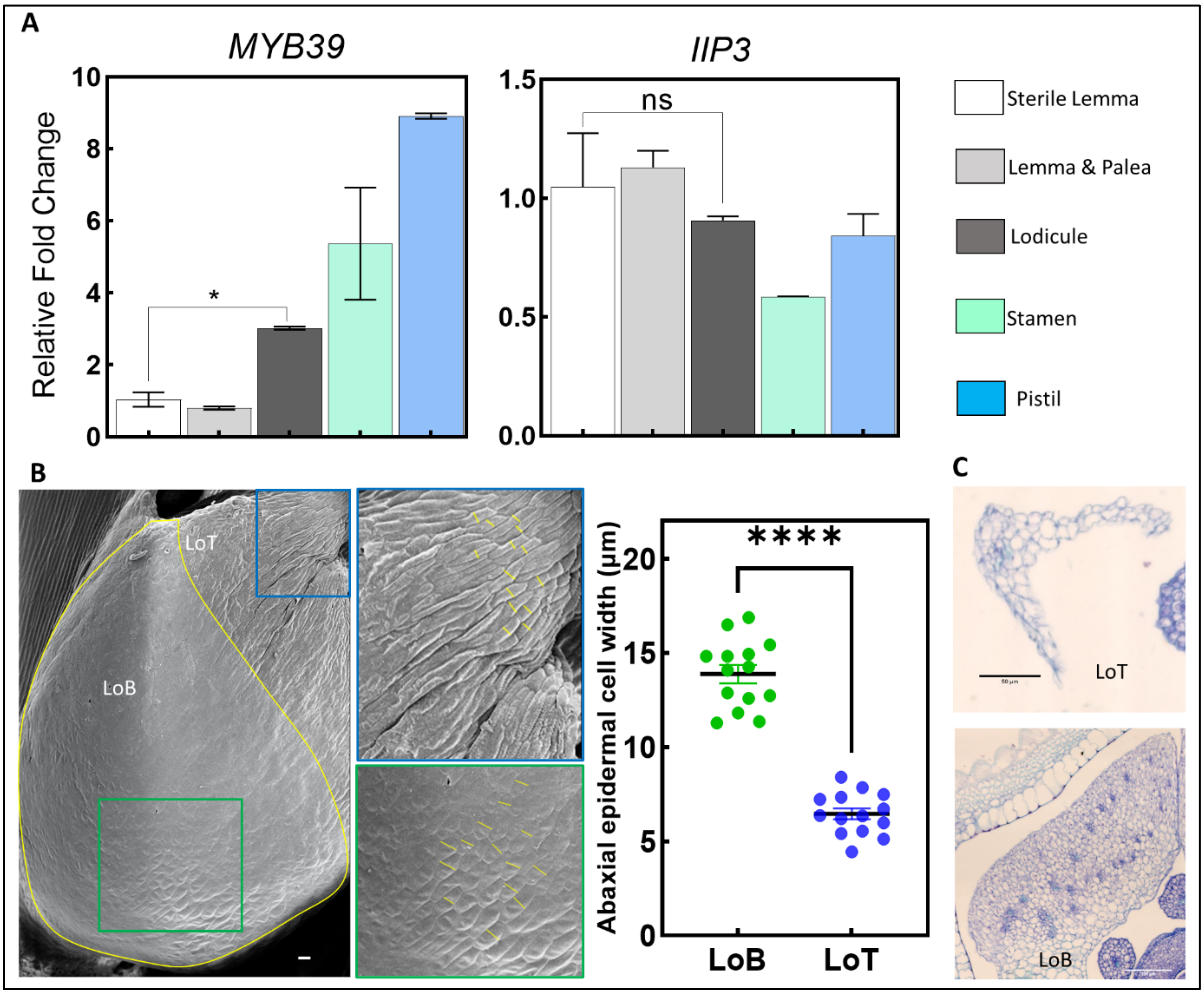


**Supplementary Figure S14. Supporting spatiotemporal gene expression analysis of OsMADS2 targets and characterization of the WT lodicule. (Supports Figure 7).** A, RT-qPCR analysis of two OsMADS2 gene targets, showing their relative expression levels in the mature floral organs from pre-anthesis WT florets. B, Scanning electron micrographs of the abaxial surface of the WT lodicule demonstrating difference in cell width between the body of lodicule (LoB) and the tip of the lodicule (LoT). In A and B, values are means ± SEM. $P < 0.05$ and $P < 0.0001$ denoted as * and ****, respectively; Student's *t* test. C, TS in the tip of lodicule (upper panel) and body of lodicule (lower panel).

**Supplementary Table S1: Analysis of segregation of different *osmads2* mutant alleles using the Chi-square test.**

| Parent and genotype | Segregated Progeny | # Expected progeny | #Observed progeny | % Expected progeny | % Observed progeny | P.value (two-tailed) | Signif-icance |
|---|---|---|---|---|---|---|---|
| 27A3a2B3 (*OsMADS2*$^{+/d8}$) | *osmads2*$^{d8/d8}$ | 11.75 | 9 | 25 | 19.15 | 0.35 | NS |
| | *OsMADS2*$^{+/d8}$; *OsMADS2*$^{+/+}$ | 35.25 | 83 | 75 | 80.85 | | |
| | Total | 47 | 47 | 100 | 100 | | |
| 27A3a2B12 (*osmads2*$^{i1/d2}$) | *osmads2*$^{i1/i1}$ | 2.25 | 1 | 25 | 11.11 | 0.24 | NS |
| | *osmads2*$^{i/1d2}$ | 4.5 | 7 | 50 | 77.78 | | |
| | *osmads2*$^{d2/d2}$ | 2.25 | 1 | 25 | 11.11 | | |

| | | | | | | | |
|---|---|---|---|---|---|---|---|
| | Total | 9 | 9 | 100 | 100 | | |
| 27A3a2B25 (*osmads2*$^{i1/d2}$) | *osmads2*$^{i1/i1}$ | 4.75 | 5 | 25 | 26.32 | 0.92 | NS |
| | *osmads2*$^{i/1d2}$ | 9.5 | 10 | 50 | 52.63 | | |
| | *osmads2*$^{d2/d2}$ | 4.75 | 4 | 25 | 21.05 | | |
| | Total | 19 | 19 | 100 | 100 | | |
| 27A3a2B31 (*osmads2*$^{i1/d2}$) | *osmads2*$^{i1/i1}$ | 6.5 | 10 | 25 | 38.46 | 0.023 | * |
| | *osmads2*$^{i/1d2}$ | 13 | 6 | 50 | 23.08 | | |
| | *osmads2*$^{d2/d2}$ | 6.5 | 10 | 25 | 38.46 | | |
| | Total | 26 | 26 | 100 | 100 | | |
| 27A3a2B12; 27A3a2B25; 27A3a2B12 | *osmads2*$^{i1/i1}$ | 13.50 | 16 | 25 | 29.63 | 0.54 | NS |
| | *osmads2*$^{i/1d2}$ | 27 | 23 | 50 | 42.59 | | |
| | *osmads2*$^{d2/d2}$ | 13.50 | 15 | 25 | 27.78 | | |
| | Total | 54 | 54 | 100 | 100 | | |

NS; not significant

### Supplementary Table S2: Summary of spikelet phenotyping data for WT and *osmads*$^{d8/d8}$

| **Phenotypic class** | **No. of WT florets (No. inner floral organs per floret)** | **No. of *osmads2d8/d8* florets (No. of inner floral organs per floret)** |
|---|---|---|
| Normal | 203 (9) | 0 |
| Only lodicules are abnormally elongated | 0 | 496 (9) |
| Abnormally elongated lodicules + medial lodicule | 0 | 2 (10) |
| Abnormally elongated lodicules with lodicule-stamen chimera and medial lodicule | 0 | 1 (10) |
| Abnormally elongated lodicules + medial lodicule +7 stamens + 2 carpels | 0 | 1 (12) |
| Abnormally elongated lodicules + medial lodicule +7 stamens | 0 | 12 (11) |
| Abnormally elongated lodicules + 7 stamens | 0 | 9 (10) |
| Abnormally elongated lodicule + 5 stamens | 0 | 2 (8) |
| Abnormally elongated lodicule + 2 carpels | 0 | 3 (10) |
| Abnormally elongated lodicule with lodicule-stamen chimera + 2 carpels | 0 | 1 (10) |
| Abnormally elongated lodicules with one of the six anthers underdeveloped and pollenless | 0 | 2 (9) |
| Cone-shaped organ instead of the lemma and palea and no inner floral organs | 0 | 1 (0) |

| | | |
|---|---|---|
| Abnormally elongated lodicules and enlarged sterile lemma | 1 | 3 (9) |

**Supplementary Table S3: Summary of genotyping and phenotyping data from the two phenotypic groups of *osmads2 osmads4kd* double mutant plants.**

| Plant Code | OsMADS2 locus | OsMADS4 dsRNAi transgene | Number of examine florets | Average stamen number* | Phenotype group |
|---|---|---|---|---|---|
| F2#26F3#5 | *osmads2*$^{d8/d8}$ | +/ | 5 | 5.8 | II |
| F2#26F3#14 | *osmads2*$^{d8/d8}$ | +/ | 3 | 5 | II |
| F2#26F3#30 | *osmads2*$^{d8/d8}$ | +/+ | 11 | 1.9 | I |
| F2#26F3#34 | *osmads2*$^{d8/d8}$ | +/ | 22 | 4.4 | II |
| F2#26F3#36 | *osmads2*$^{d8/d8}$ | +/+ | 19 | 2.1 | I |
| F2#26F3#57 | *osmads2*$^{d8/d8}$ | +/+ | 9 | 2.6 | I |
| F2#26F3#93 | *osmads2*$^{d8/d8}$ | +/ | 14 | 5.7 | II |
| C20 | *osmads2*$^{d8/d8}$ | +/+ | 41 | 2.4 | I |
| C21 | *osmads2*$^{d8/d8}$ | +/+ | 25 | 3.3 | I |
| C29 | *osmads2*$^{d8/d8}$ | +/ | 15 | 5.6 | II |

*Average stamen number is used as an indicative of the extent of *spw1*-like phenotype. The smaller number of stamens indicate a greater number of ectopic carpels

**Supplementary Table S4: Sequences of the different primers and oligos used in this study**

| Primer's Name | Sequences (Direction 5'→3') |
|---|---|
| **For Cloning** | |
| gRNA1(a) | GGCACAGGAGCGGGATCCTCAAGA |
| gRNA1(b) | AAACTCTTGAGGATCCCGCTCCTG |
| gRNA2(c) | GGCA CTCAATCATGATGAGCTCTT |
| gRNA2(d) | AAACAAGAGCTCATCATGATTGAG |
| OsMADS4 CDS FP | ATCGGATCCGAGGAGGAGTTGGATATG |
| OsMADS4 CDS RP | ATTGGTACCAAATTGGCAGTGCTCAG |
| OsMADS4 FP | GAGGTCGAGCTGAGCGGCGG |
| OsMADS4 RP | TGCTAATGCTGGGAGGAGCAC |
| **For Genotyping** | |
| OsMADS2 E3-I3 FP | CTCAGGTAAATTCTTCAGCC |
| OsMADS2 (Intron-V) RP | TGGCTAACAAAACACATGAG |
| p1300 5.0 | AGGGTTTCGCTCATGTGTTGAG |
| p1300 5.1 | TGTGTTGAGCATATAAGAAACCCTTAG |
| OsM4 ds IS FP | TCGTACGAAACACACTGTTTAGCTGTTTGA |
| OsM4 ds IS RP | CGATTCCTGGCGTCTGCCTAGT |
| **For RT-qPCR Analysis** | |
| OsMADS2 3' UTR FP | CACCACTGGATTGAATGTCCTAGTGC |
| OsMADS2 3' UTR RP | TGGTGCTATCTACAGCGAGGCAGAAT |
| OsMYB39 RT FP | CGCCGGCATCAAACTTGATTCGATCTC |
| OsMYB39 RT RP | AGCATTGTCAGGTTAACCCAAGAACGTG |
| Cyclin-P4-1-like qFP | GCTTCTTGTGTTTAGAGTTGTTTGCAGGT |
| Cyclin-P4-1-like qRP | GCTCTAGCCGATCCAACTCCAACT |
| GH9B16 qFP | AGCCGGCCACCTACATCAAC |
| GH9B16 qRP | ACTGCCACGTCACTCACACTCTCAC |

| | |
|---|---|
| CUC1 (2022) qRT FP | ATCGAAGGCTTCTTCTCGGGATGGG |
| CUC1 (2022) qRT RP | TCCTAGCTAATCCCTGGAGGTCTGGATG |
| TDR qRT FP | GGCCGGTGGAGAGCACTAC |
| TDR qRT RP | TCAAACGCGAGGTAATGCAGGTG |
| OsGI RT FP | TTGTGGATGCGCTTTGTGAC |
| OsGI RT RP | GCCTGCAGAACGATAGCAG |
| RCN2 RT FP | GTGAGTGAGCAATCGCAATC |
| RCN2 RT RP | CTAGGACTCTCCTGCCATGG |
| OsPTR2 qFP | GGCTGAACGAGCTGTGCTACAAG |
| OsPTR2 qRP | TCCACCGCCGTCACCATC |
| PCF2 qFP | TCAACAACCATCGTCCAGAACAGCAACTC |
| PCF2 Qrp | GTCGTCGTCGTCTTCGTCCTCCT |
| BGLU30 qRT FP | GGCCACCAGATTTGTGTTTCGTCAGTC |
| BGLU30 qRT RP | ATCACGCCATCATATCGGTCATCACCAC |
| MTR1 qRT FP | TGATCGCGTGAGTGCGTCTCTTTC |
| MTR1 qRT RP | AGAACCGTACAGGCAGTAGCACGATATG |
| HSFC1B qRT FP | CGCCTCGGTTTAGTTGCCTACTGTTAAG |
| HSFC1B qRT RP | TCCAACGATAGCTACTTAGCCAAACTCTCC |
| UBQ5 FP | ACCACTTCGACCGCCACTACT |
| UBQ5 RP | ACGCCTAAGCCTGCTGGTT |
| OsMADS55 FP | TCGCTTAATTCGTGCAAGTTATG |
| OsMADS55 FP | AAGTCTCAGCCGAGGTCACAA |
| OsMADS4 3'UTR FP | TCCTCCCAGCATTAGCACTCAGAA |
| OsMADS4 3'UTR RP | CCATCTGGAAACAAGAACAGTGGCT |
| OsSPL16 FP | GCCTTAGTTTCAGTAGAGTTGG |
| OsSPL16 RP | CTGCACAATACTGCAGTTCTCA |
| OsYUCCA3 qRT FP | GTGAGAACGGGCTCTACTCGGTCG |
| OsYUCCA3 qRT RP | GCTTATGCATGACCGATGAACACG |
| OsYUCCA7 FP | GGAGGTGCATCTCCGTCATCTTC |
| OsYUCCA7 RP | CACTGCTGTGTCCTACAATATCAC |
| IIP3 qRT FP | GTCTCAGGGCATGAACTTGTCAGG |
| IIP3 qRT RP | ACATTTCATTCACAAGAACCGCAGCT |
| GA20OX4 qRT FP | CTTCACCTGGGCCGACTTCA |
| GA20OX4 qRT RP | TAGCTAGCTGGACGCCCTAGAC |
| PME24 qRT FP | GATGTGCTAGCTCCTGTGTGTGTAGT |
| PME24 qRT RP | CCACATCGGAGCACATCGAGATCATAC |
| SPL qRT FP | GTGTTCTTGGGTAGGGAGAGGAAGA |
| SPL qRT RP | TACAGCCTCAGGGACAAGTCCAATG |
| OsTPKb qRT FP | GATCACTCCGGCACCCTCTC |
| OsTPKb qRT RP | TGGCGGTGAAATGAAATGTCTAACGC |
| OsDP1 FP | GCGCCGTACTGATCAACATC |
| OsDP1 RP | AGCTTGCAATTAAGCAGATTGCC |
| OsPIP1A qFP | TGCGCATCTGTCGATTCCCTCTATCT |
| OsPIP1A qRP | ACTGGATTACACGATTGAGTTGTTCAGGGT |
| JAZ6 qFP | ATCAATGCTCAATTCTGAGATGCCCCT |
| JAZ6 qRP | CGCCTTGTCATCGGATCATCTCCTAA |
| TIP4-1 qFP | AGAGGAGAGAACCCCACCTTGG |
| TIP4-1 qRP | ATGAAAAGGGCAGCAAAAAGTGCGA |
| JAZ7 qFP | GGTTGCATGGAAACTTCACCAAAGCT |
| JAZ7 qRP | GACAAGTGAACAACATAGAAGCACGAACT |
| OsMS2 qFP | TTCGACAATGGGAACACGGAGG |

| | |
|---|---|
| OsMS2 qRP | TCACGTCGAAGTGGAACCGC |
| OsHOX12 qFP | CGCCGACGGAGGATTAACTAGC |
| OsHOX12 qRP | CGTTAGGGCAACACGACTGATGA |
| DFOT1 qFP | GGGTGCTAAAATGGGTAGTATTGCTAGGC |
| DFOT1 qRP | CCACGTATCACCTCTCCTCTTCACAAC |
| CYCD2-2 qFP | GGACTCTGCTCCTGCTTCCAAG |
| CYCD2-2 qRP | GCTCGCCATGCACCACTACT |
| YUCCA1 qFP | GTAGTGGGAGCAGTGAAGGAGATGAC |
| YUCCA1 qRP | GCGAGGATTATTGTGTCGAATTGCTCC |
| OsTDD1 qFP | ATCACCCCAGAAGGCAAGAGGA |
| OsTDD1 qRP | TCACTTTCTGCCCTGAATCTACCCG |
| OsNIT1 qFP | GCAAGATAATCAGAGTCAATCGCAAAACTGA |
| OsNIT1 qRP | CTGGACTGCAATTTACAAGCAAGGGG |
| OG1 qFP | GCGGATGTAAATGTACTGCGGGATAAGTAA |
| OG1 qRP | AGCTAAATCACCTCACCGTTGCC |
| KRP3 qFP | GGGTAGAAGTGTCAACTCAACTCACACC |
| KRP3 qRP | GTCACTAACTCGATCCCCAGCCT |
| OsMADS50 qFP | GACCGTAACATCAACACCAC |
| OsMADS50 qRP | GAGATCCAGCTTATTCCTGG |
| Col15 qFP | AGGGTGAAAGGCCGGTTTGTGAAG |
| Col15 qRP | GCTAGCTATCAGGGATTGCAGGGATGAG |
| Ehd1 qFP | CGATTCCAACAACAAGCAAACACGGA |
| Ehd1 qRP | GTCCATGATTTCCTTGCATCCGTCT |
| GA20ox2 qFP | GCAAGAAGAAGCAAAACGTACGTGTG |
| GA20ox2 qRP | AGCATCCATTCATCCGTCGTTCCA |
| **For ChIP analysis** | |
| HSFC1B Peak OsM2 FP | CGAACCGGGAGGATACTTGCGAATTAC |
| HSFC1B Peak OsM2 RP | GGAGCTGGTGGACCTTGCACTC |
| HSFC1B Cont OsM2 FP | AGCGAGGAGGAAGAACGGAGAGA |
| HSFC1B Cont OsM2 RP | GCCCGAGGCCGGAGAGATTAC |
| Cyclin-P4-1 like Peak OsM2 FP | AGCTAGGCCCAAAGCTTCCGA |
| Cyclin-P4-1 like Peak OsM2 RP | AGGAGAGGCGTACGTATACGGGTATCT |
| Cyclin-P4-1 like Cont OsM2 FP | GGGTTGTCACACATCCAAGGAGCCTA |
| Cyclin-P4-1 like Cont OsM2 RP | GCAGTCATTGTCCCTGAGTCCCTCATC |
| GH9B16 Peak OsM2 FP | CGACGCAACAAACAGTATTTGCACGGA |
| GH9B16 Peak OsM2 RP | GCCGAGCCGAGTCAACAAGTACAC |
| GH9B16 Cont OsM2 FP | GGAACACAACAAATGCATCTGCCATCC |
| GH9B16 Cont OsM2 RP | AGTGGAATGAATGAACGCTCCACTCC |
| PTR2 Peak OsM2 FP | GGGAATGAATAGACGAGCAGAG |
| PTR2 Peak OsM2 RP | TTAGACTGATGGAACAGCATCTTG |
| PTR2 Cont OsM2 FP | GTTGATGTTGCTCTGGAGGTAG |
| PTR2 Cont OsM2 RP | TTCGCCTTCCTCGTCGTCGG |
| EGL1 Peak OsM2 FP | GTTAGCGAGGCACAGTTG |
| EGL1 Peak OsM2 RP | ACTATTACTATATAGAGAGGGTTGC |
| EGL1 Cont OsM2 FP | AGATGGAGATGTAATGGTTTGGT |
| EGL1 Cont OsM2 RP | CCACTGCACTCACTGACTATTAG |
| PCF2 Peak OsM2 FP | CCACCTACCTGATACGGAGATC |
| PCF2 Peak OsM2 RP | GAGCCTAAACATATTGTCAAGTAAAG |
| PCF2 Cont OsM2 FP | TAGATCGTGTCGAGGGCTAC |
| PCF2 Cont OsM2 RP | CCCGGTCTCTAAACTTGGATTTG |
| GL2.6 Peak OsM2 FP | AATCACTCCACCGTGATCCT |
| GL2.6 Peak OsM2 RP | GCCGCCATGCCATGTATAG |

| | |
|---|---|
| GL2.6 Cont OsM2 FP | AACACAGGAATAGTCGTTTGATTG |
| GL2.6 Cont OsM2 RP | TGAAATTCCTATGGATTGCCTAATG |
| DST Peak OsM2 FP | ATTGGGCCTGCGGGTCTTGT |
| DST M2 ChIP RP | TACACGCCCACATTGCTGCCATC |
| DST M2 Cont OsM2 FP | CCCAACAGAGTCGAGTAACAGCAGAGTT |
| DST M2 Cont OsM2 RP | CCCTTGCACTTAACATCACTTAGGAGCACA |
| ERF61 Peak OsM2 FP | TGCCTCCGACATCTCCGACAAC |
| ERF61 Peak OsM2 RP | GGACATGGGAAACATCTGTTCATGTCGAA |
| ERF61 Cont OsM2 FP | ATGAGATGCGGGCGTGCCAGTTAG |
| ERF61 Cont OsM2 RP | GCTGACATTGCTCCATCGGCTGTC |
| GATA12 Peak OsM2 FP | TTTTCGCAAAGCAGCAAACAGCGG |
| GATA12 Peak OsM2 RP | CAGCAGCAAAGCCTCCATCCG |
| GATA12 Cont OsM2 FP | CAGGTGTTGTGAGCCCAAATAGTCAGGT |
| GATA12 Cont OsM2 FP | ACGCCTGATGGAGAAACACAGGAACA |
| **For *in situ* hybridization** | |
| OsPTR2 qFP | GGCTGAACGAGCTGTGCTACAAG |
| OsPTR2 qRP | TCCACCGCCGTCACCATC |
| OsPIP1-1 ISH FP | TCAGGGCGATCCCATTCAAGAGCAG |
| OsPIP1-1 qRP | ACTGGATTACACGATTGAGTTGTTCAGGGT |
| Cyclin-P4-1-like qFP | GCTTCTTGTGTTTAGAGTTGTTTGCAGGT |
| Cyclin-P4-1-Like ISH RP | ACTCTGACAAAGCTAACAGTTGCTCAATATAT |
