## Supplementary material for "Functional genetics of rice *PISTILLATA* genes unravels new roles and targets in flowering time, female fertility and parthenocarpy": Main MS and figures as a word file

**Running Title:** Roles of rice *PI* paralogs in floral development

**Authors:** Mohamed Zamzam^1,4^, Ritabrata Basak^1#^, Sharad Singh ^1#^, Sandhan Prakash ^1#^, Raghavaram Peesapati^1^, Imtiyaz Khanday^1,3^, Sara Simonini^2^, Ueli Grossniklaus^2^ and Usha Vijayraghavan*^1^

^1^ Department of Microbiology and Cell Biology, Indian Institute of Science, Bangalore 560012, India

^2^ Institute of Plant and Microbial Biology, University of Zurich, Zollikerstrasse 107, CH8008, Zurich, Switzerland

^3^ Present address: Department of Plant Sciences, University of California, Davis, 95616, USA

^4^Present address: Branch of Genetics, Department of Agricultural Botany, Faculty of Agriculture at Cairo, Al-Azhar University, Cairo, 11651, Egypt, Visiting Faculty, Indian Institute of Science, Bangalore 560012, India

\# Shared second authors

*****Corresponding author: Usha Vijayraghavan

Department of Microbiology and Cell Biology,

Indian Institute of Science, Bangalore 560012, India

**Funding**

This study was funded by the Department of Biotechnology, Ministry of Science and Technology, Government of India, grant No. BT/PR38988/AGIII/103/1232/2020 to U.V. This work was also supported by the University of Zurich, a grant of the Swiss National Science Foundation (31003A_179553 to U.G.), and an Infosys Visiting Chair Professorship (to U.G.). Research fellowships to M.Z., S.P. and R.P. were received from the Indian Institute of Science. RB is supported by the Prime Minister's Research Fellowship, Ministry of Education, Government of India. M.Z. is currently funded by the International Visiting Faculty Program of the Indian Institute of Science. U.V. holds the J N Tata Chair at the Indian Institute of Science.

**Author contributions**

UV, UG and MZ designed the research. MZ, SS^1^, SP, RB, IK, and SS^2^ performed the experiments. RP and MZ performed the bioinformatic analyses. Manuscript writing was done by MZ, UG, and UV with contributions from IK and SS. All authors read and approved the final manuscript.

**References**

**Abu-Zaitoon YM.** Phylogenetic analysis of putative genes involved in the tryptophan-dependent pathway of auxin biosynthesis in rice. Appl Biochem Biotechnol. 2014:**172**(5):2480–2495. https://doi.org/10.1007/S12010-013-0710-4/FIGURES/7

**Ahmad I, Devonshire J, Mohamed R, Schultze M, and Maathuis FJM**. Overexpression of the potassium channel TPKb in small vacuoles confers osmotic and drought tolerance to rice. New Phytol. 2016:**209**(3):1040–1048. https://doi.org/10.1111/NPH.13708

**Akiyama T, Jin S, Yoshida M, Hoshino T, Opassiri R, and Ketudat Cairns JR**. Expression of an endo-(1,3;1,4)-beta-glucanase in response to wounding, methyl jasmonate, abscisic acid and ethephon in rice seedlings. J Plant Physiol. 2009:**166**(16):1814–1825. https://doi.org/10.1016/J.JPLPH.2009.06.002

**Ambrose BA, Lerner DR, Ciceri P, Padilla CM, Yanofsky MF, and Schmidt RJ**. Molecular and Genetic Analyses of the *Silky1* Gene Reveal Conservation in Floral Organ Specification between Eudicots and Monocots. Mol Cell. 2000:**5**(3):569–579. https://doi.org/10.1016/S1097-2765(00)80450-5

**Balázsi G, Van Oudenaarden A, and Collins JJ**. Cellular decision making and biological noise: from microbes to mammals. Cell. 2011:**144**(6):910–925. https://doi.org/10.1016/J.CELL.2011.01.030

**Bartlett ME, Williams SK, Taylor Z, Deblasio S, Goldshmidt A, Hall DH, Schmidt RJ, Jackson DP, and Whipplea CJ**. The Maize *PI/GLO* Ortholog *Zmm16/sterile tassel silky ear1* Interacts with the Zygomorphy and Sex Determination Pathways in Flower Development. Plant Cell. 2015:**27**(11):3081–3098. https://doi.org/10.1105/TPC.15.00679

**Bowman JL, Smyth DR, and Meyerowitz EM**. Genes directing flower development in *Arabidopsis*. Plant Cell. 1989:**1**(1):37–52. https://doi.org/10.1105/TPC.1.1.37

**Busk PK and Pagès M**. Microextraction of Nuclear Proteins from Single Maize Embryos. Plant Mol Biol Report. 1997:**15**(4):371–376. https://doi.org/10.1023/A:1007428802474/METRICS

**Cao L, Tian J, Liu Y, Chen X, Li S, Persson S, Lu D, Chen M, Luo Z, Zhang D, et al.** Ectopic expression of OsJAZ6, which interacts with OsJAZ1, alters JA signaling and spikelet development in rice. Plant J. 2021:**108**(4):1083–1096. https://doi.org/10.1111/TPJ.15496

**Chen S, Zhou Y, Chen Y, and Gu J**. fastp: an ultra-fast all-in-one FASTQ preprocessor. Bioinformatics. 2018:**34**(17):i884–i890. https://doi.org/10.1093/BIOINFORMATICS/BTY560

**Choi MG, Kim EJ, Song JY, Choi SB, Cho SW, Park CS, Kang CS, and Park Y Il**. Peptide transporter2 (PTR2) enhances water uptake during early seed germination in *Arabidopsis thaliana*. Plant Mol Biol. 2020:**102**(6):615–624. https://doi.org/10.1007/s11103-020-00967-3 **Chow CN, Lee TY, Hung YC, Li GZ, Tseng KC, Liu YH, Kuo PL, Zheng HQ, and Chang WC**. PlantPAN3.0: a new and updated resource for reconstructing transcriptional regulatory networks from ChIP-seq experiments in plants. Nucleic Acids Res. 2019:**47**(D1):D1155–D1163. https://doi.org/10.1093/NAR/GKY1081

**Chung YY, Kim SR, Kang HG, Noh YS, Park MC, Finkel D, and An G**. Characterization of two rice MADS box genes homologous to *GLOBOSA*. Plant Sci. 1995:**109**(1):45–56. https://doi.org/10.1016/0168-9452(95)04153-L

**Coen ES and Meyerowitz EM**. The war of the whorls: genetic interactions controlling flower development. Nat 1991 3536339. 1991:**353**(6339):31–37. https://doi.org/10.1038/353031a0

**Cosgrove DJ**. Enzymes and other agents that enhance cell wall extensibility. Annu Rev Plant Physiol Plant Mol Biol. 1999:**50**:391–417. https://doi.org/10.1146/ANNUREV.ARPLANT.50.1.391

**Dobin A, Davis CA, Schlesinger F, Drenkow J, Zaleski C, Jha S, Batut P, Chaisson M, and Gingeras TR**. STAR: ultrafast universal RNA-seq aligner. Bioinformatics. 2013:**29**(1):15–21. https://doi.org/10.1093/BIOINFORMATICS/BTS635

**Doi K, Izawa T, Fuse T, Yamanouchi U, Kubo T, Shimatani Z, Yano M, and Yoshimura A**. Ehd1, a B-type response regulator in rice, confers short-day promotion of flowering and controls *FT*-like gene expression independently of Hd1. Genes Dev. 2004:**18**(8):926. https://doi.org/10.1101/GAD.1189604

**Dreni L, Jacchia S, Fornara F, Fornari M, Ouwerkerk PBF, An G, Colombo L, and Kater MM**. The D-lineage MADS-box gene *OsMADS13* controls ovule identity in rice. Plant J. 2007:**52**(4):690–699. https://doi.org/10.1111/J.1365-313X.2007.03272.X

**Elowitz MB, Levine AJ, Siggia ED, and Swain PS**. Stochastic gene expression in a single cell. Science. 2002:**297**(5584):1183–1186. https://doi.org/10.1126/SCIENCE.1070919

**Ewels P, Magnusson M, Lundin S, and Käller M**. MultiQC: summarize analysis results for multiple tools and samples in a single report. Bioinformatics. 2016:**32**(19):3047–3048. https://doi.org/10.1093/BIOINFORMATICS/BTW354

**Jack T, Fox GL, and Meyerowitz EM**. *Arabidopsis* homeotic gene *APETALA3* ectopic expression: transcriptional and posttranscriptional regulation determine floral organ identity. Cell. 1994:**76**(4):703–716. https://doi.org/10.1016/0092-8674(94)90509-6

**Ji J and Braam J**. Restriction Site Extension PCR: A novel method for high-throughput characterization of tagged DNA fragments and genome walking. PLoS One. 2010:**5**(5):e10577. https://doi.org/10.1371/JOURNAL.PONE.0010577

**Joldersma D and Liu Z**. The making of virgin fruit: the molecular and genetic basis of parthenocarpy. J Exp Bot. 2018:**69**(5):955–962. https://doi.org/10.1093/JXB/ERX446

**Kang HG, Jeon JS, Lee S, and An G**. Identification of class B and class C floral organ identity genes from rice plants. Plant Mol Biol. 1998:**38**(6):1021–1029. https://doi.org/10.1023/A:1006051911291/METRICS

**Kaufmann K, Muiño JM, Jauregui R, Airoldi CA, Smaczniak C, Krajewski P, and Angenent GC**. Target genes of the MADS transcription factor SEPALLATA3: integration of developmental and hormonal pathways in the *Arabidopsis* flower. PLoS Biol. 2009:**7**(4):0854–0875. https://doi.org/10.1371/JOURNAL.PBIO.1000090

**Klepikova A V., Logacheva MD, Dmitriev SE, and Penin AA**. RNA-seq analysis of an apical meristem time series reveals a critical point in *Arabidopsis thaliana* flower initiation. BMC Genomics. 2015:**16**(1):1–16. <https://doi:10.1186/s12864-015-1688-9>

**Krizek BA and Fletcher JC**. Molecular mechanisms of flower development: an armchair guide. Nat Rev Genet 2005 69. 2005:**6**(9):688–698. https://doi.org/10.1038/nrg1675

**Krizek BA and Meyerowitz EM**. The *Arabidopsis* homeotic genes *APETALA3* and *PISTILLATA* are sufficient to provide the B class organ identity function. Development. 1996:**122**(1):11–22. <https://doi.org/10.1242/DEV.122.1.11>

**Krizek BA**. Auxin regulation of Arabidopsis flower development involves members of the AINTEGUMENTA-LIKE/PLETHORA (AIL/PLT) family. J Exp Bot. 2011:**62**(10):3311–3319. https://doi.org/10.1093/JXB/ERR127

**Lee JH, Park SH, and Ahn JH**. Functional conservation and diversification between rice OsMADS22/OsMADS55 and *Arabidopsis* SVP proteins. Plant Sci. 2012:**185**–**186**:97–104. https://doi.org/10.1016/J.PLANTSCI.2011.09.003

**Lee S, Jeon JS, An K, Moon YH, Lee S, Chung YY, and An G**. Alteration of floral organ identity in rice through ectopic expression of *OsMADS16*. Planta. 2003:**217**(6):904–911. https://doi.org/10.1007/S00425-003-1066-8

**Li N, Zhang DS, Liu HS, Yin CS, Li XX, Liang WQ, Yuan Z, Xu B, Chu HW, Wang J, et al.** The rice *tapetum degeneration retardation* gene is required for tapetum degradation and anther development. Plant Cell. 2006:**18**(11):2999–3014. https://doi.org/10.1105/TPC.106.044107

**Li Q, Tong T, Jiang W, Cheng J, Deng F, Wu X, Chen ZH, Ouyang Y, and Zeng F**. Highly Conserved Evolution of Aquaporin PIPs and TIPs Confers Their Crucial Contribution to Flowering Process in Plants. Front Plant Sci. 2022:**12**:2945. https://doi.org/10.3389/FPLS.2021.761713/BIBTEX

**Li S, Wei K, Zhang L, Ning Y, Lu F, Wang X, Guo Y, Liu L, Li X, Zhu C, et al.** Fine mapping and candidate gene validation of tomato gene *Carpelloid Stamen and Parthenocarpy* (*CSP*). Horticulturae. 2024:**10**(4):403. https://doi.org/10.3390/HORTICULTURAE10040403/S1

**Li Y, Ouyang J, Wang YY, Hu R, Xia K, Duan J, Wang Y, Tsay YF, and Zhang M**. Disruption of the rice nitrate transporter OsNPF2.2 hinders root-to-shoot nitrate transport and vascular development. Sci Reports 2015 51. 2015:**5**(1):1–10. https://doi.org/10.1038/srep09635

**Liang M, Deng L, Liu J, He A, and Chen L.** Interaction between the *eui* gene and thermo-sensitive genic male sterility in rice. Euphytica. 2008:**164**(3):637–643. https://doi.org/10.1007/S10681-008-9657-X/TABLES/2

**Liao Y, Smyth GK, and Shi W**. The R package Rsubread is easier, faster, cheaper and better for alignment and quantification of RNA sequencing reads. Nucleic Acids Res. 2019:**47**(8):e47–e47. https://doi.org/10.1093/NAR/GKZ114

**Lohmann JU and Weigel D**. Building beauty: the genetic control of floral patterning. Dev Cell. 2002:**2**(2):135–142. https://doi.org/10.1016/S1534-5807(02)00122-3

**Luan X, Liu S, Ke S, Dai H, Xie XM, Hsieh TF, and Zhang XQ**. Epigenetic modification of *ESP*, encoding a putative long noncoding RNA, affects panicle architecture in rice. Rice. 2019:**12**(1). https://doi.org/10.1186/S12284-019-0282-1

**Machanick P and Bailey TL**. MEME-ChIP: motif analysis of large DNA datasets. Bioinformatics. 2011:**27**(12):1696–1697. https://doi.org/10.1093/BIOINFORMATICS/BTR189

**Mizutani M, Naganuma T, Tsutsumi KI, and Saitoh Y**. The syncytium-specific expression of the Orysa;KRP3 CDK inhibitor: implication of its involvement in the cell cycle control in the rice (*Oryza sativa* L.) syncytial endosperm. J Exp Bot. 2010:**61**(3):791–798. https://doi.org/10.1093/JXB/ERP343

**Nagasawa N, Miyoshi M, Sano Y, Satoh H, Hirano H, Sakai H, and Nagato Y**. *SUPERWOMAN1* and *DROOPING LEAF* genes control floral organ identity in rice. Development. 2003:**130**(4):705–718. https://doi.org/10.1242/DEV.00294

**Nandi Ak, Kushalappa K, Prasad K, and Vijayraghavan U**. A conserved function for *Arabidopsis* *SUPERMAN* in regulating floral-whorl cell proliferation in rice, a monocotyledonous plant. Curr Biol. 2000:**10**(4):215–218. https://doi.org/10.1016/S0960-9822(00)00341-9

**Ning J, Zhang B, Wang N, Zhou Y, and Xiong L**. Increased Leaf Angle1, a Raf-Like MAPKKK that interacts with a nuclear protein family, regulates mechanical tissue formation in the lamina joint of rice. Plant Cell. 2011:**23**(12):4334–4347. https://doi.org/10.1105/TPC.111.093419

**O’Toole JC and Namuco OS**. Role of panicle exsertion in water stress induced sterility. Crop Sci. 1983:**23**(6):1093–1097. https://doi.org/10.2135/CROPSCI1983.0011183X002300060017X

**Pagès H, Aboyoun P, Gentleman R, and DebRoy S.** *Biostrings: Efficient manipulation of biological strings*. R package version 2.64.0, (2022). https://bioconductor.org/packages/Biostrings

**Pinyopich A, Ditta GS, Savidge B, Liljegren SJ, Baumann E, Wisman E, and Yanofsky MF**. Assessing the redundancy of MADS-box genes during carpel and ovule development. Nat 2003 4246944. 2003:**424**(6944):85–88. https://doi.org/10.1038/nature01741

**Prakash S, Rai R, Zamzam M, Ahmad O, Peesapati R, and Vijayraghavan U**. OsbZIP47 is an integrator for meristem regulators during rice plant growth and development. Front Plant Sci. 2022:**13**:883. https://doi.org/10.3389/FPLS.2022.865928/BIBTEX

**Prasad K and Vijayraghavan U**. Double-stranded RNA interference of a rice *PI/GLO* paralog, *OsMADS2*, uncovers its second-whorl-specific function in floral organ patterning. Genetics. 2003:**165**(4):2301–2305. https://doi.org/10.1093/GENETICS/165.4.2301

**Ren L, Tang D, Zhao T, Zhang F, Liu C, Xue Z, Shi W, Du G, Shen Y, Li Y, et al.** OsSPL regulates meiotic fate acquisition in rice. New Phytol. 2018:**218**(2):789–803. https://doi.org/10.1111/NPH.15017

**Robinson MD, McCarthy DJ, and Smyth GK**. edgeR: a Bioconductor package for differential expression analysis of digital gene expression data. Bioinformatics. 2010:**26**(1):139–140. https://doi.org/10.1093/BIOINFORMATICS/BTP616

**Tan H, Liang W, Hu J, and Zhang D**. *MTR1* encodes a secretory fasciclin glycoprotein required for male reproductive development in rice. Dev Cell. 2012:**22**(6):1127–1137. https://doi.org/10.1016/J.DEVCEL.2012.04.011

**Yang Y, Xiang H, and Jack T**. *pistillata-5*, an *Arabidopsis* B class mutant with strong defects in petal but not in stamen development. Plant J. 2003:**33**(1):177–188. https://doi.org/10.1046/J.1365-313X.2003.01603.X

**Yao JL, Dong YH, and Morris BAM.** Parthenocarpic apple fruit production conferred by transposon insertion mutations in a MADS-box transcription factor. Proc Natl Acad Sci U S A. 2001:**98**(3):1306–1311. https://doi.org/10.1073/PNAS.98.3.1306

**Yao SG, Ohmori S, Kimizu M, and Yoshida H**. Unequal genetic redundancy of rice PISTILLATA orthologs, OsMADS2 and OsMADS4, in lodicule and stamen development. Plant Cell Physiol. 2008:**49**(5):853–857. https://doi.org/10.1093/PCP/PCN050

https://doi.org/10.1073/PNAS.0907896106/SUPPL_FILE/0907896106SI.PDF

**Yoshida H, Itoh JI, Ohmori S, Miyoshi K, Horigome A, Uchida E, Kimizu M, Matsumura Y, Kusaba M, Satoh H, et al.** *superwoman1*-cleistogamy, a hopeful allele for gene containment in GM rice. Plant Biotechnol J. 2007:**5**(6):835–846. https://doi.org/10.1111/J.1467-7652.2007.00291.X

**Yu G, Wang LG, and He QY**. ChIPseeker: an R/Bioconductor package for ChIP peak annotation, comparison and visualization. Bioinformatics. 2015:**31**(14):2382–2383. https://doi.org/10.1093/BIOINFORMATICS/BTV145

**Yun D, Liang W, Dreni L, Yin C, Zhou Z, Kater MM, and Zhang D**. *OsMADS16* genetically interacts with *OsMADS3* and *OsMADS58* in specifying. Mol Plant. 2013:**6**(3):743–756. https://doi.org/10.1093/mp/sst003

**Zeng YX, Hu CY, Lu YG, Li JQ, and Liu XD**. Diversity of abnormal embryo sacs in indica/japonica hybrids in rice demonstrated by confocal microscopy of ovaries. Plant Breed. 2007:**126**(6):574–580. https://doi.org/10.1111/J.1439-0523.2007.01380.X

**Zhang Y, Liu T, Meyer CA, Eeckhoute J, Johnson DS, Bernstein BE, Nussbaum C, Myers RM, Brown M, Li W, et al.** Model-based analysis of ChIP-Seq (MACS). Genome Biol. 2008:**9**(9):1–9. https://doi.org/10.1186/gb-2008-9-9-r137

**Zhao ZX, Yin XX, Li S, Peng YT, Yan XL, Chen C, Hassan B, Zhou SX, Pu M, Zhao JH, et al.** miR167d-ARFs module regulates flower opening and stigma size in rice. Rice. 2022:**15**(1):1–15. https://d oi: 10.1186/s12284-022-00587-z

**Zhu LJ, Gazin C, Lawson ND, Pagès H, Lin SM, Lapointe DS, and Green MR**. ChIPpeakAnno: A Bioconductor package to annotate ChIP-seq and ChIP-chip data. BMC Bioinformatics. 2010:**11**(1):1–10. https://doi.org/10.1186/1471-2105-11-237/TABLES/2

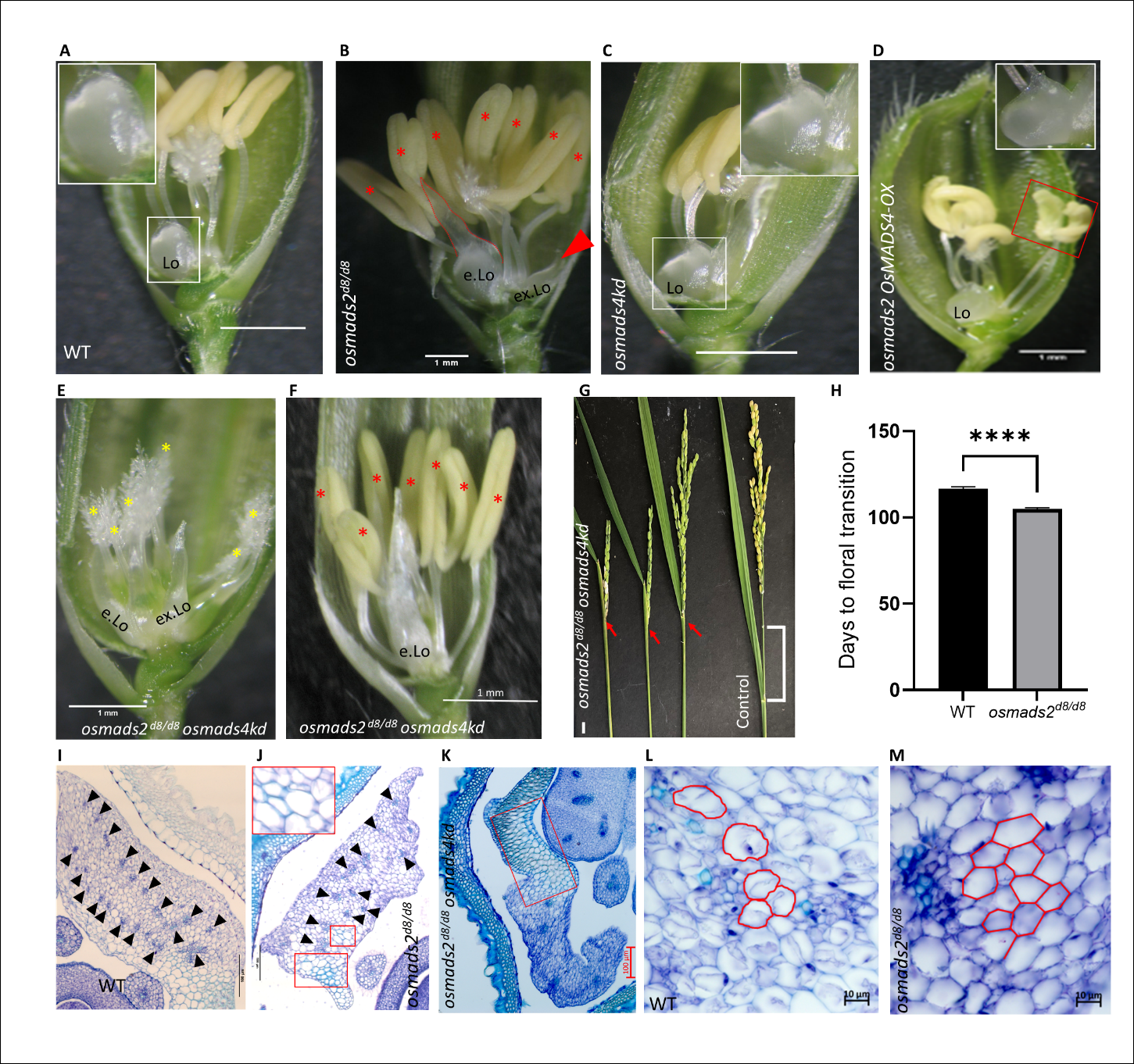
**Figure 1. Phenotypic and histological analyses of the different genotypes studied.** A, WT mature pre-anthesis spikelet dissected to show the lodicule. The inset shows a magnification of the normal lodicule. B, *osmads2* mutant spikelet dissected to show the abnormal organs. The red asterisks mark the increased stamen number and the red arrowhead marks the extra lodicule (ex.Lo). C, Spikelet dissected from *osmads4kd#1* showing normal floral organs. D, Spikelet from *osmads2 OsMADS4-OX* line #3 showing normal lodicule (complemented) but with abnormal crescent-like anthers (red rectangle). The number of floral organs is unchanged as compared to WT (n=300). E and F, Spikelets from *osmads2^d8/d8^ osmads4kd* of representative group I and group II phenotypes, respectively. The yellow asterisks represent the main and ectopic carpels in group I phenotype and the red asterisks mark the increased stamen number in group II phenotype. The elongated lodicule (e.Lo) is flattened throughout the proximal distal axis (compare lodicule in E and F *vs.* in A and B). G, Panicle partial enclosure phenotype in *osmads2^d8/d8^ osmads4kd* plants. H, Graph plot representing the number of days taken to floral transition in WT plants (n = 79) *vs.* *osmads2^d8/d8^* plants (n=125). Values are means ± the standard error of the mean (SEM). P < 0.0001 denoted as ****; Student’s *t* test. I-K, TS sections of lodicules in WT and *osmads2^d8/d8^* and *smads2^d8/d8^ osmads4kd*, respectively*.* The black arrowheads mark the vascular bundles which are reduced in number and of abnormal distribution across the lodicule of *osmads2^d8/d8^*. The red rectangles (in J and K) mark regions with green stained cell outlines, indicative of cell wall lignification in tissue sections stained with Toluidine blue. L and M, TS of lodicules in WT and *osmads2^d8/d8^* at high magnification showing the cell shape in the ground tissues. Compare the brick-like parenchyma in L to the polygonal like cells in M. Cell outlines are marked in red. Scale bars: 1 mm in A-F, 1 cm in G, 100 µm in I-K, 10 µm in L and M.

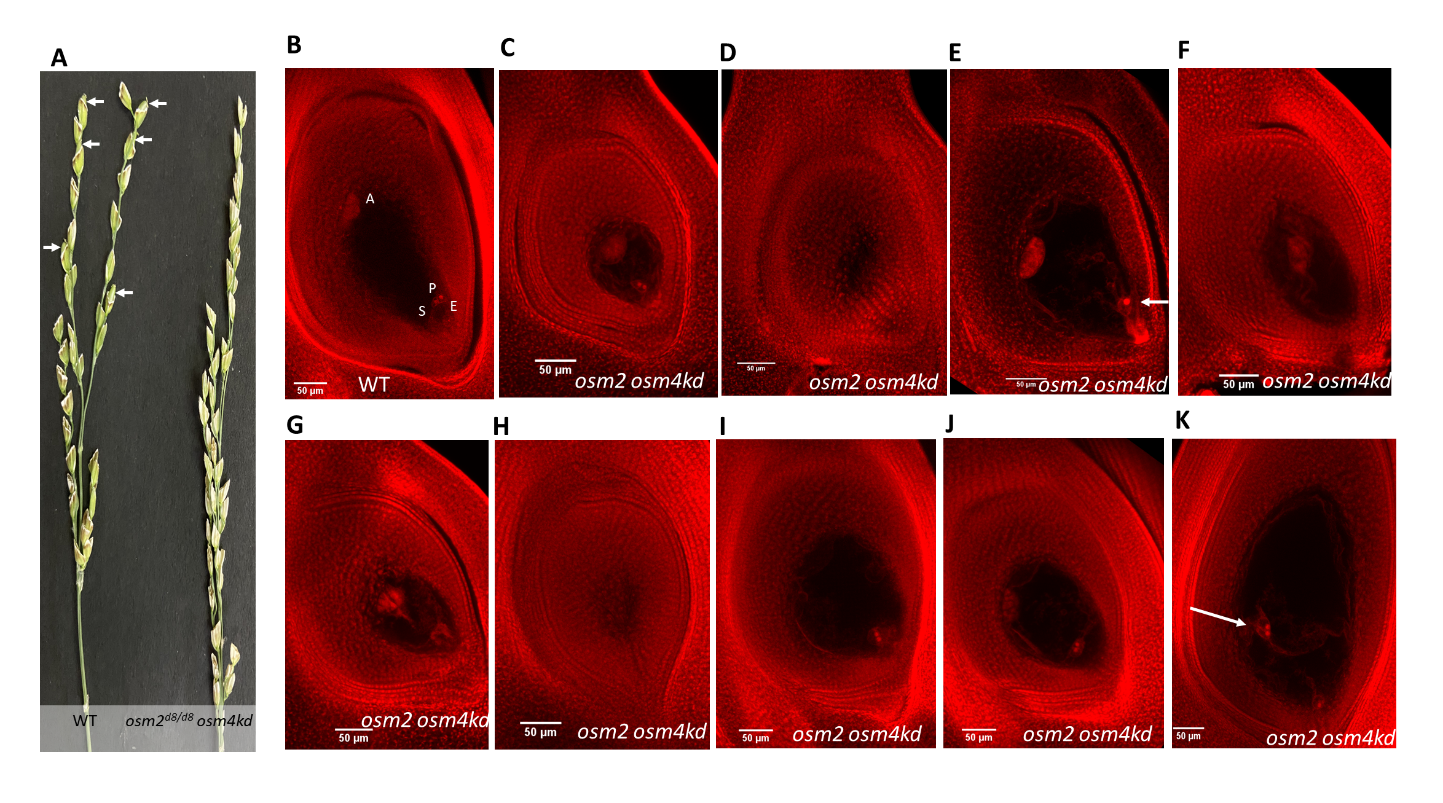
**Figure 2.** **Female sterility in *osmads2^d8/d8^ osmads4kd* double mutants.** A, Emasculated WT and *osmads2 osmads4kd* double mutant inflorescences dusted with pollens from WT, show the development of seed (white arrows) in WT but not in *osmads2 osmads4kd* (*osm2 osm4kd*) double mutant. B-K, Ovaries and ovules from pre-anthesis WT and *osmads2* *osmads4kd* florets. B, Ovule of WT with normal embryo sac constituting A; antipodal cells, P; polar nuclei, E; egg cell, S; Synergid cells. C, Ovule from *osmads2^d8/d8^* *osmads4kd* floret that has apparently normal embryo sac though smaller. D and H, Ovules with degenerated embryo sac (mass of stained cells, no full cavity). Optical sections passing through the center of the embryo sacs for panels D and H are provided in Supplementary Fig. S9, F and G, respectively. E, Embryo sac with single polar nuclei (white arrow). F, Embryo sac with no female unit (no egg cells, no polar nuclei and no synergids). G, Embryo sac with no polar nuclei. I, Embryo sac without antipodal cells. J, Embryo sac with no egg unit (no egg, no synergids). K, Embryo sac with no antipodal cells and with mislocated polar nuclei (white arrow). Scale bars in B-K are 50 µm.

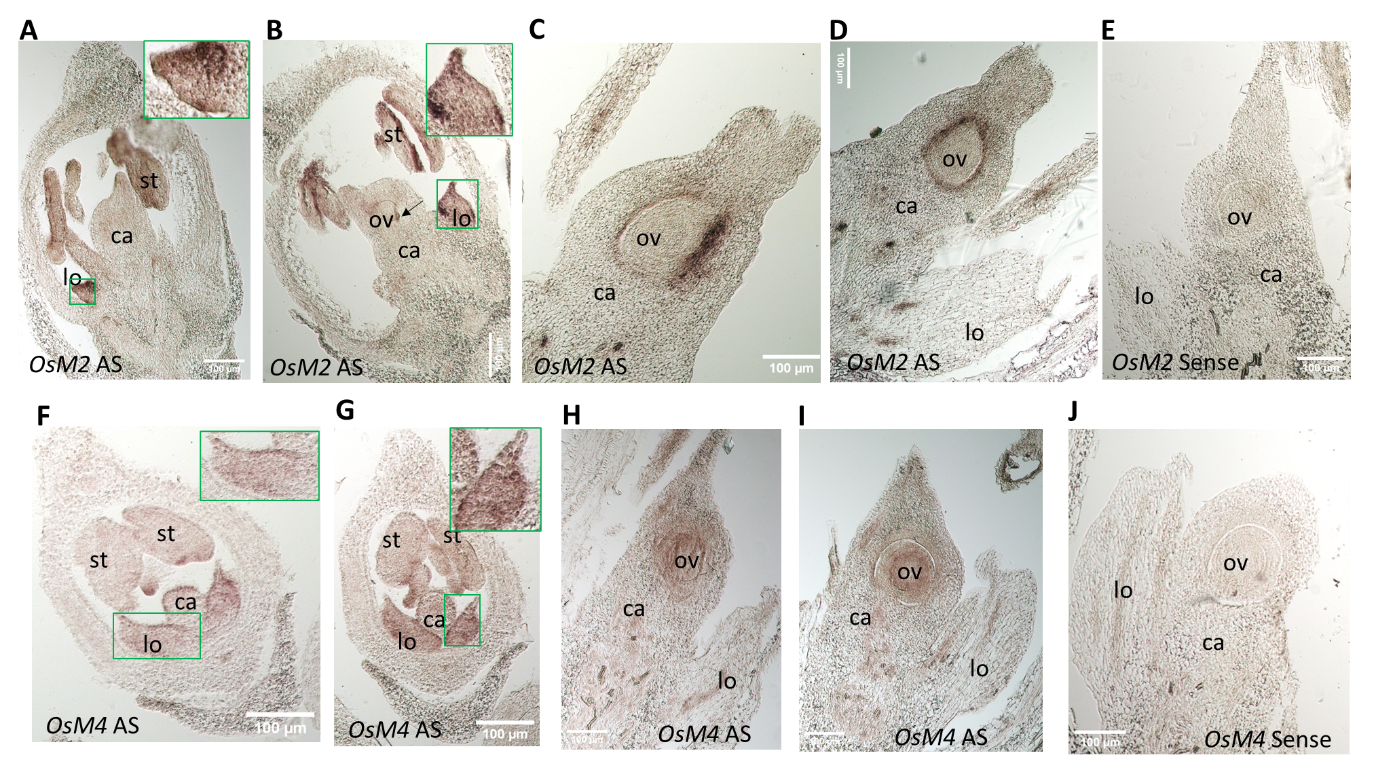
**Figure 3. RNA *in situ* hybridization for *OsMADS2* and *OsMADS4* in developing ovaries with ovules**. A and B, Sections showing signals for *OsMADS2* antisense probe (*OsM2* AS) in lodicules and stamens. The insets show the expression in lodicules which is enriched distally and at the edges. The black arrow in B points at signals in the initiated ovule. C and D, Sections showing *OsM2* AS signals in the inner ovary and outer layers of the developing ovules. E, Section showing no distinct signals for *OsMADS2* sense probe (*OsM2* Sense). F, Sections showing signals for *OsMADS4* antisense probe (*OsM4* AS) in lodicules and stamens. The insets show uniform expression in lodicules. H and I, Sections showing *OsM4* AS signals in the developing ovules. E, Section showing no distinct signals for *OsMADS4* sense probe (*OsM4* Sense). Abbreviations: lo; lodicule, ca, carpel, ov; ovaries. Scale bars are of 100 µm.

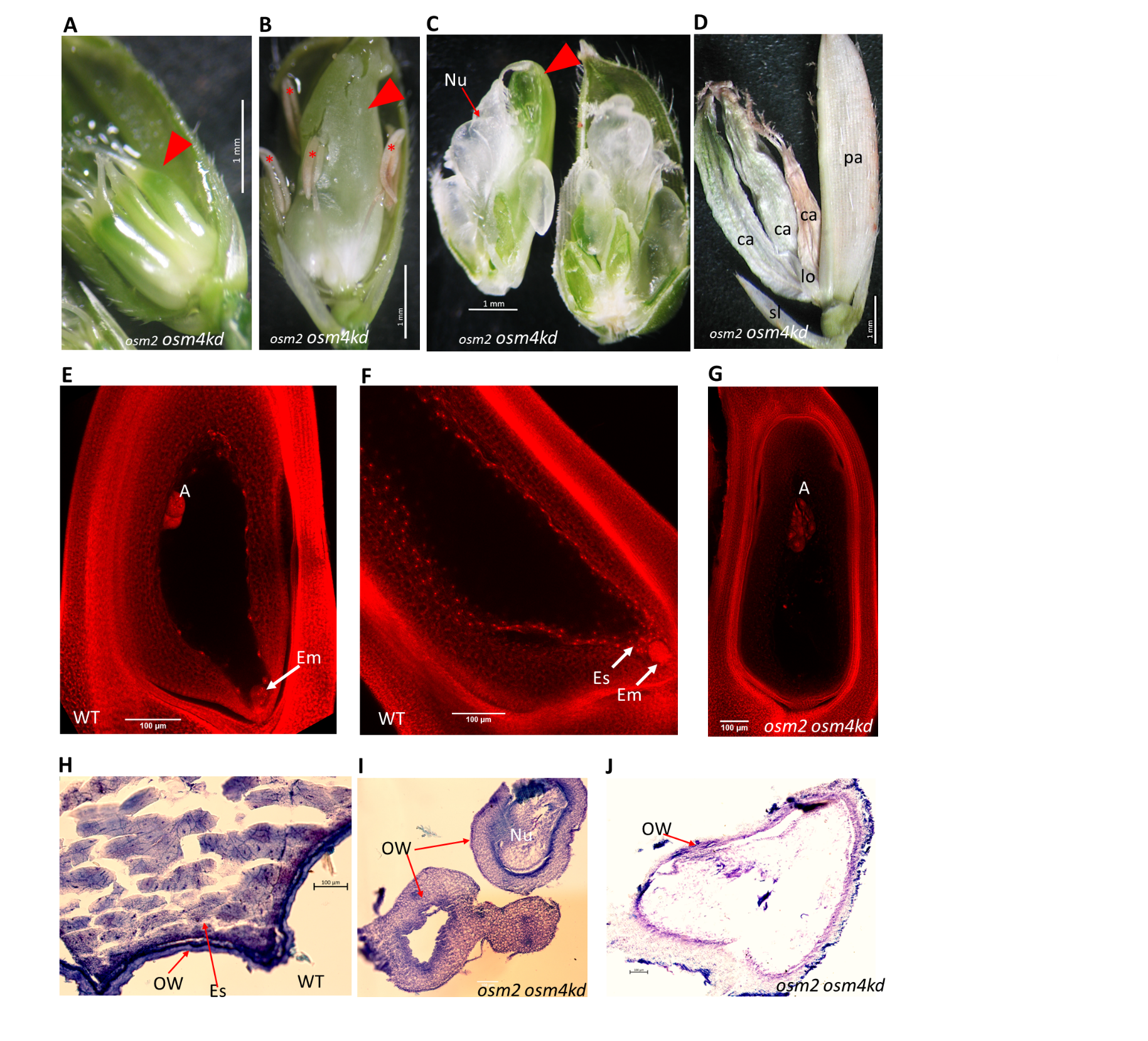
**Figure 4.** **Parthenocarpy-like phenotypes in *osmads2^d8/d8^ osmads4kd* double mutant.** A and B, Pericarpic growth in double mutant florets of type I and type II phenotypes, respectively. The red arrowhead points at the expanded pericarp. In B, asterisks mark the indehiscent empty sterile anthers. The filaments of these sterile anthers failed to extend within the unopened florets. C, Floret of *osmads2^d8/d8^ osmads4kd* double mutant with pericarpic growth. The growing ovary (red arrowhead) was manually dissected to show the over-proliferated nucellar mass inside (red arrow). D, Dry spikelet from *osmads2^d8/d8^ osmads4kd* double mutant where lemma is removed to show the expanded pericarps that are shrunken and dry. E-G, Confocal microscopy Z-stacked images of ovaries from WT (E and F) and *osmads2^d8/d8^ osmads4kd* double mutant (G). The double mutant ovary lacks embryo (Em) and endosperm (Es). H and I, Transverse section of WT and double mutant pericarps, respectively. J, Longitudinal section of double mutant expanded pericarp. Unlike WT, the double mutant pericarp lacks granular endosperm. Abbreviation: Carpel (ca), lodicule (lo), palea (pl), sterile lemma (sl), ovary wall (OW). Scale bars: 1 mm in A-D, 100 µm in E-J.

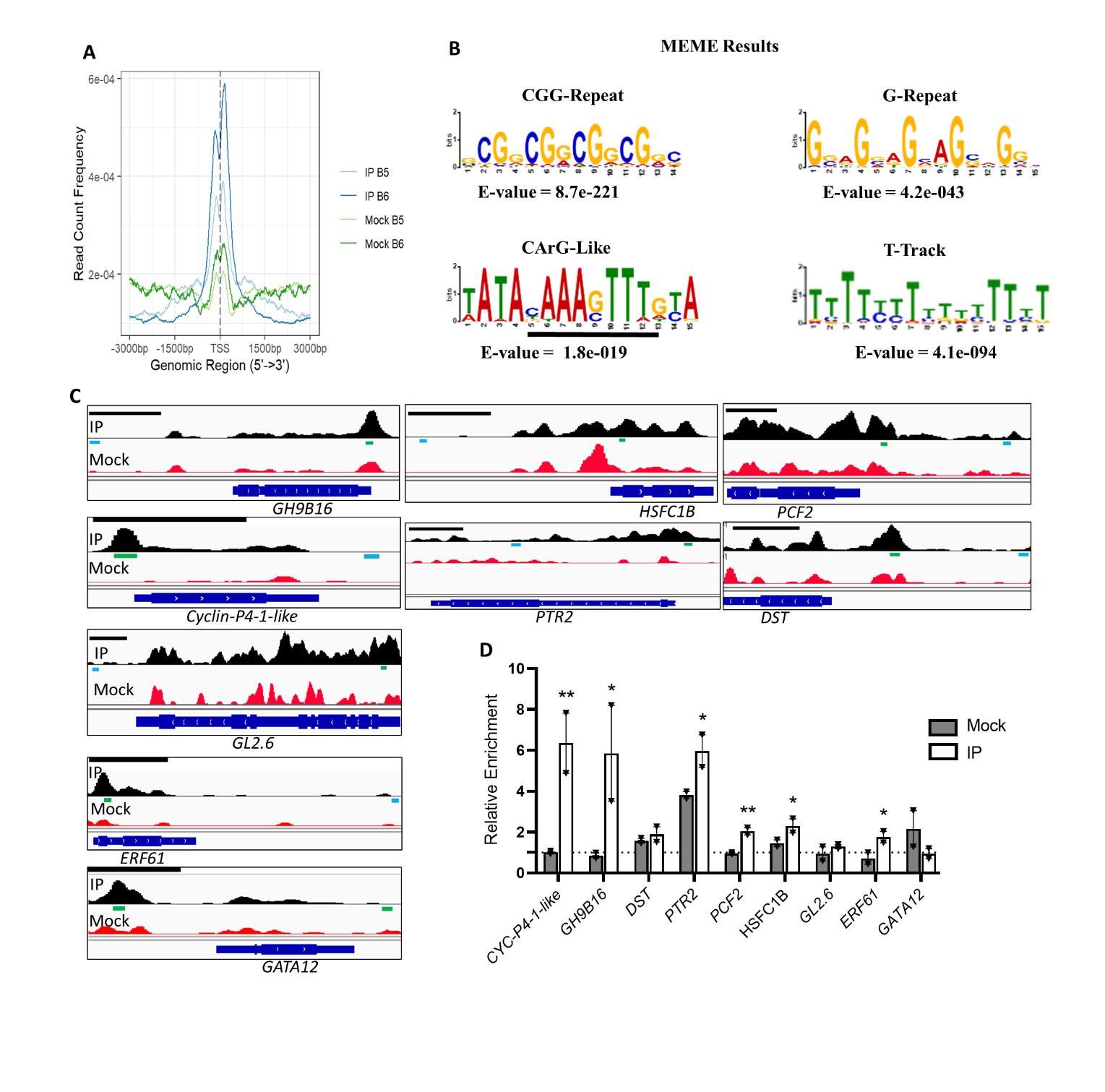
**Figure 5:** **Analysis of genome-wide binding sites of OsMADS2 in WT panicles with developing florets.** A, The plot shows relative distribution of OsMADS2 binding sites (ChIP-Seq peaks) mapped relative to TSS (from 3,000 bp upstream to 3,000 bp downstream). The blue lines represent peaks called by MACS2 algorithm using IP data over mock data. The green lines represent peaks called by MACS2 algorithm using Mock data over IP data. The Peaks of the IP over Mock show enrichment around the TSS. B, Significantly enriched motifs in the peak regions of the global data (MEME analysis). C and D, ChIP-qPCR based validation of some genomic loci bound by OsMADS2 in a pool of 0.5-1 cm inflorescences with developing florets. In C, a snapshot of IGV for each locus is shown where the ChIP-Seq peak regions and nonpeak regions (Control) are represented with green and cyan lines, respectively. The black line above each IGV tab is 1kb of the genome. In D, the bar plot represents the relative enrichment in IP and in mock. The relative enrichment (y-axis) is calculated by dividing the %input of the peak region PCR to the %input of nonpeak control region PCR. The dotted line represents a threshold of 1 on the y-axis which corresponds to the %input values of the nonpeak control region PCR after being divided by themselves. The data are retrieved from two independent biological replicates, each includes 2 technical replicates. Values are means ± SEM. Significance of enrichment in IP over mock tested by multiple unpaired Student’s *t* test. P <0.05 and P <0.001 are denoted as * and **, respectively.

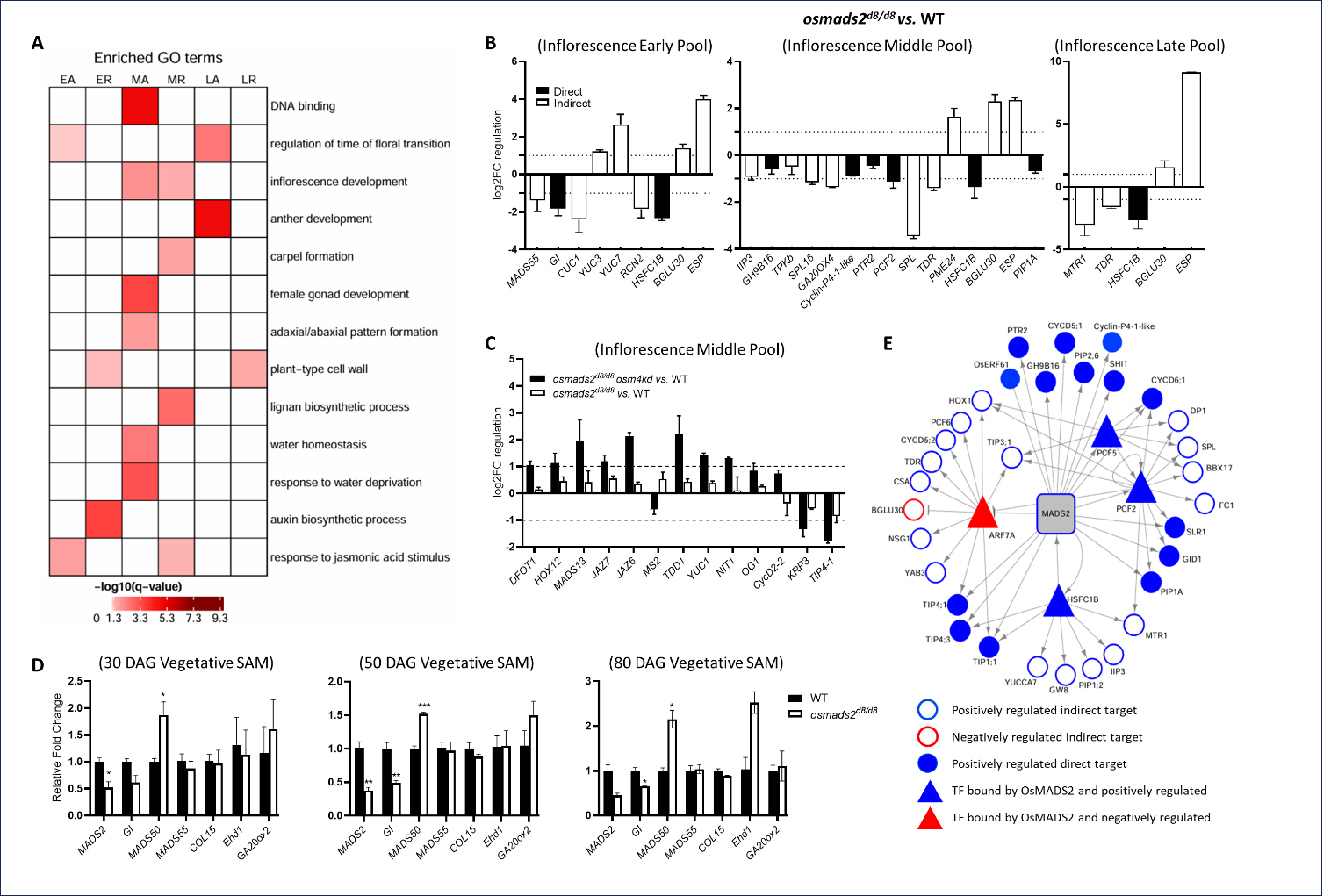
**Figure 6. Gene ontology enrichment analysis and target genes regulated by OsMADS2.** A, Heatmap representation of selected enriched GO terms in the differentially expressed gene sets. Oryzabase database (https://shigen.nig.ac.jp/rice/oryzabase/) GO annotation was used for this analysis and the significantly enriched GO terms were retrieved at qvalue ≤ 0.05. B and C, Plot representations of selected genes that are differentially expressed in panicles of early (0.2-0.5 cm), middle (0.5-1 cm) and late (1-2 cm) pools of *osmads2^d8/d8^*, as per RT-qPCR based validation of the RNA-Seq data (B), and RT-qPCR analysis of candidate genes redundantly regulated by OsMADS2 and OsMADS4 in floral tissue pool constituting 0.5-1 cm inflorescences (C). Values are means ± SEM. The data are from two biological replicates of *osmads2* or two biological replicates of *osamds2^d8/d8^ osmads4kd*, each includes 3 technical replicates. The average log2FC for three biological replicates of WT is equal to 0. The filled bars in panel B represent those genes that are deregulated as per RNA-seq and RT-qPCR data and bound by OsMADS2 as per ChIP-Seq analysis. D, graph plots displaying the relative expression of flowering time regulators interrogated by RT-qPCR in SAM (meristems and leaf primordia). Values are means ± SEM and derived from three biological replicates except for the plot of 80 DAG, the data are derived from two biological replicates. P <0.05, P <0.01d and P <0.001d denoted as *, ** and ***, respectively; Student’s *t* test. E, A small Gene Regulatory Network (GRN) constructed based on ChIP-Seq data and transcriptome (RNA-Seq) data from panicles of 0.5-1 cm (termed here middle developmental pool). The small GRN displays subset of direct and indirect OsMADS2 target genes of annotated functions in osmotic regulation, cell cycle, cell wall modulation, auxin biosynthesis and other relevant to lodicule and stamen development. The full GRN is presented as Supplementary Fig. 14. The filled blue and red nodes represent positively- and negatively regulated direct downstream targets of OsMADS2, respectively. The unfilled circles in blue and red color boundaries represent genes positively and negatively regulated by OsMADS2, respectively. Triangle shape denotes nodal OsMADS2 target genes coding for transcription factors (for e.g. PCF2 and PCF5) that were used to connect these direct target transcription factors to downstream DEGs that form the second layer in this GRN. The arrow shape and the T-shape indicate positive and negative regulation of each OsMADS2 gene target, respectively.

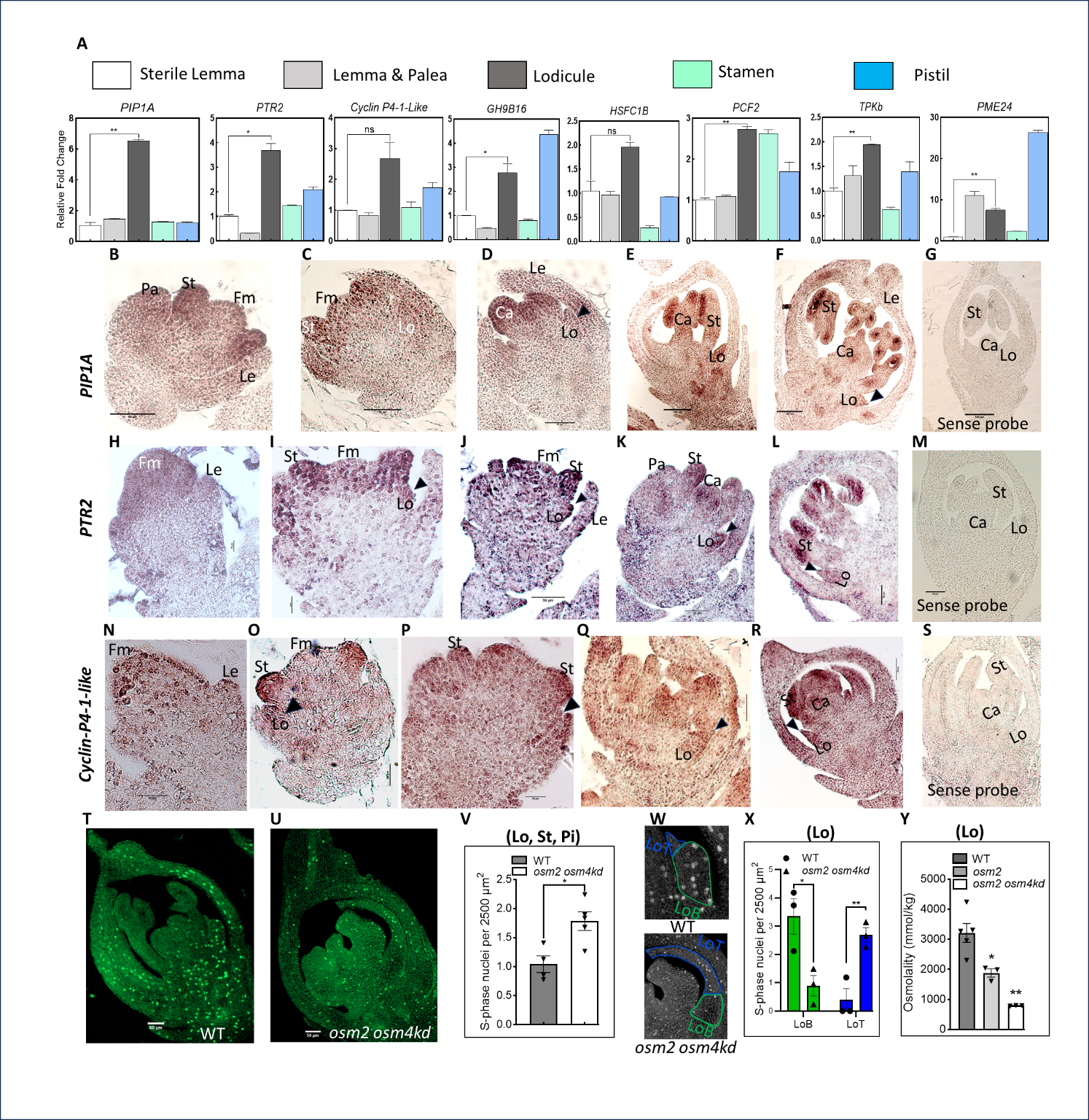
**Figure 7. Spatiotemporal gene expression analysis of OsMADS2 targets and spatial distribution of S-phase cells in the developing floral organs.** A, RT-qPCR analysis of lodicule developmentally relevant OsMADS2 gene targets examining their relative expression levels in the mature floral organs from pre-anthesis WT florets. The relative fold change is with respect to the expression in the sterile lemma. B-G, RNA *in situ* hybridization of *PIP1A* transcripts in developing WT florets. Sections in panels C-F and panel G are hybridized with antisense and sense probes, respectively. H-M, RNA *in situ* hybridization of *PTR2* transcripts in developing WT florets. Sections in panels H-L and panel M are hybridized with antisense and sense probes, respectively. N-S, RNA *in situ* hybridization of *Cyclin-P4-1-Like* transcripts in developing WT florets. Sections in panels N-R and panel S are hybridized with antisense and sense probes, respectively. The black arrowheads, in B-S, point at the lodicules. T and U, representative confocal micrographs of WT and *osmads2^d8/d8^ osmads4kd* florets, respectively. The florets are of equivalent stages as inferred from their roughly equal dimensions. The S-phase nuclei (bright green signals) are labelled with 5-ethynyl-20 -deoxyuridine (EdU). V, Bar graph plotting the number of S-phase cells per unit area, in the inner floral organs. W, confocal micrographs of WT and *osmads2^d8/d8^ osmads4kd* lodicules demonstrating the distribution of S-phase cells in the lodicule body (LoB, green outlines) and lodicule tip (LoT, blue outlines). The whole florets are shown in panel T and U. X, Bar graph plotting the number of S-phase cells per unit area, in the lodicule body and the lodicule tip. Y, Comparison of osmolality levels in the cell sap from WT, *osmads2^d8/d8^* and *osmads2^d8/d8^ osmads4kd* lodicules. Abbreviations: Fm, floral meristem, Le; lemma, Pa; palea, Lo; lodicule, St; stamen, Ca; carpel, Pi, pistil. Values are means ± SEM. P <0.05 and P <0.01d denoted as * and **, respectively; Student’s *t* test. Scale bars: 20 in I, P; 50 µm in A-D, H, J-O, Q-U; 100 µm in E-G.

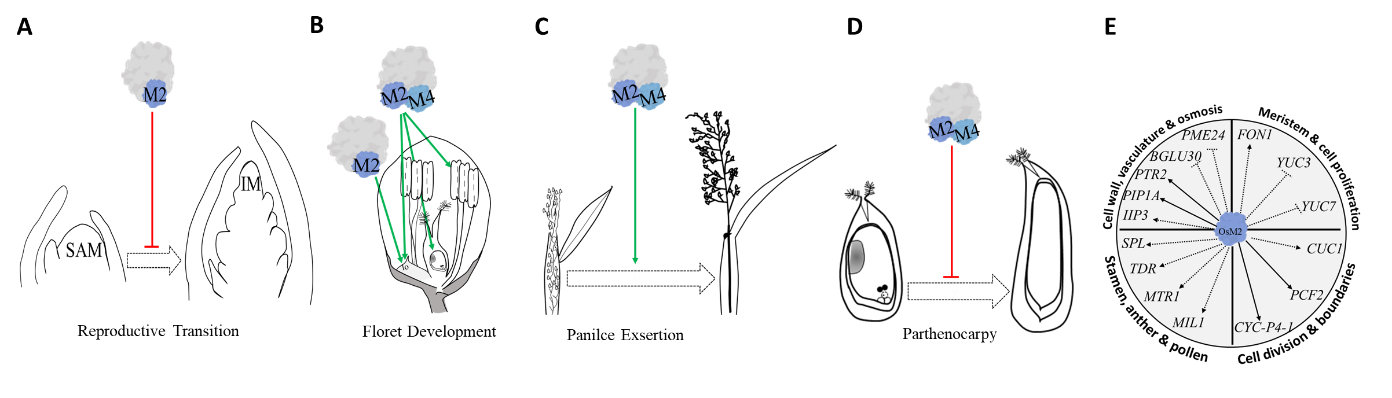
**Figure 8. Graphical summary**. A, OsMADS2 (M2) as part of a complex negatively regulates floral transition. B, OsMADS2 non-redudant role to promote normal lodicule development. Along with OsMADS4(M4) synergistically promotes normal lodicule, stamen and ovule development. C, OsMADS2 and OsMADS4 redundantly promotes inflorescence exsertion. D, OsMADS2 and OsMADS4 redundantly suppresses parthenocarpy. E, OsMADS2 regulates several downstream target genes that are annotated with functions in cell division, meristem size, meristem proliferation, cell wall modulation, vascular development, osmotic regulation, and stamen development. The solid lines and the dotted lines indicate direct and indirect regulation, respectively. The arrow shape and the T-shape indicate positive and negative regulation of each OsMADS2 gene target, respectively.
